# Wearable Focused Ultrasound Neuromodulation and Electrophysiological Recording Patch for REM Sleep Enhancement

**DOI:** 10.1101/2025.10.19.683337

**Authors:** Kai Wing Kevin Tang, Benjamin Baird, William D. Moscoso-Barrera, Mengxia Yu, Mengmeng Yao, Jinmo Jeong, Ilya Pyatnitskiy, Anakaren Romero Lozano, Jiachen Wang, Ju-Chun Hsieh, Tony Chae, Daniel Song, Julieta Garcia, Rithvik Mittapalli, Adam Bush, Wynn Legon, Vincent Mysliwiec, Gregory A. Fonzo, Huiliang Wang

## Abstract

The rise in sleep disease affecting the general population globally in the past decade has been detrimental to individually and socioeconomically. As of now, approaches often are temporary through medication, permanently using invasive implants with surgical complications or neuromodulation therapy. However, non-invasive, state-dependent neuromodulation during sleep is technically challenging, especially with the lack of flexibility, comfortability and robustness for sleep conditions. Here, we introduce a **N**on-invasive **E**lectrophysiological Recording and **U**ltrasound Neuromodulation **Slee**p **P**atch (NEUSLeeP) in delivering focused ultrasound stimulation to the subthalamic nucleus (STN) overnight with simultaneous stable polysomnography recording. Our sleep patch integrates a custom eight-channel concentric ring transducer array with real-time electroencephalography (EEG), electrooculography (EOG), electromyography (EMG) recording, and individualized line-of-sight targeting to sonicate deep brain areas while preserving mobility. Our platform operated safely and comfortably across the two-night’s sleep study. Stimulation of the left STN was delivered every 90 minutes throughout the night and was associated with a 25% increase in REM (Rapid Eye Movement) sleep duration and a 43 minutes reduction in REM sleep latency compared to a sham night in a study of 26 subjects. Blood Oxygen Level Dependent (BOLD) signal attenuation in functional Magnetic Resonance Imaging (fMRI) was localized primarily to a left ipsilateral basal-ganglia-midbrain-temporal circuit, consistent with selective network modulation rather than global arousal changes. Overall, NEUSLeeP demonstrates feasibility by (i) a light weight and wearable ultrasound neuromodulation and sleep recording during natural sleep; (ii) establishing a potential mechanism relating targeted ultrasound stimulation of STN/sleep networks to REM enhancement.

## Main

Sleep is essential for health, performance, and decision-making, but service members often suffer due to demanding schedules, operational commitments. Insufficient sleep impacts their physical and mental performance and long-term health.^1,2^ Sleep is also detrimentally impacted by contribution to symptoms depression, anxiety, and post-traumatic stress disorder (PTSD) symptoms,^3^ conditions which are also prevalent in this population. These sleep challenges extend beyond military service, with a rising trend of 14.5% adults in sleep difficulties amongst the general population^4^. Unfortunately, current sleep-enhancing treatments for enhancing sleep remain limited. Sleep primarily consists of non-rapid eye movement (NREM) and rapid eye movement (REM) stages, each playing a distinct role in physical and mental well-being. With prolonged wakefulness being a widespread phenomenon associated with stress, the impact on cognitive performance is detrimental^5,6^. In particular, recent research highlights the significance of REM sleep in emotional resilience and stress adaptation^7^. These sleep challenges extend beyond demanding professions, with a rising trend of 1/3 of adults having insufficient sleep^8^ and 14.5% having sleep difficulties amongst the general population^4^. Unfortunately, current treatments for enhancing the negative impacts of insufficient sleep, especially REM sleep, remain limited.

In the past two decades, non-invasive sleep stimulation techniques have emerged to enhance sleep using approaches such as transcranial direct current stimulation (tDCS), transcranial magnetic stimulation (TMS), and acoustic stimulation^7,9–11^ As of current, these non-invasive methods lack precision and specificity in non-invasive modulation of deep brain structures, making it difficult to achieve the most effective treatment without causing adverse effects^12,13^. Conversely, deep brain stimulation (DBS) has shown effective promotion of REM) sleep duration specifically at the pedunculopontine nucleus^14,15^ and subthalamic nucleus in Parkinsonian patients^16,17^. However, this requires invasive implanted electrodes deep in the brain. Recent advancements in transcranial-focused ultrasound stimulation (tFUS) show promise for non-invasively stimulating at depth, with the ability to stimulate with high spatial resolution and at deep brain targets in human^18–22^. However, current tFUS systems are bulky and not suitable for overnight sleep intervention, due to continuous human motions during sleep. Recently, our efforts in bioadhesive hydrogel development and ultrasound device miniaturization demonstrated feasibility in long-term and wearable cortical neuromodulation applications^23^. Nevertheless, the development of a sufficiently compact ultrasound transducer enabling adjustable target engagement and necessary acoustic power output for deep brain neuromodulation is still needed for a non-invasive sleep intervention. In conjunction with tFUS, evaluating impact of deep brain neuromodulation on sleep performance necessitates metrics to assess NREM and REM sleep. The current gold standard utilizes electroencephalography (EEG), an effective non-invasive technique commonly used to monitor sleep stages and diagnose sleep disorders and epilepsy^24^. Despite so, current clinical EEG electrodes dry out quickly, limiting their long-term usability. As such, recent work in development of long-term stable and low impedance hydrogels has been realized^25–28^. Yet, it remains a challenge to have a system that enables both FUS neuromodulation for sleep intervention and high-quality, extended electrophysiological recording for sleep staging in naturalistic settings.

Here, we report a fully integrated flexible, bioadhesive and wearable **N**on-invasive **E**lectrophysiological Recording and **U**ltrasound Neuromodulation **Slee**p **P**atch (NEUSLeeP) through targeting of the subthalamic nucleus to enhance sleep^29^ (**Fig. 1a-d**). The NEUSLeeP consists of six electrophysiological recording channels (4 EEG, 1 EOG, 1 EMG) with ultra-low impedance 2-**A**crylamido-2-methylpropane sulfonic acid (AMPS) based **S**leep **G**el (ASG) to provide sufficient coverage to capture key features in sleep recording. In addition, the device includes a custom-made **C**oncentric **R**ing **U**ltrasound **T**ransducer **A**rray (CRUTA) allowing adjustable axial focal depth for FUS to specifically target the subthalamic nucleus (STN) for neuromodulation (**Fig. 1b**). Finally, our NEUSLeeP integrates the CRUTA and electrophysiological recording units with our custom-designed long-term stable bioadhesive elastomeric substrate using **E**coflex-**Gel** modified with **P**oly**e**thylene**i**mine **e**thoxylate (Eco-PEIE-Gel). Overall, the NEUSLeeP device weighs ∼103.4 grams and easily attaches to various skin surfaces including oily and hairy skins for extended duration, allowing ease of use for overnight sleep applications. To demonstrate the application of NEUSleeP for tFUS stimulation and sleep recording, we used NEUSLeeP to monitor EEG sleep metrics and deliver tFUS stimulation to the left STN during sleep, with the goal of improving REM sleep performance with accurate sleep staging. Furthermore, we also characterized the effects of tFUS on physiological metrics of stress management and brain activity modulation, using heart rate variability measurements and functional magnetic resonance imaging (fMRI) both at rest and during the processing of emotional stimuli (**Fig. 1c**).

**Figure 1.**
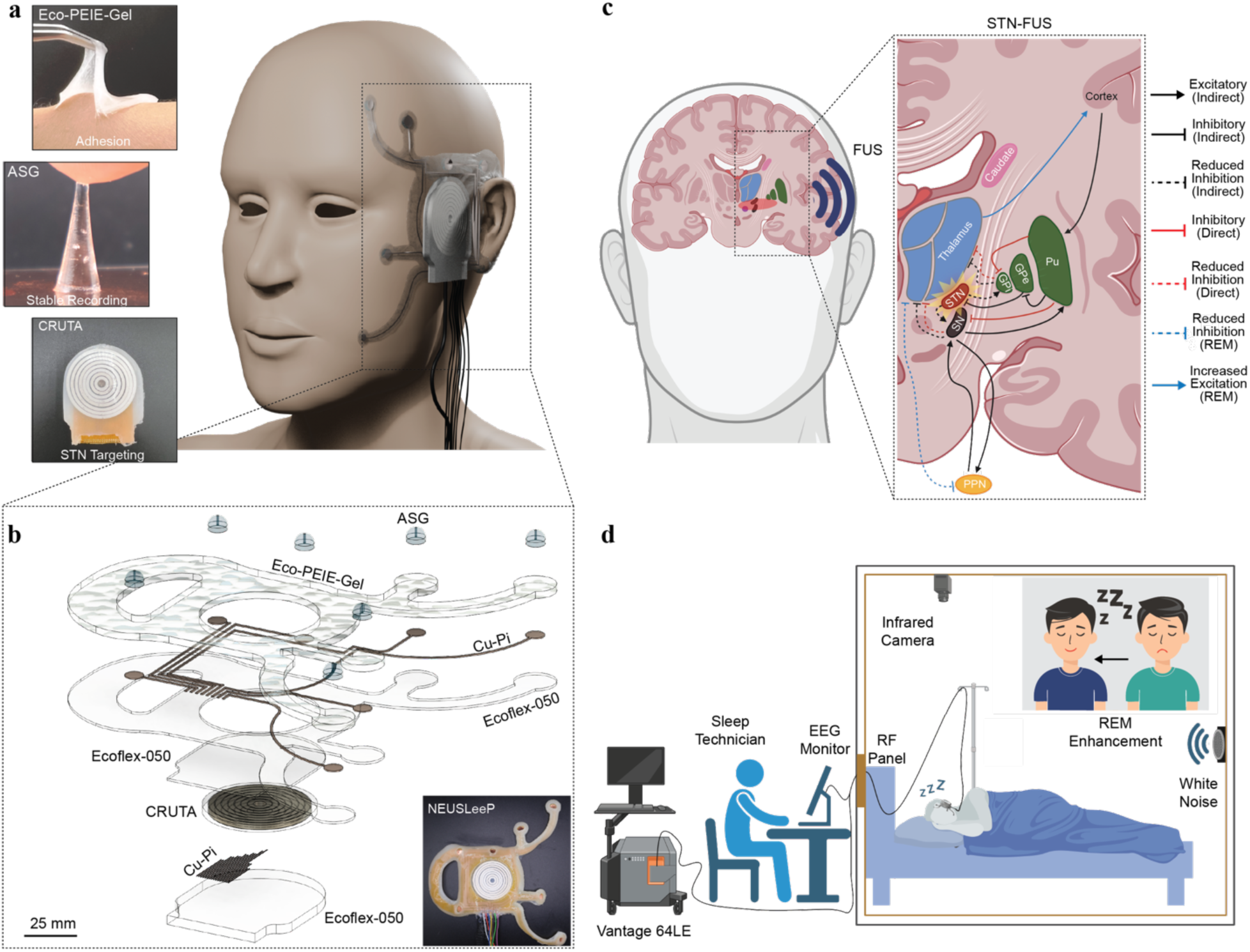
Overview of NEUSLeeP. **a)** Photograph of wearable NEUSLeeP comprised of Eco-PEIE-Gel substrate enabling stable and high adhesion to the skin, ASG for long-term low-impedance stable sleep recording, and CRUTA for focus-adjustable focused ultrasound stimulation targeting of the STN. Photo used with permission by the first author; written informed consent for publication was obtained. **b)** Design of the NEUSLeeP. The device implements a multi-layer fabrication approach consisting of an eight-channel CRUTA, transfer-printing of laser-etched copper interconnects on polyimide (Cu-Pi), mounting of conductive hydrogel (ASG), and encapsulation with bioadhesive elastomer (Eco-PEIE-Gel). The exploded view of the device shows two parts: (1) the FUS layer consisting of the CRUTA, and (2) the electrophysiological recording layer consisting of the Eco-PEIE-Gel and ASG. The two layers are bonded and could be separated allowing modular design. **c)** Proposed mechanism of STN-FUS towards modulating the downstream effects of the basal ganglia network to improve REM sleep performance through direct/indirect modulation of pedunculopontine nucleus (PPN). **d**) Schematic of sleep study with NEUSLeeP to induce improvements in REM sleep. Created in BioRender. Tang, K. (2025) https://BioRender.com/xi5ugz1

## Results

### Design of adhesive Eco-PEIE-Gel for stable interfacing with skin

The current state of non-invasive stimulation including FUS lacks wearability and ease of use due to the bulkiness and weight^30^. Whilst advancements in body-comfortable wearable electronics approaches these obstacles through improvement of device thinness and compliance to enhance interfacial adhesion^31^, this method is primarily applicable for sensor technologies as FUS deep brain stimulation modalities require high power deposition resulting in the need for a bulky and heavy device. Alternatively, bioadhesives presents numerous advantages in wearable device application due to its’ ability to adhere to the bio-tissues through either covalent bonds, non-covalent bonds, mechanical interlocking, and/or structural interfacial linking^32^. As such, a rise in development of bioadhesive hydrogel laid the foundations in bio-compatible wearable applications^33^. However, a major disadvantage in hydrogel is its dependency in water-swollen polymer network formation. This results in its susceptibility of either dehydration over time or swelling when exposed to humid environments. Furthermore, hydrogels in previous work yield insufficient adhesiveness to withstand the weight of piezoelectric transducers and are inadequate for encapsulation due to its’ conductive properties yielding shorting of electrodes and interconnects of the device. Therefore, the development of a robust and non-conductive material with strong adhesion to various skin conditions is necessary for our NEUSLeeP.

Silicon-based elastomers such as Ecoflex and polydimethylsiloxane (PDMS) has been used vastly in flexible wearable electronics in the past two decades. However, their innate adhesion strength to skin is extremely low due to the interfacial Van der Waal’s force^34^. Furthermore, when comparing Ecoflex and PDMS, Ecoflex offers a much flexible and soft mechanical property due its’ low Young’s modulus. Previous work in utilizing polyethyleneimine ethoxylated solution (PEIE) mixed with PDMS enabled a homogenous cross-linked network with skin adhesion of ∼0.48 N/cm and Young’s modulus of 24 kPa^35^. Here, we chose to utilize Ecoflex-Gel as the backbone of our bioadhesive layer modified with PEIE to create a robust and soft bioadhesive elastomer (Eco-PEIE-Gel). Essentially, the addition of PEIE creates a polymer chain of amine, hydroxyl, and carboxylic groups that are important for creating hydrogen bonding, electrostatic ion pairing and ammonium-carboxylic pairing with the stratum corneum (**Fig. 2a-b**). The surface modification of PEIE also creates small pores, potentially supplementing mechanical interlocking structures aside from the miniscule Van der Waal’s force. Evaluation of the chemical structures with PEIE addition at various weight loading using Attenuated Total Reflectance with Fourier Transform Infrared spectroscopy (ATR-FTIR) indicates the addition of N-H and O-H groups, validating the proposed mechanism of improved interfacial adhesion of skin (**Fig. 2c**). To demonstrate universal adhesiveness, Eco-PEIE-Gel substrate were prepared with 0.95:0.05 weight ratio of Ecoflex-Gel to PEIE and applied to metal (Al, Cu, Fe), thermoplastic polymers (PLA, PP), elastomers (Ecoflex-050, PDMS), and skin. Overall adhesion strength remains > 0.6 N/cm except for Ecoflex-050 and silicon (**Fig. 2d**). More importantly, adhesion strength with skin (0.749 ± 0.177 N/cm) provides a 2,024% increase compared to Ecoflex-Gel alone (0.037 N/cm)^36^. Adhesion cycling on skin was then performed at 5% and 10% weight loading for up to 20 cycles demonstrating stable and sustained adhesion (**Fig. 2e**). Moreover, the increased PEIE loading improves the stretchability of the Eco-PEIE-Gel whilst maintaining high flexibility (E = 18.11 ± 0.07 kPa at 5% wt), resembling a 25% increase in flexibility compared to previous work with PDMS (**Fig. 2f-h**). Overall, Eco-PEIE-Gel provides a significant improvement in bioadhesives with robust and stable interfacial adhesion, suitable for long-term sleep recording and neuromodulation with relatively heavy transducers.

**Figure 2.**
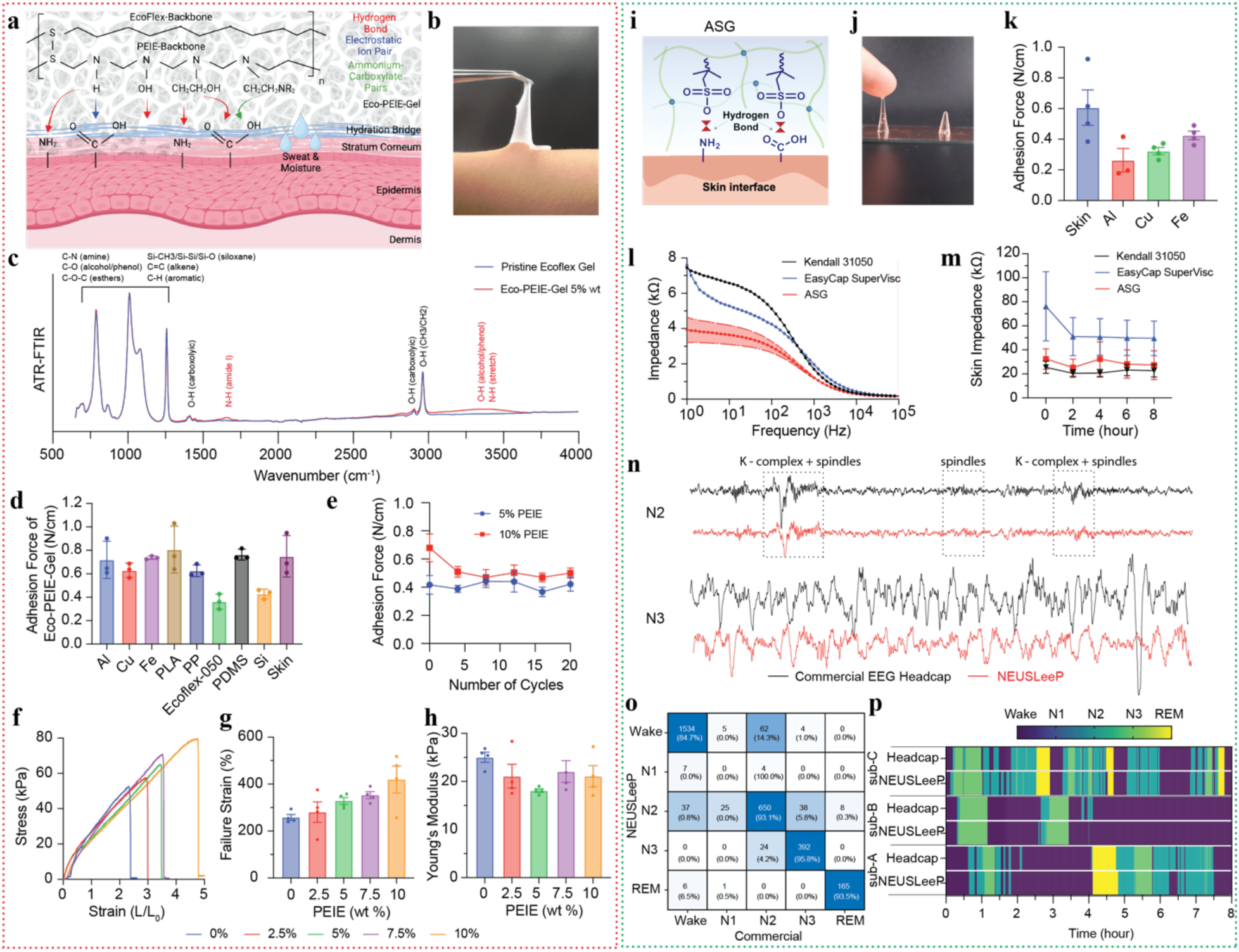
Materials characterization of Eco-PEIE-Gel bioadhesive encapsulation layer and ASG recording electrodes. **a)** Schematic of chemical structure of Eco-PEIE-Gel and its mechanism to interface with the human skin through hydrogen bonding, electrostatic pairing, and ammonium-carboxylic pairing of the stratum corneum in strong adhesion. **b)** Photograph of Eco-PEIE-Gel on the skin. **c)** ATR-FTIR of 5% weight percentage of PEIE in Eco-PEIE-Gel and Pristine Ecoflex gel, outlining the modification of chemical structure in Eco-PEIE-Gel. **d)** 90° T-peel test of Eco-PEIE-Gel (PEIE 5 wt%) on various materials to determine its adhesion strength (n = 4, independent samples). **e)** Adhesion cycling of Eco-PEIE-Gel (5 wt% and 10 wt%) on human skin (n = 4, independent samples). **f)** Stress-strain curve on various loading of PEIE in Eco-PEIE-Gel (n = 4). **g)** Failure strain of Eco-PEIE-Gel of varying PEIE loading (n = 4, independent samples). **h)** Young’s modulus derived from stress-strain curve of varying PEIE loading (n = 4, independent samples). **i)** Schematic of chemical structure of ASG and its’ adhesion allows strong interfacing with the skin. **j)** Photograph of ASG on skin and copper. **k)** 90° T-peel test of ASG on skin and metal substrates (n = 4). **l)** Bode impedance comparison of ASG to commercial-grade electrodes (Kendall 31050) and conductive gels (EasyCap SuperVisc). **m)** Long-term skin impedance over 8-hours comparing ASG to commercial-grade electrodes and gels (n = 4, independent samples). **n)** Comparison of a 30-second window in recorded signal during sleep at N2 and N3 between NEUSLeeP (with ASG) and commercial EEG headcap (32-Channel AntNeuro). **o**) Correlation matrix of hypnogram between NEUSLeeP and commercial EEG headcap (32-channel AntNeuro). **p)** Hypnogram of overnight sleep recording (n = 3 for each recording, Commercial vs. NEUSLeeP). All plots show mean ± s.e.m unless otherwise mentioned, *P < 0.05, **P < 0.01, and ***P < 0.001. All data generated in this figure are provided in the Source Data file. Created in BioRender. Tang, K. (2025) https://BioRender.com/xi5ugz1

### Design of ASG for high-qualify electrophysiological recording during sleep

Previous work in the development of a self-adhesive and stable hydrogel for sleep recording using 2-acrylamido-2-methylpropane sulfonic acid (AMPS) with poly(3,4-ethylenedioxythiophene) (PEDOT:PSS) enables a high ionic conductivity and water absorption capability^27^. Similarly, sodium-based with PEDOT:PSS hydrogels enabled similar efficacy for sleep recording^37^. Despite exemplary performance, the need for PEDOT: PSS with residues on the scalp often leads to many inconveniences and discomfort for long-term wearability with potential contamination of the device visually. In previous work, a bioadhesive acoustic hydrogel utilizing AMPS-based network provided a long-term stable adhesion to the scalp for up to 28 days^23^. Therefore, the AMPS-based Sleep Gel (ASG) was developed by leveraging the poly(AMPS) chain network with water-glycerol solvent initiated with ammonium persulfate (APS) and catalyzed with a thermo-initiator (Tetramethylethylenediamine, TEMED) cross-linked with methylenebisacrylamide (MBAA). This creates an extremely soft, clear, and adhesive hydrogel to the skin (**Fig. 2i-k**), with the on the skin adhesion of (0.606 ± 0.231 N/cm) similar to the Eco-PEIE-Gel. Whilst the adhesion may not be comparable to that of our previous work^27^, we elected to focus on the low impedance of hydrogel to improve signal quality of sleep recording. This is because the adhesion of Eco-PEIE-Gel in the large area of NEUSLeeP provides sufficient support to the skin for our system (**Fig. 1a**). By removing PEDOT: PSS; which operates via conjugated-π-electron networks and instead using solely AMPS as the primary ionic monomer which is suitable for skin interfaces; allows for excellent biocompatibility and improved conductivity with the presence of salt ions in sweat. We then increased glycerol content to improve hydration retention and utilized thermal initiator (MBAA). This effectively mitigates the inhomogeneous curing of the hydrogel using photo-initiators often due to the impermeable light of PEDOT: PSS. Additionally, thermal heating creates a drier surface with improved adhesion whilst maintaining high water content internally resulting in the improved low impedance. As a result, the trade-off in adhesion enables a ∼50% decrease in impedance within the critical EEG signal band range of 1–100 Hz, performing favorably compared to commercial-grade electrodes and conductive gels (**Fig. 2l-m**)^38^. To ensure the impedance remains stable and similar during long-term recording, a comparison of ASG and commercial electrodes/gels were applied on to the skin for up to 8-hours and measured. Here, the ASG performs similar to the commercial-grade electrodes with great stability. To further demonstrate the feasibility of a stable sleep recording with ASG, the hydrogel was integrated into the NEUSLeeP. Three subjects then wore both the NEUSLeeP and commercial EEG headcap (AntNeuro 32-channel EEG headcap) simultaneously. Results indicate virtually no difference in terms of signal quality and sleep staging (**Fig. 2n-p, Supplementary Fig. 1**). The development of ASG provides a soft, comfortable, and low impedance electrode that enables sleep recording, which would be essential for the purpose of electrophysiological recording in NEUSLeeP.

### Design and Characterization of CRUTA with adjustable focal depth

The 8-channel piezoelectric CRUTA leverages the high electromechanical coupling of bulk piezoelectric to generate high acoustic pressures at a specific resonant frequency (high Q-factor) and hybridizes it with soft elastomer encapsulation to enable a light-weight, comfortable, and acoustic coupling with soft biological tissue^39,40^. The combination of design and flexible housing enables the CRUTA to weigh less than 103.4 g and 3.5-mm thick, approximately 80% thinner than existing ultrasound stimulation devices^41,42^. The CRUTA with a thickness of 3.1-mm has a center frequency of 0.65 MHz, where successful neuromodulation was reported in cortex^23,43^, thalamus^44^, and amygdala^45^ This enables a millimeter axial resolution (**Fig. 3a-d, Supplementary Fig. 2**) by controlling the phase delay between each channel element, comparable with currently available ultrasound stimulation devices operating at similar frequencies (**Supplementary Table. 1**). Transfer printing of a laser-etched bilayer stacking of polyimide (PI, 24 μm, Cu 660 µm) was used to fabricate flexible electrodes to interconnect between the eight concentric ring arrays of transducers and LEMO-00 wires (see **Methods**). The array was designed to target the STN through the temporal skull above the zygomatic arch, with coverage of varying anatomical positions across the general population in STN (72.2 ± 4.62 mm, **Supplementary Fig. 3**) from the temporal window was verified. Acoustic pressure of the CRUTA was adjustable through changes in voltage driven from the Vantage 64LE system (**Fig. 3d**., **Supplementary Fig. 2**). Variances in device performances were accounted for by characterization of focality, acoustic pressure, and impedance.

**Figure 3.**
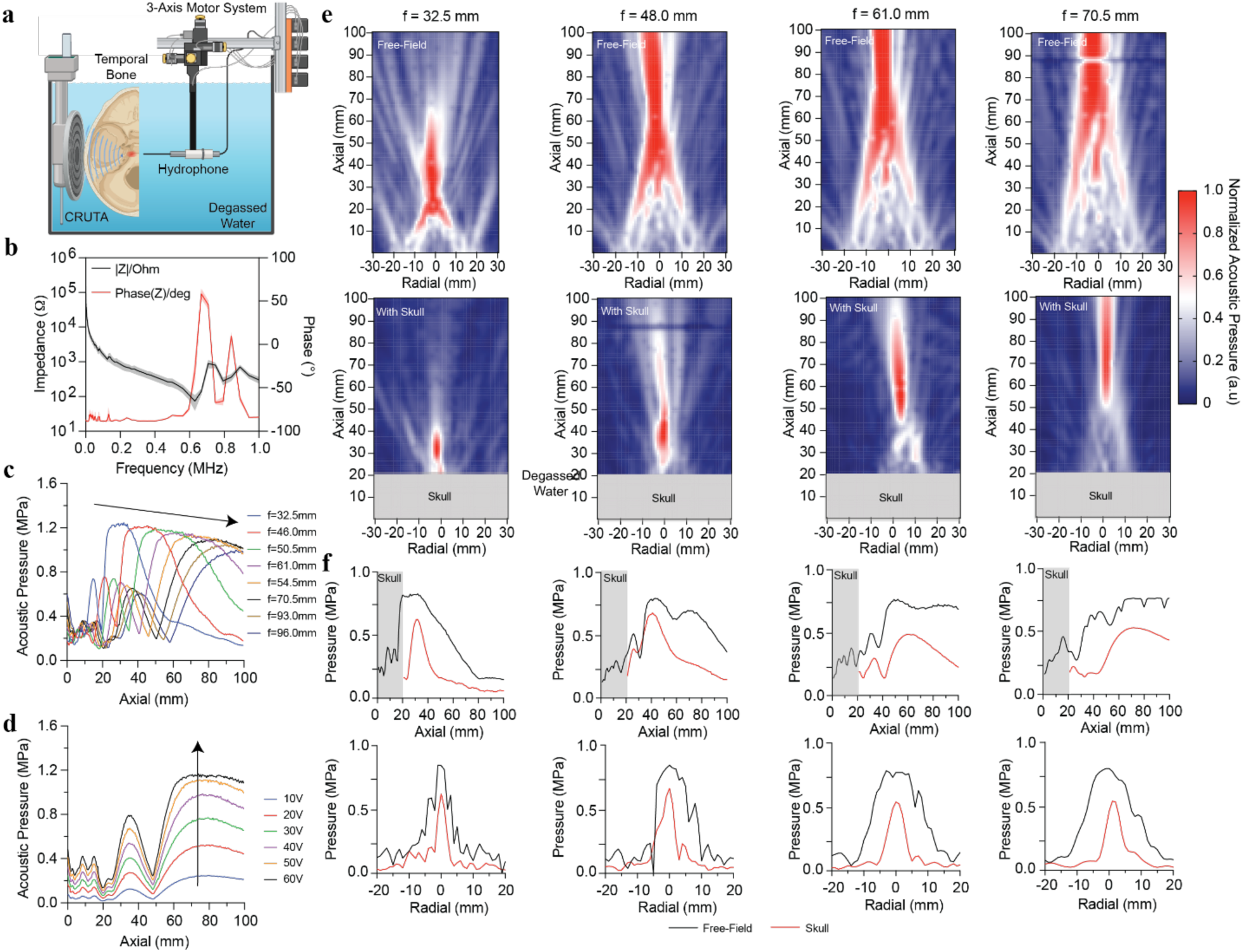
Concentric Ring Ultrasound Transducer Array (CRUTA). **a)** Experimental setup of CRUTA characterization. **b)** Bode impedance and phase response of CRUTA (n = 8). **c)** Axial acoustic pressure of CRUTA with increasing focal depth adjustments. **d)** Axial acoustic pressure of CRUTA with increasing driving voltage at a fixed focal depth. **e)** Acoustic field distribution of CRUTA with and without the temporal bone at varying focal depths (f = 32.5, 48.0, 61.0, and 70.5 mm) normalized to the peak pressure of each focal depth. **f**) Axial and radial acoustic profiles measured and the corresponding focal depths with and without skulls demonstrating the effects of attenuation due to the temporal skull.

In addition, the characterization of acoustic pressure fields emitted from the CRUTA was done by using a calibrated hydrophone on a motorized 3-axis system submerged in a degassed distilled water tank (**Fig. 3a**). Firstly, measured acoustic field distribution in free-field was performed, where the full-width half maximum of axial and radial beam profile increases exponentially (**Supplementary Fig. 3**). At the desired STN target of 72.2 mm, the axial/radial FWHM in free-field and skull were 44.6 mm/58.6 mm and 18.6/5.2 mm respectively. The focality of the beam profile decreases as focal depth increases due to the constraints of the geometric transducer diameter (47.57 mm) and center frequency due to the principle of interference in acoustics^46^. The complexity of the effects in skull altering beam properties are dependent on the morphology, thickness, and angulation of the skull^47^. As such, other works using computational models of transcranial focused ultrasound, constructed from CT and MR scans in varying sexes and skull segments, have demonstrated a spatial focusing effect on the beam profile through the temporal window^48^. As such, improvement in FWHM was observed when the temporal skull was present both axially and radially with the CRUTA (**Fig. 3e-f**, **Supplementary Fig. 4**).

### STN-FUS results in ipsilateral changes of the basal ganglia network

Deep brain stimulation at the STN (STN-DBS) for Parkinson’s disease not only improves motor symptoms but also improves sleep quality and reduced sleepiness^49,50^. Specifically, STN-DBS has been shown to increase REM duration percentage significantly from +1.3%^51^ to 6.2%^16^. Yet, the need for an invasive surgical implant often leads to long-term complications including cognitive,^52^, non-motor^53^ decline, and glial response induced inflammation resulting in reduced efficacy^54^. The proposition of STN-FUS provides a non-invasive alternative to achieve the same outcomes of STN-DBS. However, the fundamental mechanisms of STN-FUS regarding neuromodulation are still unclear due to differences in mechanism to DBS^55^. Nevertheless, the STN has been demonstrated to play a vital role in inhibitory control of movement and attentional processes^56^. Furthermore, inhibition of the STN may promote muscle atonia and reduced wakefulness^57–60^. Here, we select a unilateral approach in stimulating the left STN as previous works suggests significant improvements in sleep disturbances were observed.^29^ As such, for the first time, we utilize FUS to target left STN with CRUTA.

16 individuals were assessed, recruited, and enrolled (**Supplementary Fig. 3** and **Supplementary Table. 2**). The sample encompassed a wide spectrum of age (25.7 ± 6.28 years) and assessed through the Pittsburgh Sleep Quality Index (PSQI). An initial pilot-study test with STN-FUS with fMRI-compatible BrainSonix was performed and indicated a bi-modal effect at the STN dependent on the pulse repetition frequency (PRF) when duty cycle (5%) and a derated spatial peak temporal average intensity (∼719 mw/cm^2^) were held constant, where 10 Hz induced excitatory and 100 Hz induced inhibitory effects^41^ (**Supplementary Fig. 5**). The effects induced inhibition observed in higher frequency stimulation align with our results, where high frequencies^61,62^ (> 100 Hz) improve symptoms with broadly attenuated beta-band power and low frequencies (< 60Hz) amplified alpha/low-beta power local field potentials in the STN^63,64^. As such, 100 Hz PRF was used throughout all this study by CRUTA to promote inhibition of STN. Each individual was subjected to a session consisting of an anatomical scan followed by a pre-FUS resting state functional scan (Pre-Evaluation) to establish baseline and targeting of the STN (**Fig. 4a-b**) before being subjected to 10 blocks of FUS (**Table. 1**, Pressure: 0.90 MPa, 30s ON 30s OFF, Pulse Duration = 0.5 ms, Pulse Repetition Frequency = 100 Hz), followed by a post-FUS resting state functional scan for comparison. The threshold of derated acoustic power taking into account the temporal bone (>0.70 MPa) was selected to ensure biosafety but also sufficient intensity necessary^18,45,65,66^. Blood oxygen saturation levels, heart rate, and blood pressure were measured throughout the session, showing no significant changes with STN-FUS by CRUTA on health vitals (**Fig. 4c-d**).

**Figure 4.**
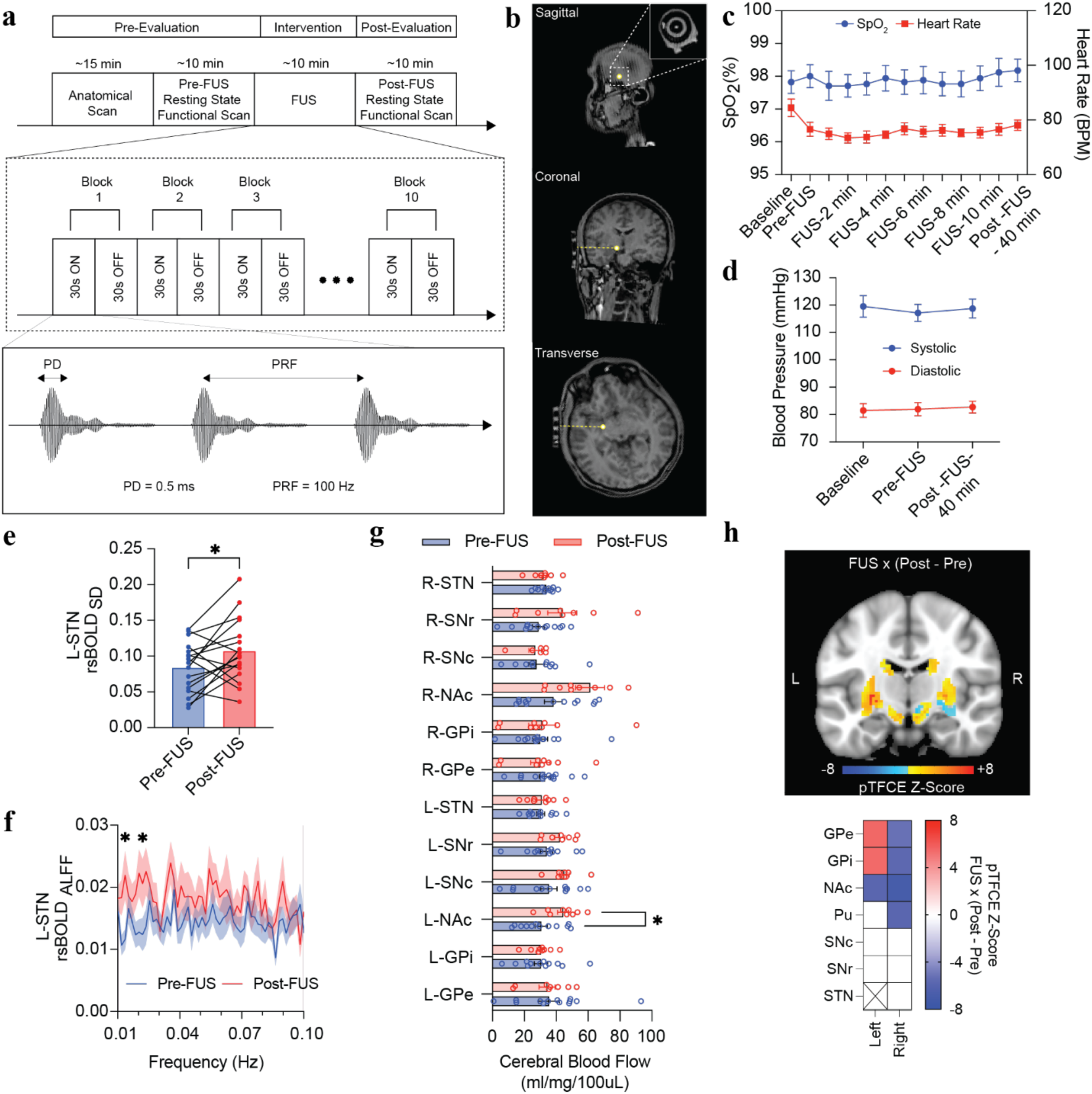
STN-FUS with CRUTA elicits immediate changes in the local basal ganglia network. **a)** Experimental protocol for evaluating the effects of STN-FUS with CRUTA by performing resting state functional scan before and after stimulation. 10 min of stimulation (10 blocks, each block 30 s ON and 30 s OFF, PRF 100 Hz, PD = 0.5 ms, 0.90 MPa). **b)** Photograph of MRI-guided positioning and targeting of CRUTA for STN-FUS. **c)** Blood oxygen saturation levels (SpO_2_) and heart rate were monitored throughout the session showing no significant changes with STN-FUS. **d)** Measured blood pressure indicates minimum changes with STN-FUS. **e)** Resting state blood oxygen level dependent variability (rsBOLD_SD_) at the left subthalamic nucleus (L-STN) increases with STN-FUS compared to baseline. **f)** Amplitude of low frequency fluctuation (rsBOLD_ALFF_) is significantly elevated below 0.03 Hz (n = 16, independent samples, Two-sided paired t-test). **g)** Cerebral blood flow (CBF) obtained from pseudo-continuous arterial spin labeling (pCASL) masked at regions-of-interest. Bilateral nucleus accumbens (NAc) indicates elevated CBF (n = 16, independent samples, Two-sided paired t-test). **h)** Group-level analysis to compare effects of left STN-FUS on rsBOLD before and after stimulation. Significant change in left GPe, GPi, NAc, and Pu were observed (n = 16, linear mixed effect model with probabilistic threshold-free cluster enhancement). pTFCE values were zeroed between -4.90 and 4.90 for statistical representation. All plots show mean ± s.e.m unless otherwise mentioned, *P < 0.05, **P < 0.01, ***P < 0.001, and *pTFCE > |4.90|. All data generated in this figure are provided in the Source Data file.

**Table. 1.** Summary of stimulation parameter for NEUSLeeP-enabled STN-FUS.

| Parameter | Free-Field | In-Situ<br>(Derated at focal depth of 72.2 mm) | FDA<br>Guideline |
| --- | --- | --- | --- |
| Center frequency ($f_0$) | 650 kHz | - | - |
| Pulse Duration (PD) | 0.5 ms | - | - |
| Pulse Repetition Frequency (PRF) | 100 Hz | - | - |
| Pulse Duty | 5% | - | - |
| Burst Duty | 50% | - | - |
| Cycles per pulse | 325 | - | - |
| Cycles per burst | 975,000 | - | - |
| Peak Pressure (P) | 0.90 MPa | 0.68 MPa | - |
| Spatial peak pulse average intensity ($I_{\text{sppa}}$) | 26.9 W/cm<sup>2</sup> | 15.4 W/cm<sup>2</sup> | - |
| Spatial peak temporal average intensity ($I_{\text{spta}}$) | 0.674 W/cm<sup>2</sup> | 0.385 W/cm<sup>2</sup> | <0.720 W/cm<sup>2</sup> |
| Mechanical Index (MI) | 0.93 | 0.70 | <1.90 |

Blood oxygen level-dependent (BOLD) time-series extracted from left STN through region-of-interest (ROI) constrained analyses (see **Methods**) revealed a significant increase in resting-state BOLD variability (rsBOLD_SD_). rsBOLD_SD_ serves as a biomarker to cognitive states correlating to brain glucose metabolism and inversely correlated to task activations^67,68^. Whilst rsBOLD_SD_ could be associated to individual differences^69^, the collective increase with statistical significance in rsBOLD_SD_ post-FUS observed may reflect elevated metabolic demand associated with increased both excitatory and inhibitory synaptic input and firing rates^70^ (**Fig. 4e**). Yet, this does not provide specificity in whether the effects were caused predominantly by either the excitatory or inhibitory synaptic inputs. To further evaluate the effects of STN-FUS, amplitude of low frequency fluctuations (ALFF)^71^ obtained by the power spectral density of the time series constrained to 0.01-0.10 Hz highlights changes in the default-mode network (DMM)^70,72,73^. Significant increase in concentrated energy was observed below 0.03 Hz with STN-FUS, which is connected to lower frequencies of the EEG band correlated to sleep^71^ (**Fig. 4f**). As the STN plays a key role in the basal ganglia network, we used arterial spin labeling (ASL)^74^ examined the effects of STN-FUS on cerebral blood flow (CBF) at various associated brain regions (**Fig. 4g**). Significant ipsilateral increase in CBF was observed at the left nucleus accumbens (NAc), potentially indicating heightened metabolism and suggesting limbic disinhibition downstream of the STN. As a result of the expected inhibition of the STN by FUS, the suppressed output of glutamatergic (GLU) projection to the globus pallidus interna (GPi), globus pallidus externa (GPe), substantia nigra pars reticulata (SNr), substantia nigra pars compacta (SNc) and ventral medial prefrontal cortex (vmPFC) (**Fig. 1c**) may result in greater excitatory drive of the NAc^75^.

Resting-state functional connectivity (rsFC) provides a helpful basis for understanding changes in resting-state networks due to brain stimulation^76^. To further evaluate the effects of STN-FUS, a group-level analysis of rsFC with the L-STN as the seed-region was performed with linear-mixed model effects and probabilistic threshold-free clustering enhancement (pTFCE)^77^. The ipsilateral (left) GPe, GPi, and Putamen (Pu) was positively connected to the STN, whereas the contralateral (right) GPe, GPi, and Pu remained negatively connected (**Fig. 4h**). The results align with the effects of STN-DBS, where an increase in rsFC in GPe, GPi, and Pu ipsilateral to the site of stimulation was induced. As the STN-GPe serves as the self-regulating pacemaker to the cortico-basal ganglia-thalamo-cortical (CBGTC) loop^78^, the observed increase in positive rsFC between GPe and STN is consistent with known mechanism, in which the GPe’s tonic GABAergic (GABA) inhibition to the STN reflects a likely inhibitory dynamic in the STN due to FUS^79^. Additionally, that the left NAc’s rsFC trended negatively in parallel with increased CBF further suggests STN-FUS potentially affects the limbic loop, which plays a crucial role in sleep regulation aside from the basal ganglia network^80^. Collectively, these findings indicate a potential alteration and elevation in the inhibitory-excitatory regulation between the STN and other basal ganglia and limbic structures, as evidenced by changes in connectivity patterns following FUS stimulation. Through the changes in connectivity patterns of the STN-FUS, these may indicate neuromodulatory effects of STN-FUS in support of improving sleep regulation shown in STN-DBS^49^.

### NEUSLeeP-enabled STN-FUS enhances REM sleep performance

Two consecutive nights of sleep study were performed to evaluate the efficacy of STN-FUS with NEUSLeeP (**Fig. 5a**). Two groups of subjects (healthy/insomnia) were screened and enrolled, with the criteria of a score less than 5 on the PSQI indicating inclusion in either the healthy or between 6-10 indicating inclusion for insomnia groups. Participant evaluation also included a series of clinical questionnaires (**Fig. 5b**) and vital measurements and heart rate variability (HRV) extracted from electroencephalography (ECG) in the evening prior to the sleep session and post-morning to monitor unexpected or adverse effects of STN-FUS (**Supplementary Fig. 6-7, Supplementary Table. 3-4**). Subjects were exposed to the sham condition on the first night and STN-FUS the second night followed by fMRI scans in the morning subsequently due to the uncertain long-term effects of FUS potentially affecting the sham condition^81,82^ (see **Methods**).

**Figure 5.**
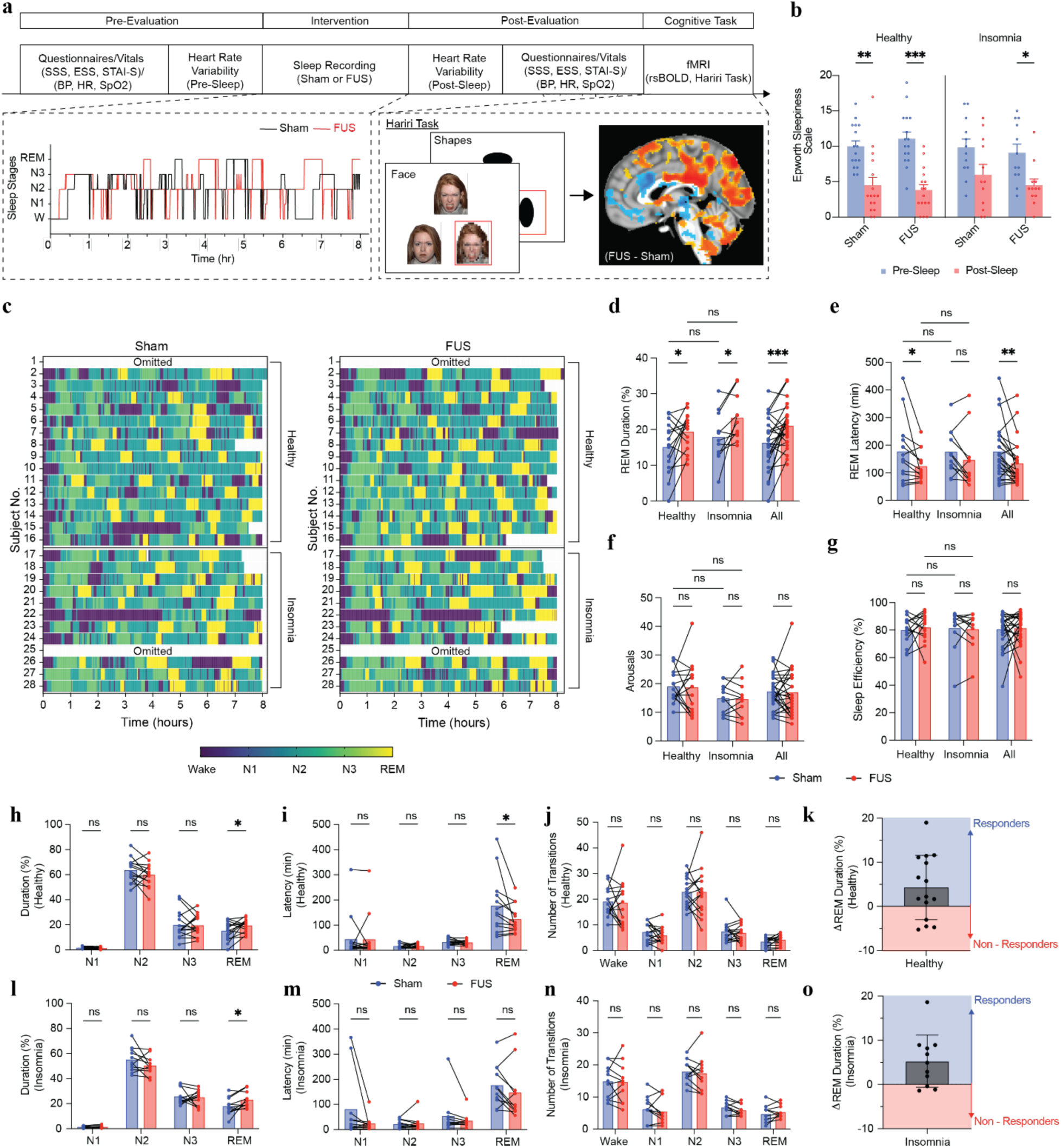
NEUSLeeP-enabled STN-FUS enhances REM and sleep performance. **a)** Experimental protocol of overnight sleep recording and stress measurements under STN-FUS using NEUSLeeP. **b)** Clinical questionnaire of Epsworth Sleepiness Scale response from both healthy and insomnia populations (n = 16 and 12, Two-way ANOVA and Tukey’s multiple comparison correction). **c)** Hypnogram of healthy and insomnia group on sham and FUS night (n = 16 and 12). **d)** REM duration normalized to total sleep duration (n = 16 and 12, Two-way ANOVA and Tukey’s multiple comparison correction). **e)** REM latency decreased (n = 16 and 12, Two-way ANOVA and Tukey’s multiple comparison correction). **f)** Number of arousals (wake) was not changed suggesting no adverse effects in sleep quality from NEUSLeeP (n = 16 and 12, Two-way ANOVA and Tukey’s multiple comparison correction). **g)** Sleep efficiency was not changed with NEUSLeeP indicating total sleep duration (N1, N2, N3, REM) was not affected (n = 16 and 12, Two-way ANOVA and Tukey’s multiple comparison correction). **h**) REM duration percentage changes across sleep stages in healthy population. **i**) REM latency changes across sleep stages in healthy population. **j**) Number of transitions in sleep stages in healthy population. **k**) Responders and non-responders of NEUSLeeP-enabled STN-FUS in healthy populations. **l**) REM duration percentage changes across sleep stages in insomnia population. **m**) REM latency changes across sleep stages in insomnia population. **n**) Number of transitions in sleep stages in insomnia population. **o**) Responders and non-responders of NEUSLeeP-enabled STN-FUS in insomnia populations. All plots show mean ± s.e.m unless otherwise mentioned, *P < 0.05, **P < 0.01, and ***P < 0.001. All data generated in this figure are provided in the Source Data file. Created in BioRender. Tang, K. (2025) https://BioRender.com/xi5ugz1

A quantitative approach in determining the effects of REM enhancement with STN-FUS using polysomnographic evaluation with EEG data recorded with NEUSleeP was performed (see **Methods**). Here, a total of 28 subjects across both groups’ (healthy, n = 16; 8 males and 8 females / insomnia, n = 12; 6 males and 6 females) polysomnographic data staged by a certified trained sleep expert were presented (**Fig. 5c**). Two subject’s polysomnography data were omitted due to extensive noise across all channels rendering it unusable (Subject No. 1 and 25). REM duration was observed to significantly increase on nights with STN-FUS compared to sham (**Table. 1**, Pressure: 0.90 MPa, 30s ON 30s OFF, Pulse Duration = 0.5 ms, Pulse Repetition Frequency = 100 Hz) for both healthy group (15.1 ± 7.38%; Sham / 19.4 ± 5.25%; FUS) and insomnia group (17.902 ± 7.10%; Sham / 23.2 ± 6.47%; FUS) (**Fig. 5d**). REM latency observed to significantly for healthy group (177 ± 117 min; Sham / 123 ± 57.1 min; FUS) but minimally for insomnia group (176 ± 82.2 min; Sham / 146 ± 106 min; FUS) (**Fig. 5e**). In addition, STN-FUS was not associated with changes in number wake arousals (healthy: 19.1 ± 5.56; Sham / 18.7 ± 9.08 min; FUS, insomnia: 14.8 ± 5.23; Sham / 14.6 ± 6.12 min) and overall sleep efficiency (healthy: 80.0 ± 9.97%; Sham / 81.4 ± 10.6%; FUS, insomnia: 81.4 ± 15.7%; Sham / 80.6 ± 13.4%) (**Fig. 5f-g**). Overall, the observed effects of NEUSLeeP enabled STN-FUS in 26 subjects suggests 25% increase in REM sleep duration (from 63.8 ± 29.9 min to 79.7 ± 26.0 min) and 43 min reduction in REM latency (from 177 ± 101.1 min to 134.7 ± 82.3 min), without significantly altering the sleep architecture in other sleep stages such as Wake, N1, N2, N3 with majority of the subjects responding positively in REM improvement (**Fig. 5h-o, Table. 2**).

**Table. 2.** Summary of sleep performance metrics with NEUSLeeP.

| Group | Metrics | Sham | FUS | Cohen's d | p-value |
| --- | --- | --- | --- | --- | --- |
| Healthy<br>(n = 15)<sup>1</sup> | REM Duration (min) | 58.9 ± 30.6 | 74.9 ± 24.6 | 0.58 | 0.0280* |
|  | REM Duration (%) | 15.1 ± 7.38 | 19.4 ± 5.25 | 0.67 | 0.0164* |
|  | REM Latency (min) | 177 ± 117 | 124 ± 57.1 | -0.57 | 0.0165* |
|  | Arousals | 19.1 ± 5.56 | 18.7 ± 9.08 | 0.05 | 0.8278 |
|  | Sleep Efficiency (%) | 80.0 ± 9.97 | 81.9 ± 10.6 | 0.18 | 0.6392 |
| Insomnia<br>(n = 11)<sup>2</sup> | REM Duration (min) | 70.4 ± 29.1 | 86.1 ± 27.6 | 0.55 | 0.065 |
|  | REM Duration (%) | 17.9 ± 7.10 | 23.2 ± 6.47 | 0.78 | 0.0113* |
|  | REM Latency (min) | 176 ± 82.2 | 146 ± 106 | -0.31 | 0.2090 |
|  | Arousals | 14.8 ± 5.23 | 14.6 ± 6.12 | 0.03 | 0.9325 |
|  | Sleep Efficiency (%) | 81.4 ± 12.4 | 81.4 ± 11.6 | 0.05 | 0.8659 |
| All<br>(Healthy +<br>Insomnia)<br>(n = 26) | REM Duration (min) | 63.8 ± 29.9 | 79.7 ± 26.0 | 0.57 | 0.0049*** |
|  | REM Duration (%) | 16.3 ± 7.26 | 20.9 ± 5.99 | 0.71 | 0.0007**** |
|  | REM Latency (min) | 177 ± 101 | 134 ± 82.3 | -0.46 | 0.0098*** |
|  | Arousals | 17.3 ± 5.73 | 16.9 ± 8.08 | 0.04 | 0.8256 |
|  | Sleep Efficiency (%) | 80.6 ± 12.4 | 81.4 ± 11.6 | 0.06 | 0.7598 |

<sup>1</sup>Due to poor EEG signal quality recording, subject #1 from healthy group was omitted.
<sup>2</sup>Due to intermittent bruxism and poor EEG signal quality recording, subject #25 from insomnia group was omitted.

Changes in high-level sleep architecture and dynamics could provide key insights into the mechanisms by which STN-FUS may impact sleep as well as differences in healthy and insomnia groups^83^. Specifically, sleep stage state transition probability demonstrates changes across sleep apnea^84^, aging^85^ and insomnia^86^. Here, a collective group-level state transition probability matrix was computed (see **Methods**) for each condition (sham/FUS) and groups (healthy, insomnia) (**Supplementary Fig. 8a-c, 8d-f**). At first glance, both matrices resemble one another greatly. However, taking the difference between sham and FUS condition matrices provides additional insight into key differences, predominantly in transition from N3 and REM. On nights of STN-FUS vs. sham, the healthy group tended to decrease occurrence in transition from N3/REM to the wake state but increase frequency of transitions to the N2 state (**Supplementary Fig. 8c**). In contrast, the insomnia group showed the opposite effect, specifically increased chance of transition from N3/REM to wake and decreased chance of transition from N3/REM to N2 (**Supplementary Fig. 8f**). This aligns with the pathology of insomnia given their susceptibility to arousals^87^. Even though an observed increase in arousal transitions was observed, the differences in change are statistically non-significant and inconclusive (see **Methods**). Despite the differences in groups, an overall increase in N2 to REM probability was observed while the remaining transitions maintained similar frequencies, consistent with the observed REM enhancement shown before. To evaluate the broad effects of STN-FUS, the Frobenius Distance provides insight in the difference between sham and FUS matrices, where higher values indicate greater effects and/or differences^88^. Fundamentally, nights of STN-FUS with NEUSLeeP showed a difference compared to sham (**Supplementary Fig. 8g**). Furthermore, this change extends similarly to both healthy and insomnia groups (**Supplementary Fig. 8h**). Similarly, to ensure sleep architecture remained unchanged broadly, stage-specific Chi-squared test and Mantel test (see **Methods**) were performed to evaluate similarity, in which the transition matrices remained very similar indicating unchanged sleep architecture. Overall, STN-FUS was not associated with an overall change in sleep architecture except for the selective enhancement of REM sleep. Together, these results provide initial compelling evidence of NEUSLeeP’s capacity for delivering STN-FUS to selectively enhance REM sleep performance while maintaining the general sleep architecture of individuals.

### Stress Response of NEUSLeeP-enabled STN-FUS

We further explored the effects of NEUSLeeP-enabled STN-FUS on stress response. The morning following each of the two consecutive sleep studies with NEUSLeeP, individuals were subjected to a sequence of fMRI scans in conjunction with a widely-utilized emotional face-matching task that serves as a validated probe of limbic circuitry^89^. We reasoned that an improvement in REM sleep may also translate to a reduction in reactivity to emotional stimuli in limbic regions, such as the amygdala, which play a key role in the detection of salient environmental stimuli and in triggering behavioral and physiological defensive threat responses. Paradigms such as the emotional face matching task we utilized here employ fearful and angry facial expressions as salient environmental cues to probe this amygdala-mediated defense threat circuit, which is similarly responsive to other legitimately threatening stressors such as confrontational or life-threatening conditions^90^. Furthermore, previous work implementing FUS for amygdala neuromodulation demonstrated that this paradigm was sensitive to potential persisting changes in amygdala function following a three-week course of repeatitive daily FUS, which itself was associated with symptomatic improvements across mood, anxiety, and trauma-related disorders^45^.

We began with amygdala ROI-constrained analyses to evaluate the main effects of FUS (FUS vs. Sham) on amygdala task-dependent responses to experimental stimulus condition blocks (Anger/Fear/Happy/Neutral/Shapes) and as a function of patient group (Healthy/Insomnia). Here, no significant FUS-related main effect was observed on either left/right amygdala BOLD activation (see **Methods**) in the healthy group (**Fig. 6a**). There was a main effect of group, such that the insomnia group exhibited overall elevated amygdala activity across all task conditions bilaterally, commonly associated with sleep disorders^91^. Interestingly, FUS was associated with increased amygdala activity to anger stimuli in the insomnia group only (Left Amygdala/Insomnia, Anger x [FUS vs. Sham]), p = 0.0045). Further ROI-constrained analyses of BOLD signal were extended to localized basal ganglia structures targeted (STN, SNc/SNr). Whilst mechanisms are unclear, the observed increase in amygdala activity could potentially be explained by the activation of subthalamic neurons through contralateral subthalamic neuromodulation demonstrated in Parkinson’s^92–94^. However, SNc in healthy groups exhibited ipsilateral attenuation of BOLD signal to anger stimuli (Left SNc/Healthy, Anger x [FUS vs. Sham]), p = 0.0419) with a similar statistical trend in response to happy stimuli (Left SNc/Healthy, Happy x [FUS vs. Sham], p = 0.0088). Similarly, the left SNr in healthy groups showed attenuation of response to anger stimuli (Left SNr/Healthy, Anger x [FUS vs. Sham]), p = 0.0264), fear stimuli (Left SNr/Healthy, Fear x [FUS vs. Sham]), p = 0.0354), and happy stimuli (Left SNr/Healthy, Happy x [FUS vs. Sham]), p = 0.0193).

**Figure 6.**
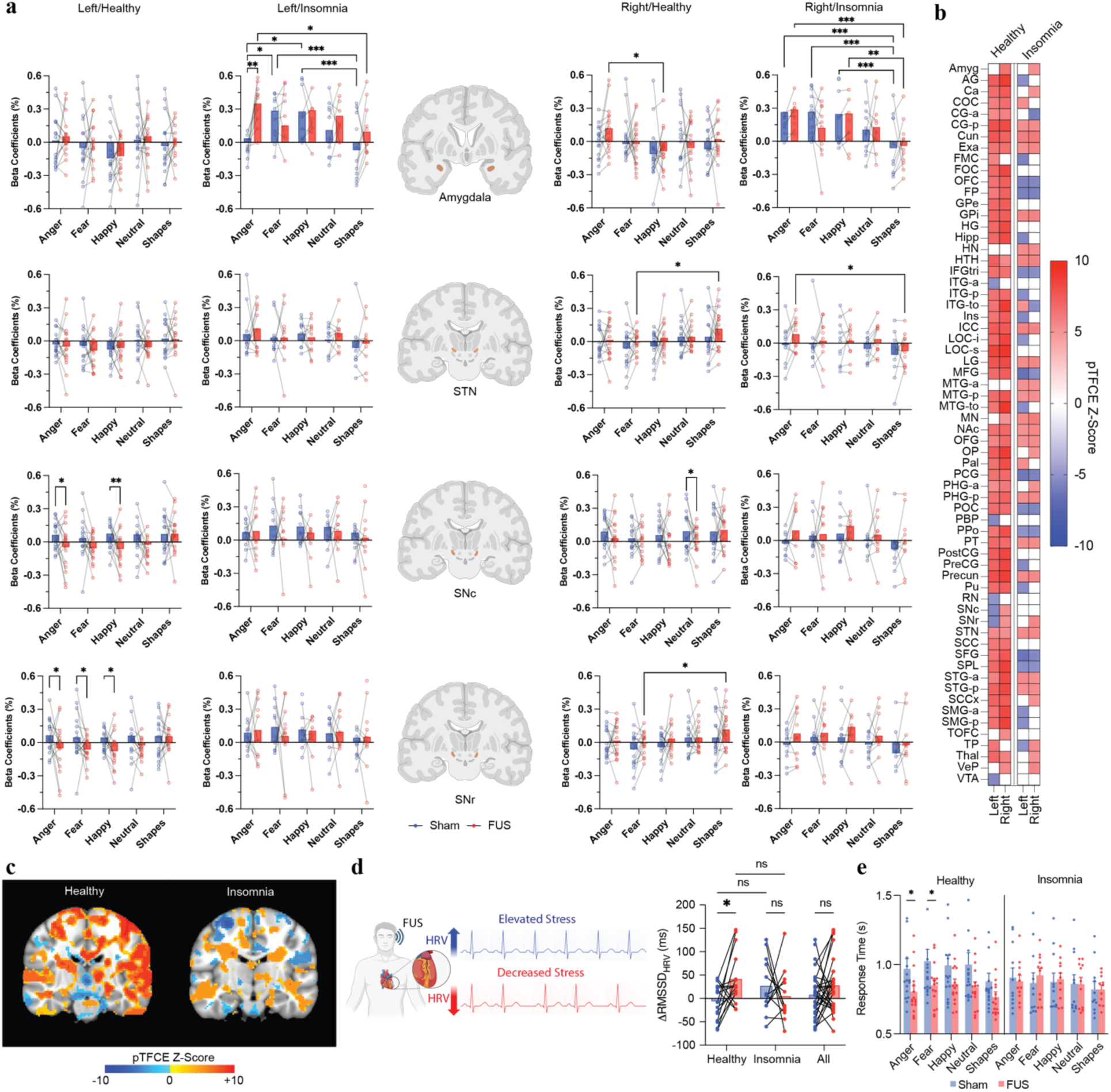
Stress adaptation and response with NEUSLeeP-enabled STN-FUS during Hariri Tasks. **a)** Effects of NEUSLeeP under emotion-based and facial-based stimuli (Hariri Task) under varying conditions (Anger, Fear, Happy, Neutral, Shapes) in left/right ipsilateral Amygdala, STN, SNc, and SNr for healthy and insomnia groups. Significant decreased activity in the left SNc/SNr under emotional salient stimuli were observed in healthy group (n = 16, Two-way ANOVA and Tukey’s multiple comparison correction). Significant elevated activity across all conditions in the Amygdala bilaterally observed in insomnia groups (n = 12, Two-way ANOVA and Tukey’s multiple comparison correction). **b)** Effects of STN-FUS comparing FUS – Sham indicates significant decreased activity the left ITG-a, PBP, RN, SNc/SNr, and VTA with elevated cortical activity was observed in healthy groups. Contrarily, insomnia groups had substantial inhibitory and excitatory effects across the brain (n = 16 and 12, linear mixed effect models with probabilistic threshold-free cluster enhancement). pTFCE values were zeroed between -4.90 and 4.90 for statistical representation. **c)** Sagittal image of voxelwise group-level analysis with pTFCE under conditions in healthy and insomnia groups. **d)** Change in heart rate variability by root mean square of successive differences of R-peak waves in ECG obtained between evening (pre-sleep) and morning (post-sleep) (n = 16 and 12, Two-way ANOVA and Sidak’s test). **e)** Response time during Hariri Task (n = 16 and 12, Three-Way ANOVA with Tukey’s multiple comparison correction). Significant decrease in response time during Anger and Fear conditions were observed in healthy groups. All plots show mean ± s.e.m unless otherwise mentioned, *P < 0.05, **P < 0.01, ***P < 0.001, and *pTFCE > |4.90|. All data generated in this figure are provided in the Source Data file. All data generated in this figure are provided in the Source Data file. Created in BioRender. Tang, K. (2025) https://BioRender.com/xi5ugz1

Evaluation of interaction between task-dependent stimuli within conditions (i.e. FUS x [Anger vs. Shapes]) was performed and exhausted through a whole-brain analyses approach using the Harvard-Oxford atlas^95–98^. The effects of left STN-FUS at the left ipsilateral localized region comprised of SNc, SNr, and STN was observed with attenuated activity in healthy groups. Two-way ANOVA with Bonferroni correction for multiple comparisons revealed significant attenuation specifically for fear and happy stimuli (**Supplementary Fig. 9**). Contrariwise, the insomnia group did not show significant changes at these regions. However, a significant elevation in activity across all conditions was observed in limbic associated regions (amygdala, intracalcarine cortex, lateral occipital cortex, and frontal operculum cortex) commonly associated with insomnia and sleep disorders, which may be indicative of known deficits in emotion and arousal regulation^99–102^.

Due to the potential heterogeneity in function of subregions of the STN, which consists of three distinct anatomical subregions responsible for motor, associative and limbic connectivity^103^, the ROI-constrained analyses may lack spatial information^104^. As such, to further assess the statistical significance of effects associated with STN-FUS in the whole-brain, voxelwise analysis conducted with pTFCE was conducted to evaluate the main effect of FUS vs Sham. The healthy group demonstrated a significant increase across the much of the brain apart from the left ITG-a, PBP, RN, SNc, SNr, and VTA (refer to **Supplementary Table 5.**) whereas the insomnia group showed a heterogenous mixture of increased and decreased activity across the brain (**Fig. 6b-c, Supplementary Fig. 10**). Additionally, connectivity analysis in left vs. right ipsilateral ROI in response to FUS vs. Sham was investigated, whereby differences were observed but inconclusive and beyond the scope of this work (**Supplementary Fig. 11**).

From an etiological perspective, stress detrimentally impacts sleep quality, effectively serving as a vulnerability factor for development of insomnia and circadian disorders^105^. The stress response is also highly complex and idiosyncratic, whereby presentation of identical stress-induced stimuli could result in different reactions across individuals^106,107^. It is therefore essential to identify and characterize stress through various metrics and levels of analysis. One of the most common assesses autonomic regulation of psychological stress through measuring the hypothalamic pituitary-adrenal (HPA) axis and sympathetic nervous system (SNS). Both play a key role in the sympathetic response to stress, i.e. the *fight or flight* response^108^. Given that heart rate is heavily influenced by fluctuations in the autonomic nervous system, the heart rate variability correlates inversely with stress levels^109^. Here, we examined the pre/post effects of NEUSLeeP STN-FUS vs. sham on HRV and found significantly increased changes in HRV in healthy group on mornings following STN-FUS vs. sham stimulation (-5.89 ± 38.4%; Sham / 41.5 ± 54.7%; FUS). Conversely, the insomnia group did not show a clear difference in HRV as a function of STN-FUS vs. sham (26.9 ± 38.4%; Sham / 4.79 ± 56.4%; FUS) (**Fig. 6d**). The lack of difference in HRV following STN-FUS vs. sham for insomnia groups aligns with several studies indicating no changes in HRV in patients with insomnia^110–112^ and even those with successful treatment of cognitive-behavioral treatment^113^ in relation to healthy individuals.

In addition, we also examined potential changes in behavioral metrics of emotion processing related to STN-FUS vs. sham stimulation, i.e. average response time during the emotional face matching task, as impaired behavioral efficiency of facial affect processing had been previously observed in sleep-deprived individuals^114,115^. Here, the healthy group demonstrated a reduction in response time specifically during processing of anger and fear task stimuli the morning following STN-FUS vs. sham (**Fig. 6e**). However, no improvement in response time was observed in the insomnia group. Overall, these results indicate that STN-FUS with NEUSLeeP may have the capacity to selectively down-modulate the ipsilateral basal-ganglia– midbrain–temporal circuit through the SNr/SNc, promote REM sleep, and modulate aspects of subsequent stress responses in healthy individuals.

## Discussion

Here, we present NEUSLeeP as novel wearable device that introduces a practical, mobile form factor for transcranial ultrasound that can be worn comfortably during overnight monitoring. In contrast to benchtop systems that constrain posture and movement, this platform enables stimulation to be delivered in naturalistic sleep, synchronized with stage detection, and repeated across the night. The current study observed that stimulation targeted to the left STN was associated with enhancement of REM-related metrics (e.g., higher REM proportion and/or shorter REM latency depending on subject), and neurobiological response patterns across brain regions suggested task-and site-selective effects rather than a global functional change. Specifically, changes were strongest in nodes that are anatomically poised to modulate REM regulation and affective processing (temporal association cortex and dopaminergic midbrain/basal ganglia), consistent with localized neuromodulation propagating through known sleep– affect networks^116–121^. While preliminary, the convergence of wearable feasibility, REM enhancement, and spatially selective effects argues that NEUSLeeP is not merely a portable stimulator; it is also a new experimental platform for state-dependent, targeted neuromodulation during sleep.

Several limitations warrant consideration. The present study did not fully counterbalance stimulation order, was single-blinded and not double-blinded, and lacked parallel control groups^122^ due to several factors: 1) constraint of population size permissible, 2) fixed project period and financial resources defined by funding mechanism, and 3) unclear durability of effects, as previous work observed only up to 40 minutes^123^, and whether a counterbalanced order would produce carryover effects. These choices increase exposure to expectancy and order effects: participants may sleep differently on the first vs. subsequent nights; circadian/ultradian variation and spontaneous night-to-night variability are substantial; and staff knowledge of condition can unintentionally influence coaching, sensor placement, or adherence. Without a within-subject, randomized, counterbalanced sham (or alternative-site) condition and double-blind masking (participant and scorer), we cannot rule out placebo or context effects. Future trials should adopt a randomized, sham-controlled, crossover design with concealed assignment, identical device cues (acoustic/thermal), automated stimulus scheduling, and pre-registered primary outcomes. Including an active control (e.g., stimulation to a non-sleep node or subthreshold intensity) would help separate site-specific effects from non-specific device effects.

Insomnia is characterized by hyperarousal, altered salience processing, conditioned wakefulness, and trait anxiety; these mechanisms vary across individuals and across nights. Standard block-design cognitive tasks (e.g., emotion–shape contrasts) probe selective aspects of cortical/subcortical processing but may only indirectly relate to sleep initiation/maintenance. The scanner context itself (noise, confinement) can exaggerate arousal and distort ecological validity, especially for insomnia. Thus, null or mixed fMRI findings do not necessarily contradict behavioral sleep improvements—they may reflect task– construct mismatch. Future studies might instead consider pairing the wearable intervention with EEG-fMRI sleep transitions, resting-state connectivity around sleep onset, or event-related paradigms that index arousal gating (auditory oddball, mismatch negativity) and emotion regulation rather than only facial affect processing and emotional reactivity. Subject-specific modeling (e.g., symptom clusters, chronotype, baseline arousal) and state-dependent analyses (time-locking to micro-arousals or NREM/REM boundaries) are likely to be more sensitive to the mechanisms by which stimulation impacts insomnia.

Current ultrasound neuromodulation experimental therapeutic protocols typically employ multi-session repeated dosing to promote clinical benefits^45,124–127^. A single night can inform feasibility and immediate effects, but it cannot speak to accumulation, consolidation, or durability of benefits. Ultrasound-induced neuromodulation may similarly require repeated, spaced sessions to reshape network excitability and sleep regulation. Future work should examine dose–response curves (intensity, duty cycle, burst structure), session schedules (daily vs. alternate-day; timing relative to circadian phase), and follow-up (1– 4 weeks) with pre-registered endpoint metrics to determine persistence and clinical meaningfulness. Adaptive algorithms that increase or taper stimulation based on nightly response (closed-loop dosing) could further improve outcomes as specific stage targeted stimulation of N2, N3 or REM using other modalities elicited improved sleep performances^11^.

The current line-of-sight approach and the spatial resolution achievable with the CRUTA method introduce two classes of risk: off-target sonication and network-mediated misattribution. Whilst characterization of the CRUTA in temporal bone was performed, variation across individuals is difficult to account for^128^. Skull curvature and heterogeneity can distort the beam, create secondary acoustic grating lobes or shift the focus; line-of-sight paths may favor superficial segments of a network over the nominal target; and the effective in situ point spread function can be larger than the anatomical target, particularly in deep structures. As a result, observed effects may partly arise from adjacent nuclei or fiber pathways (e.g., pallidal, thalamic, or cerebellar–brainstem circuits) that project into the measured region. To mitigate this, future versions should couple the wearable hardware to subject-specific CT/MRI modeling for phase correction and safety-bounded intensity estimates, validate focality with acoustic radiation force imaging and employ control targets (spatially proximal but functionally distinct targets) to estimate off-target contributions. Similarly, delivering FUS with NEUSLeeP while undering BOLD fMRI could be approached by ensuring NEUSLeeP’s MRI compatibility^129,130^. Reporting targeting error, predicted 3D intensity maps, and dose-at-target as predicted by emerging modeling software tools^48,131,132^ will aid interpretability. Where feasible, multi-element steering and closed-loop stage-locked delivery can narrow temporal and spatial windows, improving selectivity.

In conclusion, NEUSLeeP demonstrates that wearable ultrasound neuromodulation during natural sleep is feasible and can enhance REM-related outcomes while eliciting spatially selective neural effects consistent with localized modulation. With the limitations discussed in consideration, the work presented here offers a new platform and approach for leveraging ultrasound and material innovations to enable REM sleep measurement and potential enhancement for both healthy individuals and those suffering with insomnia. With additional future work, NEUSLeeP may transform from a promising prototype into a scalable, at-home neuromodulation therapy that complements behavioral treatment and pharmacotherapy by directly reshaping the neural dynamics that support healthy sleep.

## Methods

### Synthesis of Eco-PEIE-Gel

#### Materials and Preparation

Ecoflex-Gel was prepared by mixing the base elastomer with the cross-linking agent at a 1:1 ratio. The mixture was then added with a 5% weight ratio of the prepared mixture, which was then mixed uniformly with a digital stirrer at 1500 RPM. This was then poured into a desired mold and left to be cured for 48-hours followed by washing the surface under water before drying under ambient conditions for 12-hours.

### Synthesis of ASG hydrogel

To prepare the hydrogel precursor solution, 2 g of AMPS was fully dissolved in 2.5 g of deionized (DI) water under magnetic stirring until a homogeneous and transparent solution was obtained. Subsequently, 50 µL of a 0.05 M aqueous solution of MBAA was introduced as a crosslinker, followed by the addition of glycerol at a 30 wt % mass ratio relative to the total solution mass to enhance flexibility and water retention. The mixture was stirred thoroughly to ensure uniform distribution of all components. To initiate polymerization, 43 µL of a 0.097 M aqueous solution of TEMED was added as an accelerator, followed immediately by 32 µL of a 0.02 M freshly prepared APS solution as the radical initiator. The resulting precursor solution was rapidly mixed and carefully transferred using a micropipette into a custom-fabricated polylactic acid (PLA) mold produced via 3D printing. To prevent dehydration during the curing process, the filled mold was sealed inside a plastic enclosure. Thermal polymerization was carried out in a convection oven at 60 °C for 1 hour, ensuring complete gelation. After curing, the ASG hydrogel was gently demolded and stored in a sealed container prior to characterization and functional testing.

### Material characterization (Eco-PEIE-Gel and ASG)

#### Adhesion Strength/Cycle

The adhesion strength of Eco-PEIE-Gel and ASG was evaluated using modified ASTM F2255-05 and ASTM F2256-05 protocols with a custom-built testing system (FB5, Torbal) configured for 90° peel tests. Eco-PEIE-Gel and ASG specimens were fabricated with dimensions of 20 × 50 × 1 mm (width × length × thickness), and a Kapton film (7413D, 3M) was laminated to the backside of each sample to prevent stretching during peeling. For skin adhesion tests, Eco-PEIE-Gel and ASG were gently applied to the skin and peeled away at a 90° angle at a constant rate of 68 mm/min. Similarly, both Eco-PEIE-Gel and ASG were applied to a variety of materials including aluminum (Al), copper (Cu), iron (Fe), polyactic acid (PLA) plastic, polypropylene (PP) plastic, Ecoflex-0050, polydimethylsiloxane (PDMS), silicone (Si), and human skin (**Fig. 2d and 2k**.). Adhesion stability of Eco-PEIE-Gel prepared at 5% wt and 10% wt were performed similarly by repeated cycling of adhesion with 90° peeling up to 20 cycles on human skin.

#### Stress/Strain Tes

Tensile properties of the Eco-PEIE-Gel were characterized using the same custom-built mechanical testing apparatus (FB5, Torbal) employed for adhesion measurements. Samples were molded into rectangular strips with dimensions of 20 × 50 × 2 mm (width × length × thickness). Each specimen was clamped at both ends to minimize slippage, with Kapton film backing (7413D, 3M) applied when necessary to prevent stretching outside the gauge region. Tests were performed under uniaxial tensile loading at a constant crosshead speed of 68 mm/min until failure. Force–displacement data were continuously recorded and converted into stress–strain curves by normalizing the applied load to the initial cross-sectional area and the displacement to the initial gauge length (**Fig. 2f**). From these curves, the elastic modulus, tensile strength, and elongation at break were determined (**Fig. 2g-h**).

#### Electrode/Skin Impedance

The stability of the skin–electrode interface was evaluated over an 8-hour period using an impedance spectrum analyzer (SP-300, BioLogic). ASG electrodes were gently affixed to the forearm skin after standard cleaning with 70% isopropyl alcohol to minimize variability from surface oils and debris. A consistent electrode–skin contact area of 615 mm² was maintained across all tests, with the electrodes secured. Impedance spectra were collected at predetermined time intervals of every 2 hours across a frequency range of 1 Hz to 1 MHz using a sinusoidal excitation signal of 10 mV RMS. The analyzer recorded both magnitude and phase components of impedance, allowing extraction of resistive and capacitive contributions of the skin interface. Throughout the 8-hour measurement window, the subject remained at rest in a controlled laboratory environment (temperature ∼22 °C, relative humidity ∼45%) to minimize physiological and environmental variability (**Fig. 2m**). Similarly, the impedance of electrodes independently was measured **(Fig.2l)**.

#### ATR-FTIR

Fourier transform infrared spectra were collected to characterize the chemical structure of Eco-PEIE-Gel samples prepared with different PEIE concentrations (0, 2.5, 5, and 7.5 wt%). Films of each formulation were cast and cured under identical conditions and then cut into flat pieces (∼1 mm thickness) suitable for ATR measurement. Spectra were acquired using an ATR-FTIR spectrometer (Invernior R-FTIR, Bruker) equipped with a diamond/ZnSe crystal. Each sample was firmly pressed onto the ATR crystal to ensure consistent contact. Spectra were collected over a range of 4000–500 cm⁻¹ with a resolution of 4 cm⁻¹, averaging 256 scans per spectrum. Background spectra were recorded prior to each measurement and automatically subtracted. Spectral data were processed with baseline correction and normalization to facilitate comparison across different PEIE concentrations. Characteristic absorption bands corresponding to Si–O–Si, Si–CH₃, C–N, C–O–C, and N–H groups were analyzed to assess chemical interactions and compositional changes induced by increasing PEIE content (**Fig. 2c**).

### Design and Characterization of CRUTA

#### Simulation

Concentric ring array shape and its corresponding dimensions were first determined to be approximately between 60 -80 mm based on the focal depth from temporal window to the subthalamic nucleus^133^. This was then validated to be 72.2 ± 4.62 mm through anatomical MRI scans across 28 subjects (**Supplementary Fig. 3**). In a free-field, the maximum achievable focal depth *z_max_* for a flat concentric ring array transducer can be approximated by (1)^134^:

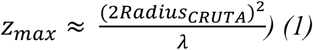

Assuming a maximum focal depth of 90 mm using center frequency of 650kHz and assuming the medium to be water (c = 1500 m/s), which is similar to that of brain tissue acoustically (*»* = speed of sound in water / center frequency) then the outer diameter of the CRUTA must be no less than 30 mm. In addition, to avoid off-targeting stimulation effects, minimization of grating lobes and high focality is necessary. Therefore, the trade-off between transducer size, ring width and number of rings was optimized through (2)^134^ requires the ring width of each element to be greater than 1.15 mm. To account for fabrication limitations, the ring width was selected to be 4.72 mm.

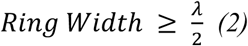

Given the constraints, using (3)^134^ to control adjustment of axial focal depth (*d_n_ : diameter of n^th^ element, λ: acoustic wavelength in medium, θ_n_: phase-delay of n^th^ element)* and (4)^134^ to determine the acoustic field pressure (*p_m_ : acoustic pressure at a 3D point, ρ: density of medium, c: speed of sound in medium, k: real wave number, A_n_: amplitude, ω: angular frequency, r_mnq_: distance to point m from point q with reference to element n, S_n_: surface area of element n)*, optimization and simulation using finite element analysis software for simulation (COMSOL Multiphysics 6.2, COMSOL Inc.) to determine acoustic field distribution at varying phase delays resulting in the design parameters used for CRUTA. (**Supplementary Table. 1,6**)

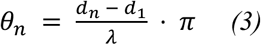

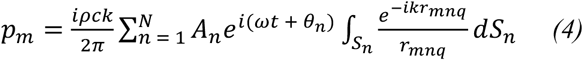

#### tFUS Waveform

Generation of tFUS acoustic profile from the CRUTA was performed using a research-grade high-intensity focused ultrasound stimulation system (Vantage 64LE HIFU, Verasonics Inc.). A train of pulse following the stimulation paradigm used in the study (Pulse Repetition Frequency: 100 Hz, Pulse Duration: 0.5 ms, Duty Cycle: 5%) was delivered at a center frequency of 650kHz (**Fig. 4a**).

#### Fabrication

A 3.5-mm thick 3D-printed PLA mold was attached to a medium tack transfer tape (Ultra 582U, TransferRite) where a silicon elastomer (Ecoflex-0050, Smooth On) was poured and cured for 30 minutes (**Supplementary Fig. 12a-b**). A 660µm thick copper adhesive tape (1126, 3M) was attached to water soluble tape (18C570, Aquasol) and medium tack transfer tape (Ultra 582U, TransferRite) were laser etched into interconnect patterns (**Supplementary Fig. 13a-c**), which was then carefully transferred printed on to 25.4 µm electrical grade polyimide film (2271K41, McMaster-Carr). Estimation of energy release between materials were identified using simplified Griffith’s equation^135^ known as the Kendall’s Peel Model^136^, which was used to determine feasibility of transfer printing copper onto polyimide, where the energy needed for the interface between the copper adhesive tape to polyimide film is much greater than the interface between the other two interfaces (**Supplementary Fig. 13d-f**). The copper interconnect laminated film was then gently laid on top of the cured elastomer mold (**Supplementary Fig. 12c-e**). A thin layer of Ecoflex-050 was applied to encapsulate interconnects exposing only the electrode terminals. Eight varying diameter piezoelectric rings with nickel coated electrodes (DL-47, Del Piezo) were acquired based on the designed dimensions (**Supplementary Table. 6**) and mounted onto copper electrode terminals with application of low-temperature solder paste (NP510-LT HRL1, Kester). The 3D-printed mold was removed and was placed in a thermo-controlled soldering oven for 5 minutes at 120°C. The 3.5 mm thick mold was reapplied. Subsequently, a 1-mm thick 3D printed PLA mold with a circular opening was placed on top and secured with mass weights. Eight pairs of 32-gauge wires were threaded through 6.35-mm silicon tubing into the circular opening, and the wires were soldered on the opening end of the copper interconnects (**Supplementary Fig. 12f**). Lastly, Ecoflex-050 was prepared and poured into the mold until uniformity for encapsulation and released from mold (**Supplementary Fig. 12g-h**).

#### Acoustic Field Mapping

The CRUTA was mounted on a 3D-printed submersible stand in a 457.2 mm x 180.34 mm x 139.7 mm (length x width x height) glass tank filled with degassed distilled water. Acoustic foams (Aptflex 48, Precision Acoustics) were padded internally along the glass walls to prevent scattering and reflection during measurements. Acoustic intensity and waveform were measured using a calibrated needle hydrophone (HNR-0500, Onda) mounted on a three-axis stage system with a workspace of 100 mm x 100 mm x 100 mm. The hydrophone was connected to an oscilloscope (SDS 1204-XE, Siglent) interfaced with a custom MATLAB program for axial, radial and automated 3D scanning with signal processing (**Supplementary Fig. 2a-b**). CRUTA was controlled and driven by a commercially available ultrasound system (Vantage 64LE HIFU, Verasonics Inc.). Each of the eight channels of CRUTA were connected independently via LEMO-00 and driven with varying delay timings with accordance to the desired focal depth (**Supplementary Fig. 2e**). 2D acoustic field scans in free-field were performed at 500µm increments (0 -100 mm from transducer in a 60 mm x 100 mm grid workspace) (**Fig. 3e**). Axial and radial profiles generated from the scans obtained were then processed to determine focal depth and spatial peak locations (**Fig. 3b-c**). In addition, a human skull (Skull Unlimited International Inc., 4-mm thick temporal cortical bone, rehydrated for 24 h in phosphate buffer solution) obtained was inserted in between the transducer and hydrophone (**Fig. 3a**), where similar 2D acoustic field scan was performed with an offset of 20 mm to prevent collision of the hydrophone with the skull (20 -100 mm from transducer in a 60 mm x 80 mm grid workspace).

#### Electrical Impedance

Each of the eight channels in CRUTA were proved with an impedance spectrum analyzer (SP300, BioLogic) using two-electrode configuration. A total of eight CRUTA devices were measured. Impedance was measured between 0 to 1MHz to identify and validate resonant frequencies centered approximately at 650kHz (**Fig. 3b**).

#### Thermal Heating

To determine the effective pressure field on the temperature in tissue, the bioheat transfer equation was used^137^. Given the pressure field measured (**Fig. 3e**) in free field, the power deposition per unit volume could be determined at the focal depth desired (**Supplementary Fig. 14a-d, Supplementary Table. 7**). Surface temperature of CRUTA was measured using thermal infrared camera (One Edge, FLIR) throughout the stimulation process to determine nominal heating does not elicit any skin burns. *Ex vivo* tests by applying the device on human transcranial skull (Skull Unlimited International Inc., temporal bone, rehydrated for 24 h in phosphate buffer solution) and measuring the thermal effects at the endocranium section of temporal bone was performed to ensure thermal diffusion did not elicit thermal damages to the dura, vessels, and cortical brain structures^23^ (**Supplementary Fig. 14e-i**).

### Fabrication and integration of NEUSLeeP

NEUSLeeP consists primarily of two components: 1) EEG layer and 2) CRUTA layer. The EEG layer was first developed by 3D-printing a 1-mm thick PLA mold (**Supplementary Fig. 12i-j**), which was then poured into with Ecoflex-050 and cured to create a thin-substrate layer. Subsequently, copper interconnects were laser-etched and transfer-printed onto polyimide (Cu-Pi) using the method described earlier and laminated onto the substrate (**Supplementary Fig. 12k**). A 3.5-mm thick 3D-printed PLA mold was then mounted on top of the electrode terminals with the addition of a separate mold (**Supplementary Fig. 12l-n**) to create an opening during encapsulation. 40 mg of Eco-PEIE-Gel was then prepared and poured onto the substrate for encapsulation. The EEG layer was then left to be cured for 48-hours before rinsed under water to remove the excess formation of ethoxylate and uncured Ecoflex-Gel on the surface before drying in ambient conditions for another 24-hours. Afterwards, a 10-blade scalpel was used to carefully remove the substrate from the mold (**Supplementary Fig. 12o**). To finish, the fabricated CRUTA mentioned previously would be bonded at the circular opening with silicon adhesive (Sil-poxy, Smooth-On) for 30-minutes (**Supplementary Fig. 12p**).

### Sleep recording performance of NEUSLeeP

Three volunteers were recruited to wear the NEUSLeeP and simultaneously with the AntNeuro 32CH EEG headcap injected with electrically conductive gel (Signagel, Parker Labs). The NEUSLeeP was connected to the EEG amplifier via a bipolar box to the amplifier (eego mylab, AntNeuro) whereas the EEG headcap was connected directly. The subjects were then placed into the sleep lab, and an 8-hour recording session was performed from 12AM-8AM. The recorded data was then split into two datasets with pseudo-randomized naming for single-blinded sleep staging. The data sent to a certified sleep staging expert and manually staged at 30-second epoch windows to evaluate the similarity in EEG recording signal quality between NEUSLeeP and standard 10-20 commercial-grade EEG headcaps (**Fig. 2n-p, Supplementary Fig. 1**).

### Subthalamic Nucleus Target Engagement using CRUTA

#### Participants

The procedure was designed in accordance with the Declaration of Helsinki guidelines and relevant ethical regulations regarding human research. The Institute of Research Board (IRB) at the University of Texas at Austin and Human Research Protection Office (HRPO) at the Defense Advanced Research Projects Agency (DARPA) approved all experimental procedures under the study [STUDY0005703]. Sixteen healthy participants (8 male, 8 female, aged 19-38 with a mean age of 25.7 ± 6.3 years) were screened for contraindications and neurological impairments (**Supplementary Table. 2**).

#### Experimental Setup

Participants were received at the Biomedical Imaging Center at Health Discovery Building, University of Texas at Austin. Prior to the start of the session, baseline measurements of vitals including SpO_2_, heart rate, and blood pressure were obtained. They were then placed in the magnetic resonance imaging machine (3T Vida, Siemens) for an standard T1-weighted Magnetization Prepared Rapid Gradient Echo (MPRAGE) scan followed by a Fast Gray Matter Acquisition T1 Inversion Recovery (FGATIR)^138^ scan to provide improved identification and localization of the subthalamic nucleus (**Fig. 4a**). Subsequently, a 10-minute pre-FUS resting-state functional (T2-weighted single-shot gradient-echo planar imaging, EPI) was performed to establish resting-state blood oxygen level dependent signals prior to stimulation. Next, a surrogate CRUTA device created by 3D-printed PLA and embedded with 50 mg of gadodiamide (APExBio) dissolved in PBS encapsulated with Eco-PEIE-Gel was attached to the zygomatic arch initially. Iterative rapid gradient scans and positioning of the surrogate device were performed until alignment and line-of-sight was verified towards the subthalamic nucleus. Additionally, distance between the device and STN was measured to determine focal depth by changing the delay parameter (**Fig. 4b**). Thereafter, medical markers were used to mark the position before removal of the surrogate device and attachment of the CRUTA. The CRUTA was connected to the ultrasound system (Vantage 64LE, Verasonics) via LEMO-00 connectors. Blood pressure was measured once again before FUS to ensure no effects were caused during the MRI. Participants were then subjected to 10 minutes of FUS (**Table. 1**, Pressure: 0.90 MPa, Pulse Repetition Frequency: 100 Hz, Pulse Duration: 0.5 ms, Duty Cycle: 5%, 30s ON 30s OFF) whilst simultaneously measuring their SpO_2_ and heart rate every minute to ensure no adverse effects due to FUS (**Fig. 4c**). Upon completion of FUS intervention, the device was removed, and participants were brought back to perform a post-FUS resting state functional scan. Follow-up emails were sent to participants one-week after the session to check-up and ensure no detrimental health effects were present due to FUS.

### Stress Adaptation and REM Enhancement Evaluation Post-FUS with NEUSLeeP

#### Participants

The procedure was designed in accordance with the Declaration of Helsinki guidelines and relevant ethical regulations regarding human research. The Institute of Research Board (IRB) at the University of Texas at Austin and Human Research Protection Office (HRPO) at the Defense Advanced Research Projects Agency (DARPA) approved all experimental procedures under the study [STUDY0005703]. All participants recruited were subjected to several screening processes involving compliance with MRI for health and safety, PSQI questionnaire to evaluate and assign study groups, health history and demographics. The study involves two groups: 1) healthy (PSQI < 5), and 2) insomnia (PSQI 6-10). The healthy group consists of sixteen participants (8 male, 8 female, aged 19-38 with a mean age of 22.7 ± 5.4 years) and insomnia group consists of twelve participants (6 male, 6 female, aged 19-38 with a mean age of 25.4 ± 5.9 years) (**Supplementary Table. 3-4**). All participants were screened for contraindications and neurological impairments.

#### Anatomical Measurements

All participants recruited and enrolled in the study were subjected to a session of anatomical scan prior to the sleep study. Subjects were placed in the magnetic resonance imaging machine (3T Vida, Siemens) for a standard T1-weighted Magnetization Prepared Rapid Gradient Echo (MPRAGE) scan. Next, a surrogate CRUTA device created by 3D-printed PLA and embedded with 50 mg of gadodiamide (APExBio) dissolved in PBS encapsulated with Eco-PEIE-Gel was attached to the zygomatic arch initially. Iterative rapid gradient scans and positioning of the surrogate device were performed until alignment and line-of-sight was verified towards the subthalamic nucleus. Additionally, distance between the device and STN was measured to determine focal depth by changing the delay parameter (**Fig. 4b**).

#### Sleep Study with NEUSLeeP

The sleep study consists of two consecutive overnight sessions wearing the NEUSLeeP, where the first night is sham and the second night with FUS (**Fig. 5a**). Participants were received at the Sleep Lab in the Institute of Mental Health at The University of Texas at Austin. Clinical questionnaires including Sleepiness Stanford Scale (SSS)^139,140^, Epsworth Sleepiness Scale (EPS)^141^, and State-Trait Anxiety Inventory (STAI-S)^142^ were given before the session to determine secondary outcome of individual’s perception of sleepiness and anxiety between sham and FUS. Vitals including blood pressure, heart rate and SpO_2_ were also collected before to establish as baseline measurement for comparison using a finger pulse oximeter (CMS60D, FaceLake) and sphygmomanometers (Track, iHealth). Subjects were then asked to wear the NEUSLeeP device with the assistance of research assistant personnel using the measured position performed in *Anatomical Measurements*. Ultrasound gel (Aquasonics 100, Parker Labs) was applied generously and topically to the CRUTA surface of the NEUSLeeP to ensure sufficient coupling between the transducer and the skin. Four gold-plated cup electrodes (630-020-60, BrainMaster Technologies) were additionally placed and injected with conductive gel (Signagel, Parker Labs) at Oz, Pz, Cz, and Fz based on the international 10-20 EEG system^143^ measured using a measuring tape from the nasion to inion. Similarly, a conductive adhesive hydrogel electrode (31050, Kendall) was placed laterally to the outer canthi of the right eye. The additional electrodes were fixated with medical tape (Nexcare, 3M) and were used to serve as a supplementary channel in assisting ground-truth sleep staging and for simultaneous comparison with signal quality compared with the ASG on NEUSLeeP. A pair of bipolar electrodes (Red Dot, 3M) was applied on the subject’s lower section of sternum (avF) and left shoulder (aVL) following the Einthoven’s Triangle configuration^143^ to allow simultaneous ECG monitoring throughout the night for safety and heart rate variability measurements. NEUSLeeP was then connected to an electrode box with an electrophysiological recording amplifier (BrainAmp ExG, BrainVision), which was connected to a computer in a separate room through fiber optic cable via a radiofrequency (RF) shielded panel. Similarly, the NEUSLeeP was connected with eight BNC-LEMO-00 connectors through the RF panel to the ultrasound system (Vantage 64LE, Verasonics). Infrared cameras were used to monitor the subject’s sleep behavior and detailed logs were recorded throughout the night. In addition, white noise^144–147^ was played through a speaker (Dohm Nova, Yogasleep) with modulated 100 Hz square wave at less than 60dB^148^ to mask all external noises and chirping noises generated by the CRUTA during the 100 Hz stimulation with NEUSLeeP for both nights throughout the sleep session. Lights were turned off during the sleep session and temperature was adjusted in accordance with the subject’s preference within the range of 18°C to 24°C.

Upon completion of device and electrode setup, ECG was recorded for 5 minutes to obtain baseline “*pre-sleep*” heart rate variability. Subsequently, the 8-hour sleep recording session began. On the night of the sham session, no FUS was delivered from NEUSLeeP. On the second night with the FUS session, FUS was delivered with the ultrasound system (Vantage 64LE, Verasonics) using 100 Hz PRF protocol (**Fig. 4a**, **Table. 1**, Pressure: 0.90 MPa, Pulse Repetition Frequency: 100 Hz, Pulse Duration: 0.5 ms, Duty Cycle: 5%, 30s ON 30s OFF for 5 minutes) per block every 90 minutes for a total of five blocks overnight following an ultradian sleep cycle schedule^149,150^. Focal depth targeting was adjusted based on the previously measured depth of STN in the *Anatomical Measurements*. Triggers corresponding to the stimulation were recorded simultaneously. Recording will be stopped upon reaching 8-hour or when the subject wakes up and signals to the monitoring personnel.

In the morning, ECG recording will be performed again for a “*post-sleep*” heart rate variability comparison. Next, NEUSLeeP and other electrodes will be removed from the subject followed by completion and recording of SSS, EPS, STAI-S, blood pressure, heart rate, and SpO_2_. Subjects will then be escorted to the Biomedical Imaging Center at the University of Texas Austin for functional magnetic resonance imaging.

#### fMRI Post-Sleep with NEUSLeeP

Participants were inserted into the imaging suite (3T Prisma, Siemens) and a series of scans were performed to evaluate the effects post-sleep with NEUSLeeP (sham or FUS). An anatomical scan using standard T1-weighted Magnetization Prepared Rapid Gradient Echo (MPRAGE) followed by a 10-minute resting-state resting-state functional (T2-weighted single-shot gradient-echo planar imaging, EPI) was performed to establish resting-state blood oxygen level dependent signals. The subjects were then asked to perform an emotional face-matching task, which asks participants to matching varying shapes and emotions to a reference image on each trial. Trials were organized into 2 x 10-trial blocks of each of the following conditions pseudorandomly ordered: anger faces, fear faces, happy faces, neutral faces, and shapes (circles and ovals). This is a common paradigm used to assess neurobiological reactivity to salient emotional cues, which is one important component of stress adaptation and response (**Fig. 5a**).

#### Stanford Sleepiness Scale

Subjective sleepiness was assessed using the Stanford Sleepiness Scale (SSS), a validated self-report measures widely used in clinical and research settings. The SSS is a 7-point Likert-type scale in which participants rate their current level of sleepiness, ranging from 1 (“feeling active, vital, alert, or wide awake”) to 7 (“no longer fighting sleep, sleep onset soon, having dream-like thoughts”). Participants were instructed to complete the SSS at predefined time points during the experimental session, including immediately prior to the MRI scan.

#### Epsworth Sleepiness Scale

The ESS assesses general daytime sleepiness by asking participants to rate their likelihood of dozing off or falling asleep in eight common situations (e.g., sitting and reading, watching television, as a passenger in a car) on a scale from 0 (“would never doze”) to 3 (“high chance of dozing”). The total ESS score was calculated by summing across all eight items, yielding a range of 0–24, with higher scores indicating greater habitual sleepiness. Participants completed the ESS during the baseline assessment prior to the experimental protocol. Both questionnaires were administered in paper form or electronically using secure data capture tools, and participants were given standardized instructions to ensure comprehension. Completed questionnaires were reviewed for missing data, and total scores were computed according to published guidelines.

#### State-Trait Anxiety Inventory

The STAI-S is a 20-item self-report inventory designed to assess current (state) anxiety. Participants rated how they felt “right now, at this moment” on a 4-point scale ranging from 1 (“not at all”) to 4 (“very much so”). Items included both positively and negatively worded statements, and total scores (range: 20–80) were calculated by reversing positively worded items and summing all item scores, with higher scores indicating greater state anxiety.

#### Sleep Recording and Staging

EEG, EMG, EoG data recorded from NEUSLeeP and additional channels using at right EoG and Oz, Pz, Cz, Fz sampled at 500Hz and preprocessed to remove ECG/motion artifacts and baseline drift using a third order butterworth bandpass filter in EEGLab 2025. Recordings were manually scored in 30-second epochs by a trained rater blinded to the study condition, using American Academy of Sleep Medicine (AASM) guidelines. Each epoch was classified into one of five stages: wake (W), non-rapid eye movement sleep stages (N1, N2, N3) or rapid eye movement (REM) sleep, based on the characteristic of EEG, EoG, and EMG patterns. Epochs were scored as N1 based on low-amplitude mixed-frequency EEG with slow eye movements; N2 based on presence of sleep spindles and/or K-complexes; N3 by high-amplitude (>75 µV), low-frequency (< 4 Hz) delta waves; and REM sleep by low-amplitude mixed-frequency with rapid eye movements in EoG, and muscle atonia in EMG. Sleep staging data were then summarized and calculated for, total sleep time (TST), sleep efficiency, proportion of time spent in each sleep stage, number of arousals, number of transitions for each stage, and state transition probability matrices. These metrics were used for group comparisons by performing multiple paired t-tests.

#### Heart Rate Variability (RMSSD_HRV_)

5-minute ECG recordings were extracted and filtered using a third order butterworth bandpass filter using MATLAB 2024b between 5 to 15 Hz to enhance R -wave peak. The root mean square of successive differences (RMSSD_HRV_) were calculated from the filtered enhanced RR interval time series (5)^151,152^.

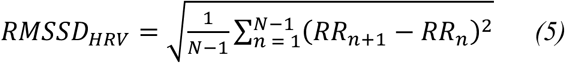

#### State Transition Probability Matrix

For each participant, a transition matrix was created by counting the number of transitions from state *i* to state *j*, where *i, j* ∈ {W, N1, N2, N3, REM}. Transitions were defined as consecutive changes between epochs, excluding self-transitions (i.e., remaining in the same stage). The raw counts were normalized by the total number of transitions from each originating state to obtain state-to-state transition probabilities (6).

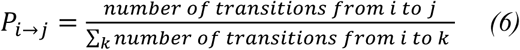

Group-level transition matrices were computed by summing individual matrices within each experimental condition (FUS vs. Sham). A difference matrix (FUS -Sham) was calculated to visualize absolute differences in transition counts between conditions (**Supplementary Fig. 8**). The overall magnitude of difference between group and individual matrices were quantified using the Frobenius norm (7)^153^

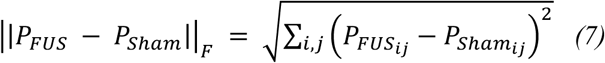

To assess row-wise differences in transition patterns (i.e., from each originating state), we performed chi-square tests of independence on each row of the aggregated transition matrices, comparing the distribution of outgoing transitions between sham and FUS conditions^154^. P-values were corrected for multiple comparisons using both Tukey’s correction and false discovery rate (FDR) correction (Benjamini-Hochberg procedure). At the matrix level, we computed the Mantel test to evaluate the similarity between the sham and FUS transition matrices. This test calculated the Spearman correlation between the off-diagonal elements of the two matrices and assessed significance using 10,000 label permutations^155^. All analyses were implemented in MATLAB (MathWorks, R2024b) using custom scripts.

### Functional Magnetic Resonance Imaging

Functional MRI data were analyzed using AFNI (version 24.3.06). Preprocessing included motion correction, slice timing correction, spatial normalization to the 1-mm slice MNI152 standard space, and spatial smoothing with a 6 mm full-width at half-maximum (FWHM) Gaussian kernel^156^. First-level statistical analyses were conducted using a general linear model (GLM), modeling task conditions as blocks convolved with a canonical hemodynamic response function and including six motion parameters as nuisance regressors (rotations and translations in the x, y, and z dimensions). Temporal autocorrelation correction and high-pass filtering (cutoff = 100 s) were applied. Individual contrast maps were carried forward to group-level analyses using AFNI’s 3dLME for linear mixed-effects modeling for mixed-effects estimation^157^. Subsequent statistical maps of interest for various effects were then subjected to probabilistic threshold-free cluster enhancement^77^ for enhancement of signal-to-noise ratio and for Type I error correction using a Gaussian random field correction on the pTFCE-enhanced Z-values using spatial smoothness metrics derived from subject-level model residual maps. All statistical maps were visualized on the MNI152 template using AFNI.

#### Resting-State Blood Oxygen Level Dependence Variability (rsBOLD_SD_)

BOLD time series was extracted from a predefined anatomical mask of the left subthalamic nucleus^158^. The ROI was defined in MNI152 standard space using a high-resolution subcortical atlas^95^ and resampled to match the spatial resolution of each subject’s preprocessed functional data. Functional images were temporally bandpass filtered (0.01 – 0.1 Hz) and detrended to remove low-frequency drifts. The left STN mask was then applied to each subject’s preprocessed 4D fMRI for both pre-FUS and post-FUS dataset. Resting-state BOLD variability was then obtained through (8) by taking the standard deviation of the BOLD time-series data extracted within-subject for interpreting activity and behavior at the left STN^159^ (**Fig. 4e**). The resulting time series served as the seed regressor for subsequent correlation analyses. Head motion parameters and white matter and CSF signals were included as nuisance regressors to reduce physiological and movement-related confounds.

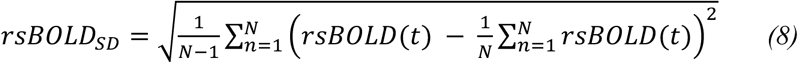

#### Resting-State Blood Oxygen Level Dependence – Amplitude of Low Frequency Fluctuations (rsBOLD_ALFF_)

ALFF in resting-state fMRI signals has been suggested to reflect the intensity of regional spontaneous brain activity^160^. As such, BOLD time series extracted from the left STN was filtered between 0.01 -0.10 Hz using a band-pass filter (MATLAB 2024b) to remove low-frequency drift and high-frequency respiratory and cardiac noise^161,162^. The power spectral density of the time series was then obtained through (9), where N is the number of voxels within the mask and [*FFT*(*rsBOLD*((*t*))] is the fast fourier transform of the BOLD time series within the mask^163^. The resulting power spectral density reflects the ALFF at the left STN, where paired t-test was used to evaluate statistical difference amongst each frequency bin (**Fig. 4f**).

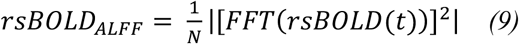

#### Resting-State Functional Connectivity

Within each subject, the left STN mask was used as the region-of-interest, serving as the seed region. A voxel-wise Pearson correlation between the seed region and every other brain voxel was calculated through AFNI. Correlation maps were then Fisher Z-transformed to generate a Fisher-Z score map. Group-level statistical modeling was performed using AFNI’s multi-variate model (multivariate modeling of variance) and linear mixed effect models, allowing the inclusion of both within-subject factors (e.g., condition: Pre-FUS vs. Post-FUS or Sham vs. FUS) and between-subject variables (e.g., group, sex). The output included voxelwise F-and t-statistics for main effects and interactions. The statistical map for each effect of interest was then subjected to probablistic threshold-free cluster enhancement to improve sensitivity to true effects and corrected for false positives using Gaussian random field correction on the pTFCE-enhanced Z values based upon spatial smoothness of the statistical effect map estimated by analysis of individual subject residual maps. Significant connectivity maps were visualized on the MNI152 template using AFNI. Selected ROI masks of brain structures associated with the basal ganglia circuit were used to mask and extract the mean t-score to determine the group-level effects of FUS compared to the sham condition (**Fig. 4g**).

#### Arterial Spin Labelling

Pseudo-continuous arterial spin labelling (pCASL) data were acquired using a 3T MRI scanner with a 64-channel head coil. A rapid gradient-echo scout was used to position the labeling plane between the carotid bifurcation and vessel 3 segment perpendicular to the carotidartery. pCASL data were then acquired using background-suppressed control/label pairs and a 3D gradient and spin echo (GRASE) echo-planar imaging (EPI) readout. FSL’s Brain Extraction Tool was used for brain extraction^164^ and custom python scripts were used for quantifications. Preprocessing included motion correction, brain masking, and spatial smoothing with a 6-mm FWHM Gaussian kernel. Pairwise subtraction of label and control images was performed to generate perfusion-weighted images, followed by calibration to compute quantitative cerebral blood flow (CBF) maps in units of mL/100 g/min. Standard quantification assumed a labeling efficiency of 0.85, T1 blood = 1.65 s, and a blood–brain partition coefficient of 0.9 mL/g. All CBF maps were registered to the MNI152 1-mm standard space using affine and nonlinear warping^165^.

#### Hariri Task

A widely utilized emotional face processing task^166,167^ measuring brain reactivity to salient emotional cues, which is one component of the stress response but not the only component, was used to test the effects of FUS (**Fig. 5a**). With reference to a similar previous study^45^, faces from the NimStim set were used and organized into blocks of 10 trials of various conditions, 2 blocks of each condition pseudo-randomly ordered (Anger, Fear, Happy, Neutral, Shapes) and balanced for gender and ethnicity^168^. The task involves both neutral face and shape comparator conditions to account and separate emotion-specific or face-processing effects. Each block consists of 10 trials, each trial lasting for 4-seconds involving individuals prompted to a reference face/shape (top) and two selection face/shape (bottom) for 3 seconds followed by a 1 second fixation cross between trials. Participants were asked to match the reference to one of the two selection faces/shapes using a button box. Tasks were presented using PsychoPy version 3.0.1 where visual stimuli were synchronized to scanner triggers, stimulus onset times and response times (**Fig. 6c**) were logged for subsequent modeling^169^. Simultaneously, functional MRI using EPI scans were conducted (T2-weighted single-shot gradient-echo planar imaging). Condition onset times were derived from PsychoPy logs and formatted to AFNI-compatible stimulus timing files. In addition to the condition regressors, the model included six motion parameters (three translational and three rotational components), baseline drift terms (Legendre polynomials up to third order), and censoring of TRs with motion exceeding 0.3 mm or outlier signal spikes.

Beta coefficients and t-statistics were estimated for each condition, and linear contrasts were defined to assess task-related effects (e.g., Anger vs. Shapes, Fear vs. Shapes). Two-way ANOVA with Tukey’s multiple comparison with each group (FUS/Sham) on the left and right ipsilateral ROI under each condition (Anger, Fear, Happy, Neutral) were compared to the control condition (Shapes) and labelled with an asterisk for p < 0.05. Individual-level contrast maps were carried forward to group-level analysis using linear-mixed effect models. The model incorporated within-subject factors (Condition, i.e. Anger vs Shapes) and between-subject factors (e.g., Group: FUS vs Sham). Voxelwise statistical maps for main effects and interactions were corrected for multiple comparisons using probabilistic threshold-free cluster enhancement (pTFCE) to improve sensitivity and true positive signal^77^. Individual-level model residual maps were used to estimate spatial smoothness using AFNI’s 3dFWHMx tool with the autocorrelation function, and pTFCE thresholds for the enhanced Z-value images were determined under Gaussian Random Field theory to control the family-wise error (FWE) rate (voxelwise FWE-corrected p < 0.05). Significant statistical maps were visualized on the MNI152 template using AFNI (**Supplementary Fig. 9-10**).

## Data availability

Additional data supporting the study’s findings are included in the article and its supplementary information and supplementary data. Source data are provided and included and can be found in the repository

## Code availability

The codes used for this study are available on a publicly available repository at GitHub https://github.com/kevintang725/DARPA-NEUSLEEP.

## Acknowledgements

We would like to thank Nanshu Lu for providing access to the use of the LPKF Protolaser U4 system from the Department of Aerospace Engineering and Engineering Mechanics, University of Texas at Austin. We would also like to thank David M. Schnyer for providing access and usage of the Sleep Lab at the Institute of Mental Health Research, University of Texas at Austin. Huiliang Wang would like to acknowledge support from the Defense Advanced Research Projects Agency (DARPA) (REM-REST) grant, Alzheimer’s Association New to the Field (AARG-NTF) research grant, UT Austin Proof of Concept Award nad Cockrell Innovation Grant. Kai Wing Kevin Tang would like to acknowledge support from the Jess Hay Chancellor’s Graduate Student Research Fellowship and Gregg and Marilyn Harris Endowed Graduate Fellowship. We acknowledge BioRender.com for the figures drawing.

## Author Contributions

K.W.K.T. and H.W. conceptualized the idea and designed the study. K.W.K.T. led the design, development/fabrication and characterization of NEUSLeeP device (Eco-PEIE-Gel and CRUTA); mechanical and chemical characterization of ASG; human clinical studies in STN target engagement with CRUTA, sleep studies, fMRI studies and their analysis. B.B. provided staging of sleep recording and advised on sleep study design. M.M.Y, J-C.H developed and optimized the ASG. W.D.M.B., M.M.Y., J.J. assisted with human clinical studies in STN targeted stimulation/validation. W.D.M.B., M.X.Y., D.S., T.C., J.G., R.M. assisted with human clinical sleep studies. J.J., M.M.Y., A.R.L., and I.P. assisted with fMRI studies. I.P. assisted with IRB submission and reviews. A.R.L. assisted with animal tests in H&E staining. J.W. and A.B. assisted with arterial spin labeling analysis. W.L. provided consultation in stimulation paradigm design. K.W.K.T led the writing of the original manuscript, with contributions from all other authors. V.M., G.F., and H.W. supervised the project.

## Ethics Declaration

All procedures involving human participants were conducted in accordance with the ethical standards of the Declaration of Helsinki. Ethical approval for this study was obtained from the Institutional Review Board (IRB) at The University of Texas at Austin [STUDY0005703]. Written informed consent was obtained from all participants prior to their involvement. Participant confidentiality and data protection were rigorously maintained throughout the study. The study enrolled both healthy individuals and individuals with self-reported insomnia symptoms through PSQI screening who had not received a formal clinical diagnosis. No vulnerable populations were specifically targeted, and all participants could provide informed consent. Participation was voluntary, and no coercion or undue influence was applied during the recruitment process.

This study is registered at ClinicalTrials.gov under registration number [NCT07190287], where detailed study protocols and outcome measures are available. No compensation was provided beyond standard reimbursement for participation-related expenses. Anonymized datasets generated during the current study, along with supporting materials, will be made available from the corresponding author upon reasonable request, in accordance with institutional guidelines and participant consent.

## References

1. Mysliwiec, V. et al. Sleep disorders and associated medical comorbidities in active duty military personnel. Sleep 36, 167–174 (2013).

2. Troxel, W. M., et al. Sleep in the Military: Promoting Healthy Sleep Among U.S. Servicemembers. (Rand Corporation, 2015).

3. Seelig, A. D. et al. Sleep and Health Resilience Metrics in a Large Military Cohort. Sleep 39, 1111– 1120 (2016).

4. Adjaye-Gbewonyo, D., Ng, A. E. & Black, L. I. Sleep Difficulties in Adults: United States, 2020. (2022) doi:10.15620/cdc:117490.

5. Alhola, P. & Polo-Kantola, P. Sleep deprivation: Impact on cognitive performance. Neuropsychiatric Disease and Treatment 3, 553 (2007).

6. Nollet, M., Wisden, W. & Franks, N. P. Sleep deprivation and stress: a reciprocal relationship. Interface Focus (2020) doi:10.1098/rsfs.2019.0092.

7. Geiser, T. et al. Targeting Arousal and Sleep through Noninvasive Brain Stimulation to Improve Mental Health. Neuropsychobiology 79, 284–292 (2020).

8. Grandner, M. A. Epidemiology of insufficient sleep and poor sleep quality. in Sleep and health 11– 20 (Academic Press, 2019).

9. Malkani, R. G. & Zee, P. C. Brain Stimulation for Improving Sleep and Memory. Sleep Med Clin 17, 505–521 (2022).

10. Dondé, C. et al. The Effects of Transcranial Electrical Stimulation of the Brain on Sleep: A Systematic Review. Front Psychiatry 12, 646569 (2021).

11. Grimaldi, D., Papalambros, N. A., Zee, P. C. & Malkani, R. G. Neurostimulation techniques to enhance sleep and improve cognition in aging. Neurobiol Dis 141, 104865 (2020).

12. Sonmez, A. I. et al. Accelerated TMS for Depression: A systematic review and meta-analysis. Psychiatry Res 273, 770–781 (2019).

13. Nuninga, J. O. et al. Immediate and long-term effects of bilateral electroconvulsive therapy on cognitive functioning in patients with a depressive disorder. J Affect Disord 238, 659–665 (2018).

14. Lim, A. S. et al. Selective enhancement of rapid eye movement sleep by deep brain stimulation of the human pons. Ann Neurol 66, 110–114 (2009).

15. Arnulf, I. et al. Sleep induced by stimulation in the human pedunculopontine nucleus area. Ann Neurol 67, 546–549 (2010).

16. Nishida, N. et al. Subthalamic nucleus deep brain stimulation restores normal rapid eye movement sleep in Parkinson’s disease. Mov Disord 26, 2418–2422 (2011).

17. Effects of deep brain stimulation of the subthalamic nucleus on sleep architecture in parkinsonian patients. Sleep Medicine 5, 207–210 (2004).

18. Legon, W. et al. Transcranial focused ultrasound modulates the activity of primary somatosensory cortex in humans. Nat Neurosci 17, 322–329 (2014).

19. Attali, D. et al. Deep transcranial ultrasound stimulation using personalized acoustic metamaterials improves treatment-resistant depression in humans. Brain Stimul. 18, 1004–1014 (2025).

20. Darmani, G. et al. Individualized non-invasive deep brain stimulation of the basal ganglia using transcranial ultrasound stimulation. Nat. Commun. 16, 2693 (2025).

21. Chou, T. et al. Transcranial focused ultrasound of the amygdala modulates fear network activation and connectivity. Brain Stimul. 17, 312–320 (2024).

22. Sarica, C., Darmani, G., Ramezanpour, H., Chen, R. & Lozano, A. M. Subthalamic local field potential dynamics during motor cortex and basal ganglia transcranial ultrasound stimulation. Brain Stimul. 18, 539 (2025).

23. Tang, K. W. K. et al. Bioadhesive hydrogel-coupled and miniaturized ultrasound transducer system for long-term, wearable neuromodulation. Nature Communications 16, 1–17 (2025).

24. Acharya, U. R. et al. A Novel Depression Diagnosis Index Using Nonlinear Features in EEG Signals. Eur Neurol 74, 79–83 (2015).

25. Hsieh, J.-C. et al. Design of hydrogel-based wearable EEG electrodes for medical applications. J Mater Chem B 10, 7260–7280 (2022).

26. A highly stable electrode with low electrode-skin impedance for wearable brain-computer interface. Biosensors and Bioelectronics 218, 114756 (2022).

27. Hsieh, J.-C. et al. Design of an injectable, self-adhesive, and highly stable hydrogel electrode for sleep recording. Device 2, (2024).

28. Yuk, H., Lu, B. & Zhao, X. Hydrogel bioelectronics. Chem Soc Rev 48, 1642–1667 (2019).

29. Andrews, L., Keller, S. S., Bhojak, M., Osman-Farah, J. & Macerollo, A. Investigating the neural correlates of subjective sleep changes following subthalamic nucleus deep brain stimulation for Parkinson’s disease. Parkinsonism Relat. Disord. 136, 107887 (2025).

30. Pellow, C., Pichardo, S. & Pike, G. B. A systematic review of preclinical and clinical transcranial ultrasound neuromodulation and opportunities for functional connectomics. Brain Stimul. 17, 734– 751 (2024).

31. Liu, S., Rao, Y., Jang, H., Tan, P. & Lu, N. Strategies for body-conformable electronics. Matter 5, 1104–1136 (2022).

32. Yang, Z., Ma, Z., Liu, S. & Li, J. Tissue adhesion with tough hydrogels: Experiments and modeling. Mech. Mater. 157, 103800 (2021).

33. Xiong, Y. et al. A review of the properties and applications of bioadhesive hydrogels. Polym. Chem. 12, 3721–3739 (2021).

34. Jeong, J.-W. et al. Materials and optimized designs for human-machine interfaces via epidermal electronics. Adv. Mater. 25, 6839–6846 (2013).

35. Jeong, S. H., Zhang, S., Hjort, K., Hilborn, J. & Wu, Z. PDMS-based elastomer tuned soft, stretchable, and sticky for epidermal electronics. Adv. Mater. 28, 5830–5836 (2016).

36. Nishikawa, T., Yamane, H., Matsuhisa, N. & Miki, N. Stretchable strain sensor with small but sufficient adhesion to skin. Sensors (Basel) 23, (2023).

37. Xue, H. et al. Hydrogel electrodes with conductive and substrate-adhesive layers for noninvasive long-term EEG acquisition. Microsyst. Nanoeng. 9, 79 (2023).

38. Campbell, I. G. EEG recording and analysis for sleep research. Curr. Protoc. Neurosci. **Chapter** 10, Unit10.2 (2009).

39. Jae Lee, H., Zhang, S., Meyer, R. J., Sherlock, N. P. & Shrout, T. R. Characterization of piezoelectric ceramics and 1-3 composites for high power transducers. Appl. Phys. Lett. 101, 032902 (2012).

40. Ates, H. C. et al. End-to-end design of wearable sensors. Nature Reviews Materials 7, 887–907 (2022).

41. Schafer, M. E., Spivak, N. M., Korb, A. S. & Bystritsky, A. Design, Development, and Operation of a Low-Intensity Focused Ultrasound Pulsation (LIFUP) System for Clinical Use. IEEE Trans Ultrason Ferroelectr Freq Control 68, 54–64 (2021).

42. Meng, Y., Hynynen, K. & Lipsman, N. Applications of focused ultrasound in the brain: from thermoablation to drug delivery. Nature Reviews Neurology 17, 7–22 (2020).

43. Legon, W., Bansal, P., Tyshynsky, R., Ai, L. & Mueller, J. K. Transcranial focused ultrasound neuromodulation of the human primary motor cortex. Sci. Rep. 8, 10007 (2018).

44. Neuromodulation with single-element transcranial focused ultrasound in human thalamus. 10.1002/hbm.23981 doi:10.1002/hbm.23981.

45. Barksdale, B. R. et al. Low-intensity transcranial focused ultrasound amygdala neuromodulation: a double-blind sham-controlled target engagement study and unblinded single-arm clinical trial. Molecular Psychiatry 1–15 (2025).

46. Sarti, A. & Luca Lorini, F. Echocardiography for Intensivists. (Springer Science & Business Media, 2012).

47. Li, H. et al. Effects of skull properties on continuous-wave transcranial focused ultrasound transmission. J. Acoust. Soc. Am. 157, 2336–2349 (2025).

48. Mueller, J. K., Ai, L., Bansal, P. & Legon, W. Numerical evaluation of the skull for human neuromodulation with transcranial focused ultrasound. J Neural Eng 14, 066012 (2017).

49. Bjerknes, S., Skogseid, I. M., Hauge, T. J., Dietrichs, E. & Toft, M. Subthalamic deep brain stimulation improves sleep and excessive sweating in Parkinson’s disease. npj Parkinson’s Disease 6, 1–7 (2020).

50. Baumann-Vogel, H. et al. The impact of subthalamic deep brain stimulation on sleep–wake behavior: A prospective electrophysiological study in 50 Parkinson patients. Sleep (2017) doi:10.1093/sleep/zsx033.

51. Romigi, A. et al. Pedunculopontine nucleus stimulation influences REM sleep in Parkinson’s disease. Eur. J. Neurol. 15, e64–5 (2008).

52. Aviles-Olmos, I. et al. Long-term outcome of subthalamic nucleus deep brain stimulation for Parkinson’s disease using an MRI-guided and MRI-verified approach. J Neurol Neurosurg Psychiatry 85, 1419–1425 (2014).

53. Lilleeng, B., Gjerstad, M., Baardsen, R., Dalen, I. & Larsen, J. P. The long-term development of non-motor problems after STN-DBS. Acta Neurol Scand 132, 251–258 (2015).

54. Salatino, J. W., Ludwig, K. A., Kozai, T. D. Y. & Purcell, E. K. Glial responses to implanted electrodes in the brain. Nat Biomed Eng 1, 862–877 (2017).

55. Ultrasound Neuromodulation: A Review of Results, Mechanisms and Safety. Ultrasound in Medicine & Biology 45, 1509–1536 (2019).

56. Soh, C. et al. The human subthalamic nucleus transiently inhibits active attentional processes. Brain 147, 3204–3215 (2024).

57. Ghodrati, M., Alwis, D. S. & Price, N. S. C. Orientation selectivity in rat primary visual cortex emerges earlier with low-contrast and high-luminance stimuli. Eur J Neurosci 44, 2759–2773 (2016).

58. Beard, J. L. Iron status and periodic limb movements of sleep in children: a causal relationship? Sleep medicine vol. 5 89–90 (2004).

59. Chahine, L. M., Amara, A. W. & Videnovic, A. A systematic review of the literature on disorders of sleep and wakefulness in Parkinson’s disease from 2005 to 2015. Sleep Med Rev 35, 33–50 (2017).

60. Benarroch, E. E. Subthalamic nucleus and its connections: Anatomic substrate for the network effects of deep brain stimulation. Neurology 70, 1991–1995 (2008).

61. McConnell, G. C., So, R. Q., Hilliard, J. D., Lopomo, P. & Grill, W. M. Effective deep brain stimulation suppresses low-frequency network oscillations in the basal ganglia by regularizing neural firing patterns. J Neurosci 32, 15657–15668 (2012).

62. Neumann, W.-J., Steiner, L. A. & Milosevic, L. Neurophysiological mechanisms of deep brain stimulation across spatiotemporal resolutions. Brain 146, 4456–4468 (2023).

63. Blumenfeld, Z. et al. Sixty-hertz stimulation improves bradykinesia and amplifies subthalamic low-frequency oscillations. Mov Disord 32, 80–88 (2017).

64. Su, D. et al. Frequency-dependent effects of subthalamic deep brain stimulation on motor symptoms in Parkinson’s disease: a meta-analysis of controlled trials. Scientific Reports 8, 1–9 (2018).

65. Stern, J. M. et al. Safety of focused ultrasound neuromodulation in humans with temporal lobe epilepsy. Brain Stimul. 14, 1022–1031 (2021).

66. Yaakub, S. N. et al. Transcranial focused ultrasound-mediated neurochemical and functional connectivity changes in deep cortical regions in humans. Nat. Commun. 14, 5318 (2023).

67. He, B. J. Scale-free properties of the functional magnetic resonance imaging signal during rest and task. J Neurosci 31, 13786–13795 (2011).

68. Garrett, D. D., Kovacevic, N., McIntosh, A. R. & Grady, C. L. The Modulation of BOLD Variability between Cognitive States Varies by Age and Processing Speed. Cereb Cortex 23, 684–693 (2012).

69. Zhang, C. et al. Resting-state BOLD signal variability is associated with individual differences in metacontrol. Sci. Rep. 12, 18425 (2022).

70. Oswal, A. et al. Deep brain stimulation modulates synchrony within spatially and spectrally distinct resting state networks in Parkinson’s disease. Brain 139, 1482–1496 (2016).

71. Horovitz, S. G. et al. Low frequency BOLD fluctuations during resting wakefulness and light sleep: a simultaneous EEG-fMRI study. Hum Brain Mapp 29, 671–682 (2008).

72. Fransson, P. How default is the default mode of brain function? Further evidence from intrinsic BOLD signal fluctuations. Neuropsychologia 44, 2836–2845 (2006).

73. Fukunaga, M. et al. Large-amplitude, spatially correlated fluctuations in BOLD fMRI signals during extended rest and early sleep stages. Magn Reson Imaging 24, 979–992 (2006).

74. Jezzard, P., Chappell, M. A. & Okell, T. W. Arterial spin labeling for the measurement of cerebral perfusion and angiography. J. Cereb. Blood Flow Metab. 38, 603–626 (2018).

75. Rouaud, T. et al. Reducing the desire for cocaine with subthalamic nucleus deep brain stimulation. Proc. Natl. Acad. Sci. U. S. A. 107, 1196–1200 (2010).

76. Fox, M. D. et al. Resting-state networks link invasive and noninvasive brain stimulation across diverse psychiatric and neurological diseases. Proc. Natl. Acad. Sci. U. S. A. 111, E4367–75 (2014).

77. Spisák, T. et al. Probabilistic TFCE: A generalized combination of cluster size and voxel intensity to increase statistical power. Neuroimage 185, 12–26 (2019).

78. Wichmann, T. & DeLong, M. R. Oscillations in the basal ganglia. Nature vol. 400 621–622 (1999).

79. Bevan, M. D., Hallworth, N. E. & Baufreton, J. GABAergic control of the subthalamic nucleus. Prog Brain Res 160, 173–188 (2007).

80. Goldstein, A. N. & Walker, M. P. The role of sleep in emotional brain function. Annu. Rev. Clin. Psychol. 10, 679–708 (2014).

81. Kim, H.-J. et al. Long-lasting forms of plasticity through patterned ultrasound-induced brainwave entrainment. Sci. Adv. 10, eadk3198 (2024).

82. Clennell, B. et al. Transient ultrasound stimulation has lasting effects on neuronal excitability. Brain Stimul. 14, 217–225 (2021).

83. Laffan, A., Caffo, B., Swihart, B. J. & Punjabi, N. M. Utility of sleep stage transitions in assessing sleep continuity. Sleep 33, 1681–1686 (2010).

84. Bianchi, M. T., Cash, S. S., Mietus, J., Peng, C.-K. & Thomas, R. Obstructive sleep apnea alters sleep stage transition dynamics. PLoS One 5, e11356 (2010).

85. Schlemmer, A., Parlitz, U., Luther, S., Wessel, N. & Penzel, T. Changes of sleep-stage transitions due to ageing and sleep disorder. Philos. Trans. A Math. Phys. Eng. Sci. 373, 20140093 (2015).

86. Wei, Y. et al. Sleep stage transition dynamics reveal specific stage 2 vulnerability in insomnia. Sleep 40, (2017).

87. Di Marco, T. et al. Hyperarousal features in the sleep architecture of individuals with and without insomnia. J. Sleep Res. 34, e14256 (2025).

88. Sun, H. et al. Large-scale automated sleep staging. Sleep 40, (2017).

89. Hariri, A. R., Tessitore, A., Mattay, V. S., Fera, F. & Weinberger, D. R. The amygdala response to emotional stimuli: a comparison of faces and scenes. Neuroimage 17, 317–323 (2002).

90. Davis, M. & Whalen, P. J. The amygdala: vigilance and emotion. Mol. Psychiatry 6, 13–34 (2001).

91. Baglioni, C. et al. Insomnia disorder is associated with increased amygdala reactivity to insomnia-related stimuli. Sleep 37, 1907–1917 (2014).

92. Walker, H. C. et al. Activation of subthalamic neurons by contralateral subthalamic deep brain stimulation in Parkinson disease. J. Neurophysiol. 105, 1112–1121 (2011).

93. Farokhniaee, A., Marceglia, S., Priori, A. & Lowery, M. M. Effects of contralateral deep brain stimulation and levodopa on subthalamic nucleus oscillatory activity and phase-amplitude coupling. Neuromodulation 26, 310–319 (2023).

94. Hasegawa, H. et al. The effect of unilateral subthalamic nucleus deep brain stimulation on contralateral subthalamic nucleus local field potentials. Neuromodulation 23, 509–514 (2020).

95. Makris, N. et al. Decreased volume of left and total anterior insular lobule in schizophrenia. Schizophr. Res. 83, 155–171 (2006).

96. Frazier, J. A. et al. Structural brain magnetic resonance imaging of limbic and thalamic volumes in pediatric bipolar disorder. Am. J. Psychiatry 162, 1256–1265 (2005).

97. Desikan, R. S. et al. An automated labeling system for subdividing the human cerebral cortex on MRI scans into gyral based regions of interest. Neuroimage 31, 968–980 (2006).

98. Goldstein, J. M. et al. Hypothalamic abnormalities in schizophrenia: sex effects and genetic vulnerability. Biol. Psychiatry 61, 935–945 (2007).

99. Van Someren, E. J. W. Brain mechanisms of insomnia: new perspectives on causes and consequences. Physiol. Rev. 101, 995–1046 (2021).

100. Buysse, D. J., Germain, A., Hall, M., Monk, T. H. & Nofzinger, E. A. A neurobiological model of insomnia. Drug Discov. Today Dis. Models 8, 129–137 (2011).

101. Gong, L. et al. Amygdala changes in chronic insomnia and their association with sleep and anxiety symptoms: Insight from shape analysis. Neural Plast. 2019, 8549237 (2019).

102. Nofzinger, E. A. et al. Functional neuroimaging evidence for hyperarousal in insomnia. Am. J. Psychiatry 161, 2126–2128 (2004).

103. Prasad, A. A. & Wallén-Mackenzie, Å. Architecture of the subthalamic nucleus. Commun. Biol. 7, 78 (2024).

104. Poldrack, R. A. Region of interest analysis for fMRI. Soc. Cogn. Affect. Neurosci. 2, 67–70 (2007).

105. Kalmbach, D. A., Anderson, J. R. & Drake, C. L. The impact of stress on sleep: Pathogenic sleep reactivity as a vulnerability to insomnia and circadian disorders. J. Sleep Res. 27, e12710 (2018).

106. Ellis, B. J., Jackson, J. J. & Boyce, W. T. The stress response systems: Universality and adaptive individual differences. Developmental Review 26, 175–212 (2006).

107. Pacák, K. & Palkovits, M. Stressor specificity of central neuroendocrine responses: implications for stress-related disorders. Endocr. Rev. 22, 502–548 (2001).

108. Rotenberg, S. & McGrath, J. J. Inter-relation between autonomic and HPA axis activity in children and adolescents. Biol. Psychol. 117, 16–25 (2016).

109. Kim, H.-G., Cheon, E.-J., Bai, D.-S., Lee, Y. H. & Koo, B.-H. Stress and heart rate variability: A meta-analysis and review of the literature. Psychiatry Investig. 15, 235–245 (2018).

110. Maes, J. et al. Sleep misperception, EEG characteristics and autonomic nervous system activity in primary insomnia: a retrospective study on polysomnographic data. Int. J. Psychophysiol. 91, 163– 171 (2014).

111. Monroe, L. J. Psychological and physiological differences between good and poor sleepers. J. Abnorm. Psychol. 72, 255–264 (1967).

112. Varkevisser, M., Van Dongen, H. P. A. & Kerkhof, G. A. Physiologic indexes in chronic insomnia during a constant routine: evidence for general hyperarousal? Sleep 28, 1588–1596 (2005).

113. Johann, A. F. et al. Cognitive behavioural therapy for insomnia does not appear to have a substantial impact on early markers of cardiovascular disease: A preliminary randomized controlled trial. J. Sleep Res. 29, e13102 (2020).

114. Huang, Y. et al. Insomnia and impacts on facial expression recognition accuracy, intensity and speed: A meta-analysis. J. Psychiatr. Res. 160, 248–257 (2023).

115. Motomura, Y. et al. Sleep debt elicits negative emotional reaction through diminished amygdala-anterior cingulate functional connectivity. PLoS One 8, e56578 (2013).

116. Krause, A. J. et al. The sleep-deprived human brain. Nat. Rev. Neurosci. 18, 404–418 (2017).

117. Lazarus, M., Chen, J.-F., Urade, Y. & Huang, Z.-L. Role of the basal ganglia in the control of sleep and wakefulness. Curr. Opin. Neurobiol. 23, 780–785 (2013).

118. Taber, K. H. & Hurley, R. A. Functional neuroanatomy of sleep and sleep deprivation. J. Neuropsychiatry Clin. Neurosci. 18, 1–5 (2006).

119. Bishir, M. et al. Sleep deprivation and neurological disorders. Biomed Res. Int. 2020, 5764017 (2020).

120. Wu, J. C. et al. Frontal lobe metabolic decreases with sleep deprivation not totally reversed by recovery sleep. Neuropsychopharmacology 31, 2783–2792 (2006).

121. Lazarus, M., Huang, Z.-L., Lu, J., Urade, Y. & Chen, J.-F. How do the basal ganglia regulate sleep-wake behavior? Trends Neurosci. 35, 723–732 (2012).

122. Cole, E. J. et al. Stanford neuromodulation therapy (SNT): A double-blind randomized controlled trial. Am. J. Psychiatry 179, 132–141 (2022).

123. Nakajima, K. et al. A causal role of anterior prefrontal-putamen circuit for response inhibition revealed by transcranial ultrasound stimulation in humans. Cell Rep. 40, 111197 (2022).

124. Mehta, D. D. et al. A systematic review and meta-analysis of neuromodulation therapies for substance use disorders. Neuropsychopharmacology 49, 649–680 (2024).

125. Fan, J. M. et al. Thalamic transcranial ultrasound stimulation in treatment resistant depression. Brain Stimul. 17, 1001–1004 (2024).

126. Matt, E. et al. Ultrasound neuromodulation with transcranial pulse stimulation in Alzheimer disease: A randomized clinical trial. JAMA Netw. Open 8, e2459170 (2025).

127. Matt, E., Radjenovic, S., Mitterwallner, M. & Beisteiner, R. Current state of clinical ultrasound neuromodulation. Front. Neurosci. 18, 1420255 (2024).

128. Visvanathan, V. & Morrissey, M. S. C. Anatomical variations of the temporal bone on high-resolution computed tomography imaging: how common are they? J. Laryngol. Otol. 129, 634–637 (2015).

129. Martin, E. et al. Ultrasound system for precise neuromodulation of human deep brain circuits. Nat. Commun. 16, 8024 (2025).

130. Beisteiner, R., Hallett, M. & Lozano, A. M. Ultrasound neuromodulation as a new brain therapy. Adv. Sci. (Weinh.) 10, e2205634 (2023).

131. Seo, H. et al. Numerical investigation of layered homogeneous skull model for simulations of transcranial focused ultrasound. Neuromodulation 28, 103–114 (2025).

132. Daneshzand, M. et al. Model-based navigation of transcranial focused ultrasound neuromodulation in humans: Application to targeting the amygdala and thalamus. Brain Stimul. 17, 958–969 (2024).

133. Samoudi, M. A., Van Renterghem, T. & Botteldooren, D. Computational modeling of a single-element transcranial focused ultrasound transducer for subthalamic nucleus stimulation. J Neural Eng 16, 026015 (2019).

134. Fjield, T., Fan, X. & Hynynen, K. A parametric study of the concentric-ring transducer design for MRI guided ultrasound surgery. J Acoust Soc Am 100, 1220–1230 (1996).

135. Creton, C. & Ciccotti, M. Fracture and adhesion of soft materials: a review. Rep Prog Phys 79, 046601 (2016).

136. Kendall, K. Thin-film peeling-the elastic term. J. Phys. D Appl. Phys. 8, 1449–1452 (1975).

137. Pennes, H. H. Analysis of tissue and arterial blood temperatures in the resting human forearm. 1948. J Appl Physiol (1985) 85, 5–34 (1998).

138. Sudhyadhom, A., Haq, I. U., Foote, K. D., Okun, M. S. & Bova, F. J. A high resolution and high contrast MRI for differentiation of subcortical structures for DBS targeting: the Fast Gray Matter Acquisition T1 Inversion Recovery (FGATIR). Neuroimage 47 Suppl 2, T44–52 (2009).

139. MacLean, A. W., Fekken, G. C., Saskin, P. & Knowles, J. B. Psychometric evaluation of the Stanford Sleepiness Scale. J Sleep Res 1, 35–39 (1992).

140. Subjective and Objective Assessment of Hypersomnolence. Sleep Medicine Clinics 12, 313–322 (2017).

141. Johns, M. W. A new method for measuring daytime sleepiness: the Epworth sleepiness scale. Sleep 14, 540–545 (1991).

142. Julian, L. J. Measures of anxiety: State-Trait Anxiety Inventory (STAI), Beck Anxiety Inventory (BAI), and Hospital Anxiety and Depression Scale-Anxiety (HADS-A). Arthritis Care Res (Hoboken) 63 Suppl 11, S467–72 (2011).

143. Oostenveld, R. & Praamstra, P. The five percent electrode system for high-resolution EEG and ERP measurements. Clin Neurophysiol 112, 713–719 (2001).

144. Kawada, T. & Suzuki, S. Sleep induction effects of steady 60 dB (A) pink noise. Ind Health 31, 35– 38 (1993).

145. Messineo, L. et al. Broadband Sound Administration Improves Sleep Onset Latency in Healthy Subjects in a Model of Transient Insomnia. Front. Neurol. 8, 288775 (2017).

146. Capezuti, E. et al. Systematic review: auditory stimulation and sleep. Journal of Clinical Sleep Medicine (2022) doi:10.5664/jcsm.9860.

147. The influence of white noise on sleep in subjects exposed to ICU noise. Sleep Medicine 6, 423–428 (2005).

148. Afshar, P. F., Bahramnezhad, F., Asgari, P. & Shiri, M. Effect of White Noise on Sleep in Patients Admitted to a Coronary Care. Journal of Caring Sciences 5, 103–109 (2016).

149. State transitions between wake and sleep, and within the ultradian cycle, with focus on the link to neuronal activity. Sleep Medicine Reviews 8, 473–485 (2004).

150. Ultrashort sleep-waking schedule. I. Evidence of ultradian rhythmicity in ‘sleepability.’ Electroencephalography and Clinical Neurophysiology 52, 163–174 (1981).

151. RMSSD, a measure of vagus-mediated heart rate variability, is associated with risk factors for SUDEP: The SUDEP-7 Inventory. Epilepsy & Behavior 19, 78–81 (2010).

152. Shaffer, F. & Ginsberg, J. P. An Overview of Heart Rate Variability Metrics and Norms. Front. Public Health 5, 290215 (2017).

154. Cortical functional connectivity indexes arousal state during sleep and anesthesia. NeuroImage 211, 116627 (2020).

153. McHugh, M. L. The Chi-square test of independence. Biochem Med 23, 143–149 (2013).

155. Iglesias, T. L., Boal, J. G., Frank, M. G., Zeil, J. & Hanlon, R. T. Cyclic nature of the REM sleep-like state in the cuttlefish Sepia officinalis. J Exp Biol 222, jeb174862 (2019).

156. Woolrich, M. W. et al. Bayesian analysis of neuroimaging data in FSL. Neuroimage 45, S173–86 (2009).

157. Chen, G., Saad, Z. S., Britton, J. C., Pine, D. S. & Cox, R. W. Linear mixed-effects modeling approach to FMRI group analysis. Neuroimage 73, 176–190 (2013).

158. Pauli, W. M., Nili, A. N. & Tyszka, J. M. A high-resolution probabilistic in vivo atlas of human subcortical brain nuclei. Scientific Data 5, 1–13 (2018).

159. Grady, C. L. & Garrett, D. D. Understanding variability in the BOLD signal and why it matters for aging. Brain Imaging Behav 8, 274–283 (2014).

160. An improved approach to detection of amplitude of low-frequency fluctuation (ALFF) for resting-state fMRI: Fractional ALFF. Journal of Neuroscience Methods 172, 137–141 (2008).

161. Biswal, B., Yetkin, F. Z., Haughton, V. M. & Hyde, J. S. Functional connectivity in the motor cortex of resting human brain using echo-planar MRI. Magn Reson Med 34, 537–541 (1995).

162. Lowe, M. J., Mock, B. J. & Sorenson, J. A. Functional connectivity in single and multislice echoplanar imaging using resting-state fluctuations. Neuroimage 7, 119–132 (1998).

163. Zang, Y.-F. et al. Altered baseline brain activity in children with ADHD revealed by resting-state functional MRI. Brain Dev 29, 83–91 (2007).

164. Smith, S. M. Fast robust automated brain extraction. Hum Brain Mapp 17, 143–155 (2002).

165. Alsop, D. C. et al. Recommended implementation of arterial spin-labeled perfusion MRI for clinical applications: A consensus of the ISMRM perfusion study group and the European consortium for ASL in dementia. Magnetic Resonance in Medicine 73, 102–116 (2015).

166. Fonzo, G. A. et al. Cognitive-behavioral therapy for generalized anxiety disorder is associated with attenuation of limbic activation to threat-related facial emotions. J Affect Disord 169, 76–85 (2014).

167. Fonzo, G. A. et al. Common and disorder-specific neural responses to emotional faces in generalised anxiety, social anxiety and panic disorders. Br J Psychiatry 206, 206–215 (2015).

168. Tottenham, N. et al. The NimStim set of facial expressions: judgments from untrained research participants. Psychiatry Res 168, 242–249 (2009).

169. Peirce, J. et al. PsychoPy2: Experiments in behavior made easy. Behav Res Methods 51, 195–203 (2019).

