## Supplementary Information for "Wearable Focused Ultrasound Neuromodulation and Electrophysiological Recording Patch for REM Sleep Enhancement"

**Supplementary Table 1. Phase control parameters of CRUTA.**

| Delay Variable | Channel Time Delay (μs) | | | | | | | |
|---|---|---|---|---|---|---|---|---|
| | 1 | 2 | 3 | 4 | 5 | 6 | 7 | 8 |
| 50 | 3.41251506 | 3.21489905 | 2.91713277 | 2.52042039 | 2.02777903 | 1.44043567 | 0.76370241 | 0 |
| 60 | 2.8737777 | 2.70889273 | 2.46007496 | 2.12789311 | 1.71431743 | 1.21972224 | 0.64785615 | 0 |
| 70 | 2.47926528 | 2.33782904 | 2.12420323 | 1.83864537 | 1.48255151 | 1.05588634 | 0.56147943 | 0 |
| 80 | 2.17864985 | 2.05483266 | 1.8677085 | 1.61737021 | 1.30486884 | 0.92996337 | 0.4948993 | 0 |
| 90 | 1.9423205 | 1.83222387 | 1.6657681 | 1.44295417 | 1.16461072 | 0.83039216 | 0.44214872 | 0 |
| 100 | 1.7518221 | 1.65271136 | 1.5028217 | 1.30210118 | 1.05122677 | 0.74980136 | 0.39939385 | 0 |
| 110 | 1.59509505 | 1.50497837 | 1.36866152 | 1.18606156 | 0.95774725 | 0.68329893 | 0.36407685 | 0 |
| 120 | 1.46394362 | 1.3813255 | 1.25633103 | 1.0888592 | 0.87939861 | 0.62752329 | 0.33443344 | 0 |
| 130 | 1.3526117 | 1.2763408 | 1.16093443 | 1.00628135 | 0.81280893 | 0.5800941 | 0.30921082 | 0 |
| 140 | 1.25694214 | 1.18611325 | 1.07893039 | 0.93527697 | 0.75553236 | 0.53928156 | 0.2874966 | 0 |
| 150 | 1.17385986 | 1.10774844 | 1.00769618 | 0.87358425 | 0.70575351 | 0.50379986 | 0.26861146 | 0 |
| 160 | 1.10104252 | 1.03905965 | 0.94524921 | 0.81949208 | 0.66209767 | 0.47267425 | 0.2520397 | 0 |
| 170 | 1.03670423 | 0.97836474 | 0.89006363 | 0.77168276 | 0.6235054 | 0.44515276 | 0.23738307 | 0 |
| 180 | 0.97944952 | 0.92434901 | 0.84094643 | 0.72912548 | 0.58914737 | 0.42064634 | 0.22432933 | 0 |
| 190 | 0.92817246 | 0.87597028 | 0.79695164 | 0.69100252 | 0.55836534 | 0.39868716 | 0.21263031 | 0 |
| 200 | 0.88198543 | 0.83239199 | 0.75731968 | 0.65665708 | 0.53063038 | 0.37889906 | 0.20208634 | 0 |

**Supplementary Table 2. Participant demographics - STN-FUS with CRUTA.**

| Subject ID | Age | Ethnicity | Sex | PSQI | STN Distance (mm) |
|---|---|---|---|---|---|
| 1 | 28 | White | Female | 8 | 76.5 |
| 2 | 19 | White | Male | 6 | 74.9 |
| 3 | 20 | Asian | Female | 5 | 64.2 |
| 4 | 37 | White | Male | 3 | 74.1 |
| 5 | 24 | White | Female | 5 | 69.5 |
| 6 | 31 | Asian | Female | 1 | 71.5 |
| 7 | 19 | Asian | Female | 5 | 69.5 |
| 8 | 38 | Asian | Male | 7 | 71.4 |
| 9 | 33 | Black or African American | Male | 4 | 74.6 |
| 10 | 19 | Asian | Female | 3 | 76.5 |
| 11 | 19 | Prefer not to say | Male | 3 | 73.8 |
| 12 | 25 | Asian | Female | 4 | 73.9 |
| 13 | 24 | Asian | Male | 9 | 77.2 |
| 14 | 26 | White | Male | 7 | 71.9 |
| 15 | 23 | Asian | Female | 2 | 74.1 |
| 16 | 25 | Asian | Male | 7 | 73.1 |

**Supplementary Table 3. Participant demographics - NEUSLeeP on healthy group.**

| Subject ID | Age | Ethnicity | Sex | PSQI | STN Distance (mm) |
|---|---|---|---|---|---|
| 1 | 19 | White | Female | 2 | 76.1 |
| 2 | 19 | Asian | Male | 4 | 77.4 |
| 3 | 22 | Prefer not to say | Male | 2 | 73.1 |
| 4 | 20 | White | Female | 2 | 68.4 |
| 5 | 19 | White | Female | 2 | 64.4 |
| 6 | 33 | Asian | Male | 2 | 72.6 |
| 7 | 20 | White | Male | 3 | 70.7 |
| 8 | 25 | White | Female | 5 | 69.3 |
| 9 | 37 | White | Male | 2 | 72.8 |
| 10 | 25 | Asian | Female | 3 | 73.3 |
| 11 | 20 | Native Hawaiian or Other Pacific Islander | Male | 4 | 77.1 |
| 12 | 20 | Black or African American | Male | 2 | 78.1 |
| 13 | 18 | White | Female | 3 | 72.6 |
| 14 | 19 | White | Female | 3 | 79.3 |
| 15 | 19 | White | Male | 5 | 71.4 |
| 16 | 24 | Asian | Female | 2 | 76.1 |

**Supplementary Table 4. Participant demographics - NEUSLeeP on insomnia group.**

| Subject ID | Age | Ethnicity | Sex | PSQI | STN Distance (mm) |
|---|---|---|---|---|---|
| 17 | 18 | Asian | Female | 10 | 68.8 |
| 18 | 24 | Asian | Male | 5 | 69.1 |
| 19 | 25 | Asian | Female | 9 | 74.2 |
| 20 | 28 | White | Female | 7 | 62.7 |
| 21 | 27 | Asian | Female | 8 | 66.6 |
| 22 | 38 | Asian | Male | 7 | 69.3 |
| 23 | 18 | Asian | Female | 6 | 65.1 |
| 24 | 26 | White | Male | 7 | 72.8 |
| 25 | 33 | Asian | Male | 10 | 82.3 |
| 26 | 27 | Asian | Male | 9 | 73.6 |
| 27 | 21 | Prefer not to say | Male | 7 | 74.1 |
| 28 | 20 | White | Female | 6 | 69.8 |

**Supplementary Table 5. Abbreviation of region-of-interests in brain structures.**

| Abbreviation | Brain Structure | Abbreviation | Brain Structure |
|---|---|---|---|
| **Amyg** | Amygdala | **NAc** | Nucleus Accumbens |
| **AG** | Angular Gyrus | **OFG** | Occipital Fusiform Gyrus |
| **Ca** | Caudate | **OP** | Occipital Pole |
| **COC** | Central Opercular Cortex | **Pal** | Pallidum |
| **CG-a** | Cingulate Gyrus - anterior | **PCG** | Paracingulate Gyrus |
| **CG-p** | Cingulate Gyrus - posterior | **PHG-a** | Parahippocampal Gyrus - anterior |
| **Cun** | Cuneal Cortex | **PHG-p** | Parahippocampal Gyrus - posterior |
| **Exa** | Extrastriate Cortex | **POC** | Parietal Operculum Cortex |
| **FMC** | Frontal Medial Cortex | **PBP** | Parabrachial Pigmented Nucleus |
| **FOC** | Frontal Operculum Cortex | **PPo** | Planum Polare |
| **OFC** | Frontal Orbital Cortex | **PT** | Planum Temporale |
| **FP** | Frontal Pole | **PostCG** | Postcentral Gyrus |
| **GPe** | Globus Pallidus externa | **PreCG** | Precentral Gyrus |
| **GPi** | Globus Pallidus interna | **Precun** | Precuneous Cortex |
| **HG** | Heschls Gyrus | **Pu** | Putamen |
| **Hipp** | Hippocampus | **RN** | Red Nucleus |
| **HN** | Hypothalamus-1 | **SNc** | Substantia Nigra pars compacta |
| **HTH** | Hypothalamus-2 | **SNr** | Substantia Nigra pars reticulata |
| **IFGtri** | Inferior Temporal Gyrus - pars triangularis | **STN** | Subthalamic Nucleus |
| **ITG-a** | Inferior Temporal Gyrus - anterior | **SCC** | Subcallosal Cortex |
| **ITG-p** | Inferior Temporal Gyrus - posterior | **SFG** | Superior Frontal Gyrus |
| **ITG-to** | Inferior Temporal Gyrus - temporoccipital | **SPL** | Superior Parietal Lobule |
| **Ins** | Insular Cortex | **STG-a** | Superior Temporal Gyrus - anterior |
| **ICC** | Intracalcarine Cortex | **STG-p** | Superior Temporal Gyrus - posterior |
| **LOC-i** | Lateral Occipital Cortex - inferior | **SCCx** | Supracalcarine Cortex |
| **LOC-s** | Lateral Occipital Cortex - superior | **SMG-a** | Supramarginal Gyrus - anterior |
| **LG** | Lingual Gyrus | **SMG-p** | Supramarginal Gyrus - posterior |
| **MFG** | Middle Frontal Gyrus | **TOFC** | Temporal Occipital Fusiform Cortex |
| **MTG-a** | Middle Temporal Gyrus - anterior | **TP** | Temporal Pole |
| **MTG-p** | Middle Temporal Gyrus - posterior | **Thal** | Thalamus |
| **MTG-to** | Middle Temporal Gyrus - temporoccipital | **VeP** | Ventral Palldium |
| **MN** | Medial Thalamic Nucleus | **VTA** | Ventral Tegmental Area |

**Supplementary Table 6. Dimensions and parameters of CRUTA.**

| Channel Element | ID (mm) | OD (mm) | Material | Geometry | Electrode Plating |
|---|---|---|---|---|---|
| 1 | 5.35 | 10.07 | DL-47 | Circular Ring | Nickel, both electrodes same side |
| 2 | 10.72 | 15.42 | DL-47 | Circular Ring | Nickel, both electrodes same side |
| 3 | 16.07 | 20.78 | DL-47 | Circular Ring | Nickel, both electrodes same side |
| 4 | 21.43 | 26.14 | DL-47 | Circular Ring | Nickel, both electrodes same side |
| 5 | 26.79 | 31.49 | DL-47 | Circular Ring | Nickel, both electrodes same side |
| 6 | 32.15 | 36.86 | DL-47 | Circular Ring | Nickel, both electrodes same side |
| 7 | 37.51 | 42.21 | DL-47 | Circular Ring | Nickel, both electrodes same side |
| 8 | 42.86 | 47.57 | DL-47 | Circular Ring | Nickel, both electrodes same side |

**Supplementary Table 7. Parameters used in thermal simulations.**

| Parameter | Value | Units | Ref |
|---|---|---|---|
| $CBF_{gray}$ - cerebral blood flow in gray matter | 80 | ml/min/100g | 1 |
| w - skull thickness | 3.42 | mm | - |
| $\rho_t$ - brain tissue density | 1040 | $kg/m^3$ | 2 |
| $\rho_b$ - blood density | 1060 | $kg/m^3$ | 3 |
| $\alpha_{skull}$ - attenuation coefficient of skull | 5 | Np/cm/MHz | 4 |
| $\alpha_{brainl}$ - attenuation coefficient brain | 0.23 | Np/cm/MHz | 4 |
| $c_b$ - specific heat of blood | 3770 | J/kg/°C | 4 |
| $k_t$ - tissue thermal conductivity | 0.195 | W/m/°C | 5 |
| $\omega_t$ - blood perfusion rate | 0.000142 | $kg/m^3/s$ | - |
| $T_a$ - arterial blood temperature | 37 | °C | 4 |
| $Z_{brain}$ - acoustic impedance of brain tissues | 1.5 | MRayl | 6 |
| $Z_{waterl}$ - acoustic impedance of water | 1.48 | MRayl | 6 |
| $Z_{skull}$ - acoustic impedance of skull | 7.8 | MRayl | 6 |

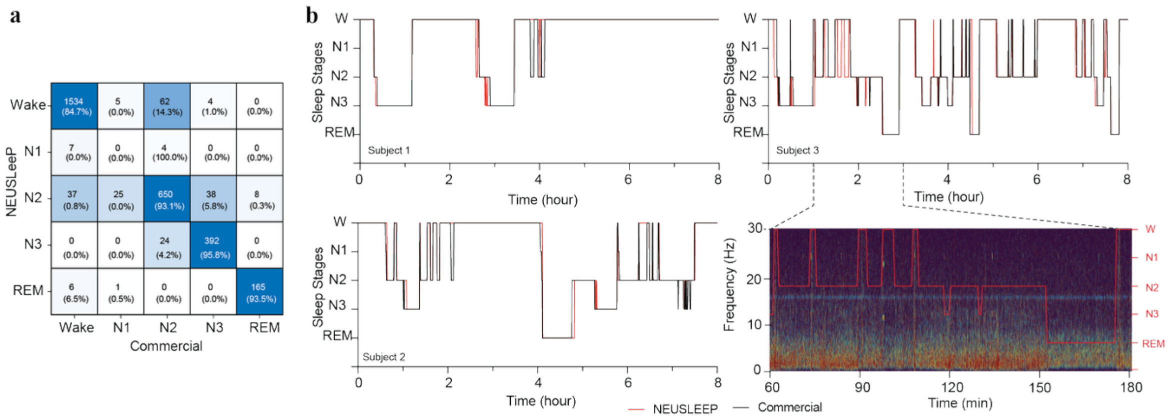

**Supplementary Figure 1.** Sleep recording performance and benchmarking of NEUSLeeP with commercial EEG headcap. **a**) Correlation matrix of hypnogram between NEUSLeeP and commercial EEG headcap (32-channel AntNeuro). **b**) Individual hypnogram comparisons with demonstration of spectrogram under varying sleep stages.

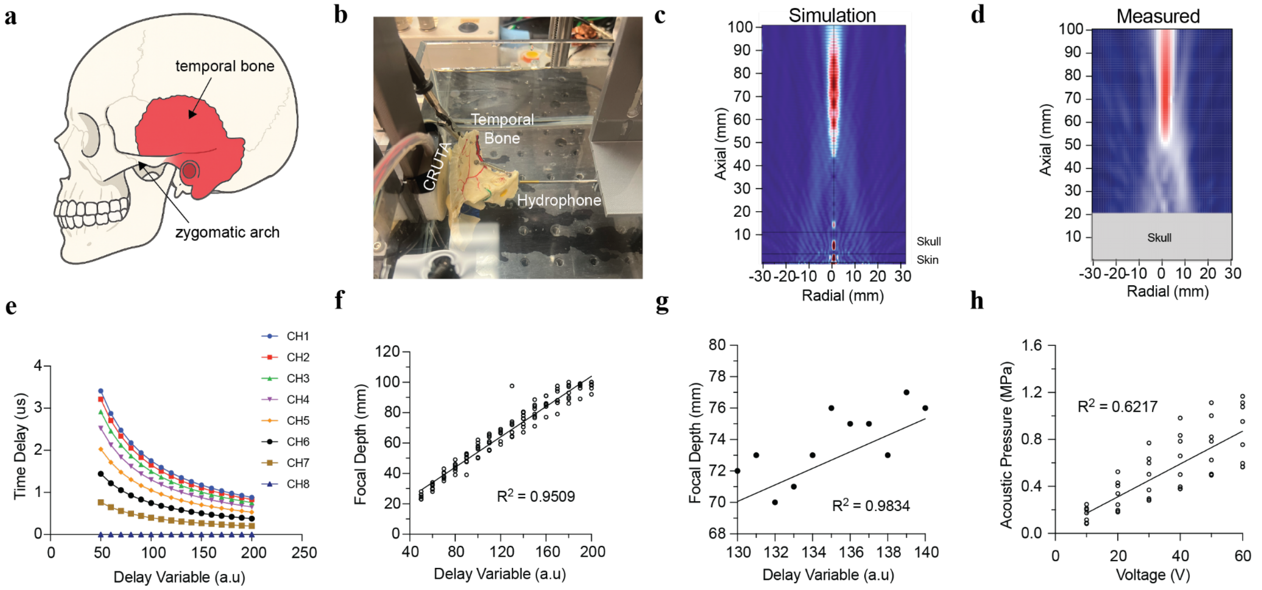

**Supplementary Figure 2. Phase control of CRUTA and characterization. a)** Schematic of temporal bone in human skull and position of zygomatic arch. **b)** Photograph of CRUTA and temporal bone used in experimental setup for acoustic characterization. **c)** Simulation of *in vivo* acoustic field distribution of CRUTA with temporal bone. **d)** Measured *in-vitro* acoustic field distribution of CRUTA with temporal bone. **e)** Time/phase delay of individual elements of CRUTA with respect to a placeholder variable (delay variable) used for determining focal depth. **f)** Linear regression and relationship of focal depth measured across multiple CRUTA (n = 8, $R^2$ = 0.9509). **g)** Zoomed in view at approximal average STN depth from temporal windows for demonstration of spatial resolution. **h)** Linear regression and relationship of acoustic pressure measured across multiple CRUTA (n = 8, $R^2$ = 0.6217) in response to driven voltage.

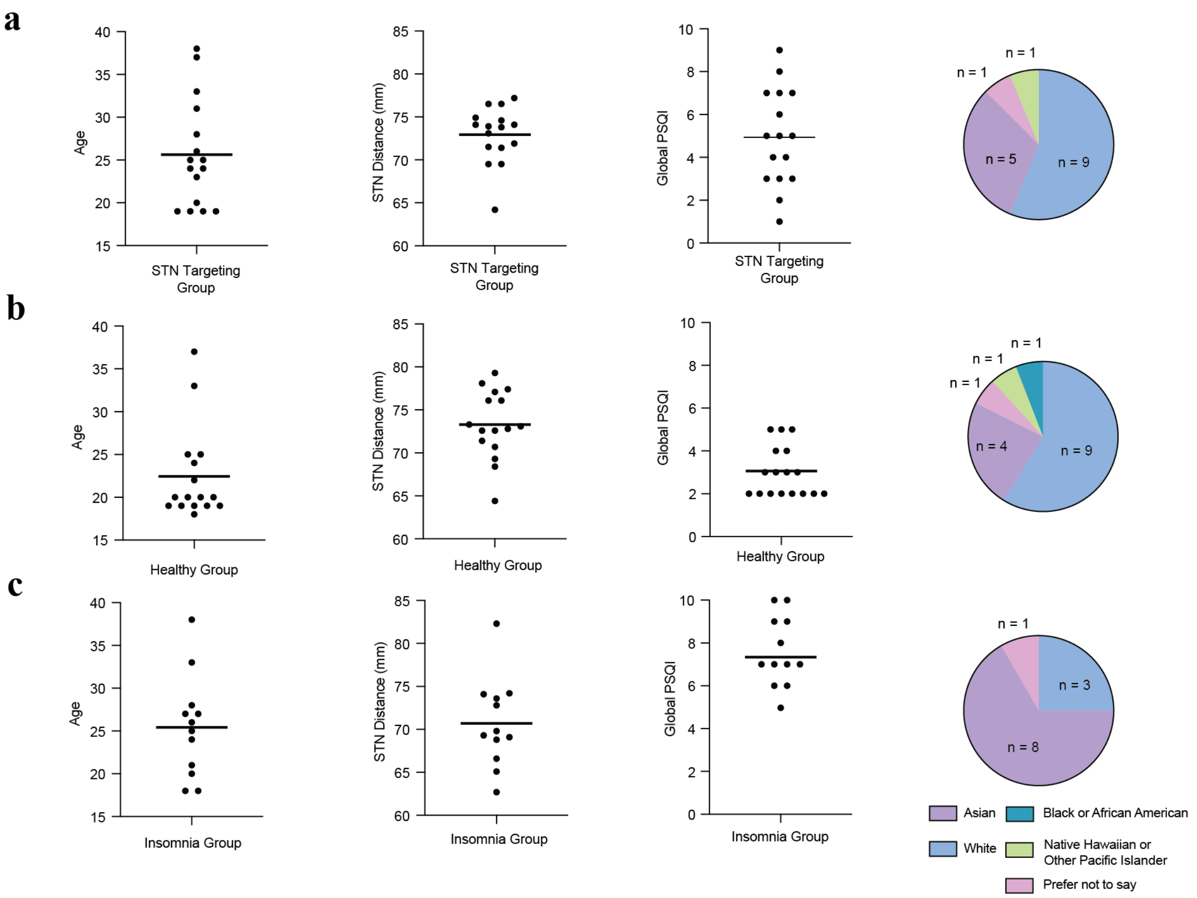

**Supplementary Figure 3. Participant information and study groups. a)** Demographic and subject PSQI scores recruited/enrolled for STN-FUS Target engagement study with CRUTA. **b)** Demographic and subject PSQI scores recruited/enrolled for NEUSLeeP study for healthy group. **c)** Demographic and subject PSQI scores recruited/enrolled for NEUSLeeP study for insomnia groups.

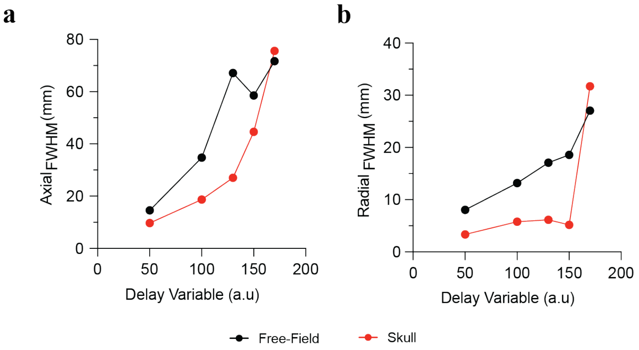

**Supplementary Figure 4. Spatial resolution of CRUTA. a)** Average axial full width half maximum measured at various focal depth dependent on delay variable (n = 8) in free-field and with temporal skull. **b)** Average radial full width half maximum measured at various focal depth dependent on delay variable (n = 8) in free-field and with temporal skull.

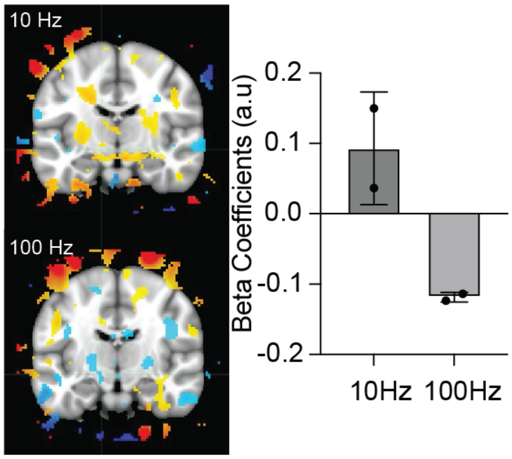

**Supplementary Figure 5. Pilot-study investigation on pulse repetition frequency effects for STN-FUS measured by functional Magnetic Resonance Imaging (fMRI).** Beta coefficient Blood oxygen level-dependent (BOLD) signal response across whole-brain with respect to 10 Hz vs 100 Hz STN-FUS using BrainSonix BX Pulsar 1002 (n = 2).

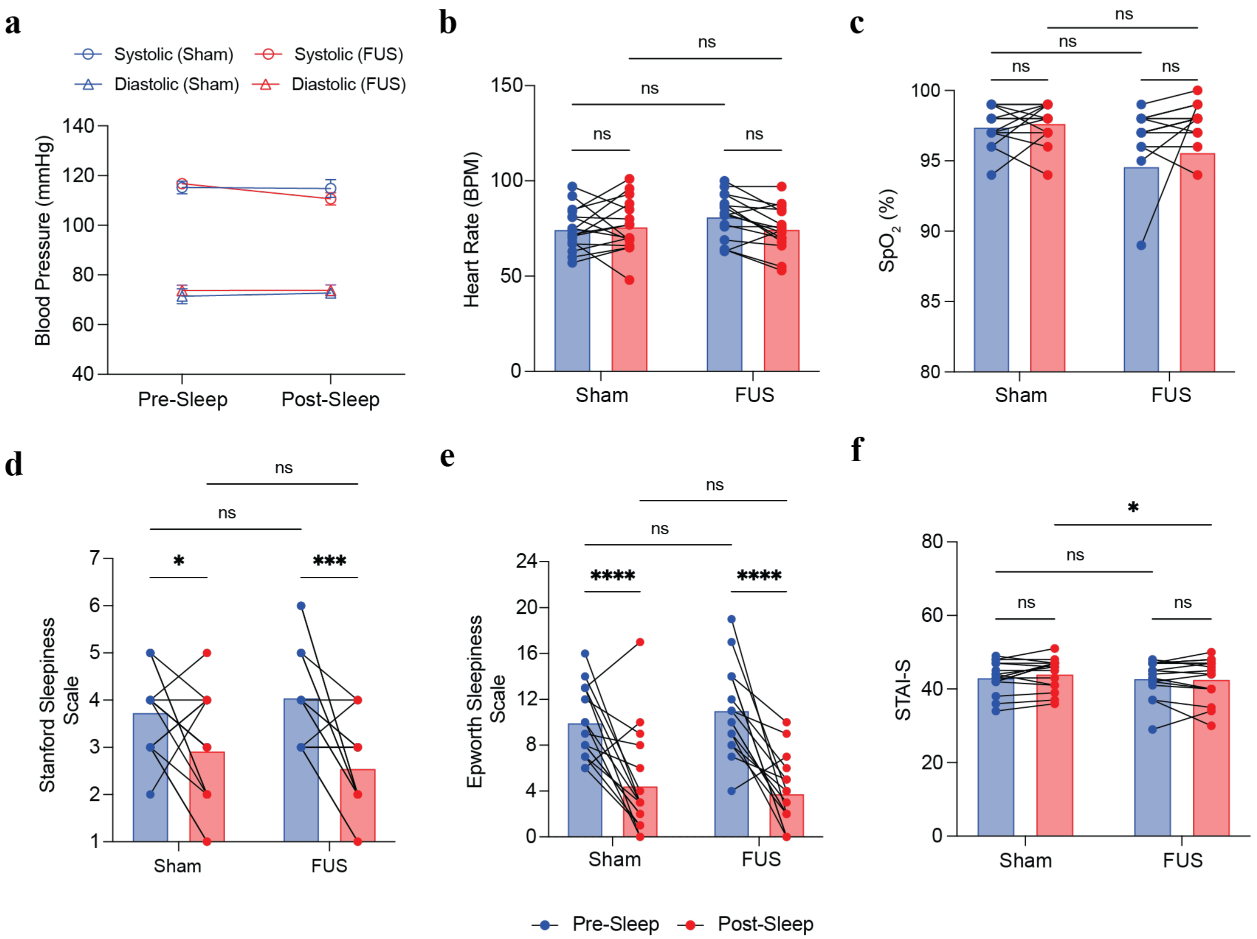

**Supplementary Figure 6. Secondary outcome of NEUSLeeP on healthy group.** NEUSLeeP did not create adverse vital changes in blood pressure, heart, and $SpO_2$. Quantitative results of subject self-report showed improved wakefulness in both sham and FUS nights, with more prominent decrease in Stanford Sleepiness Scale in healthy groups.

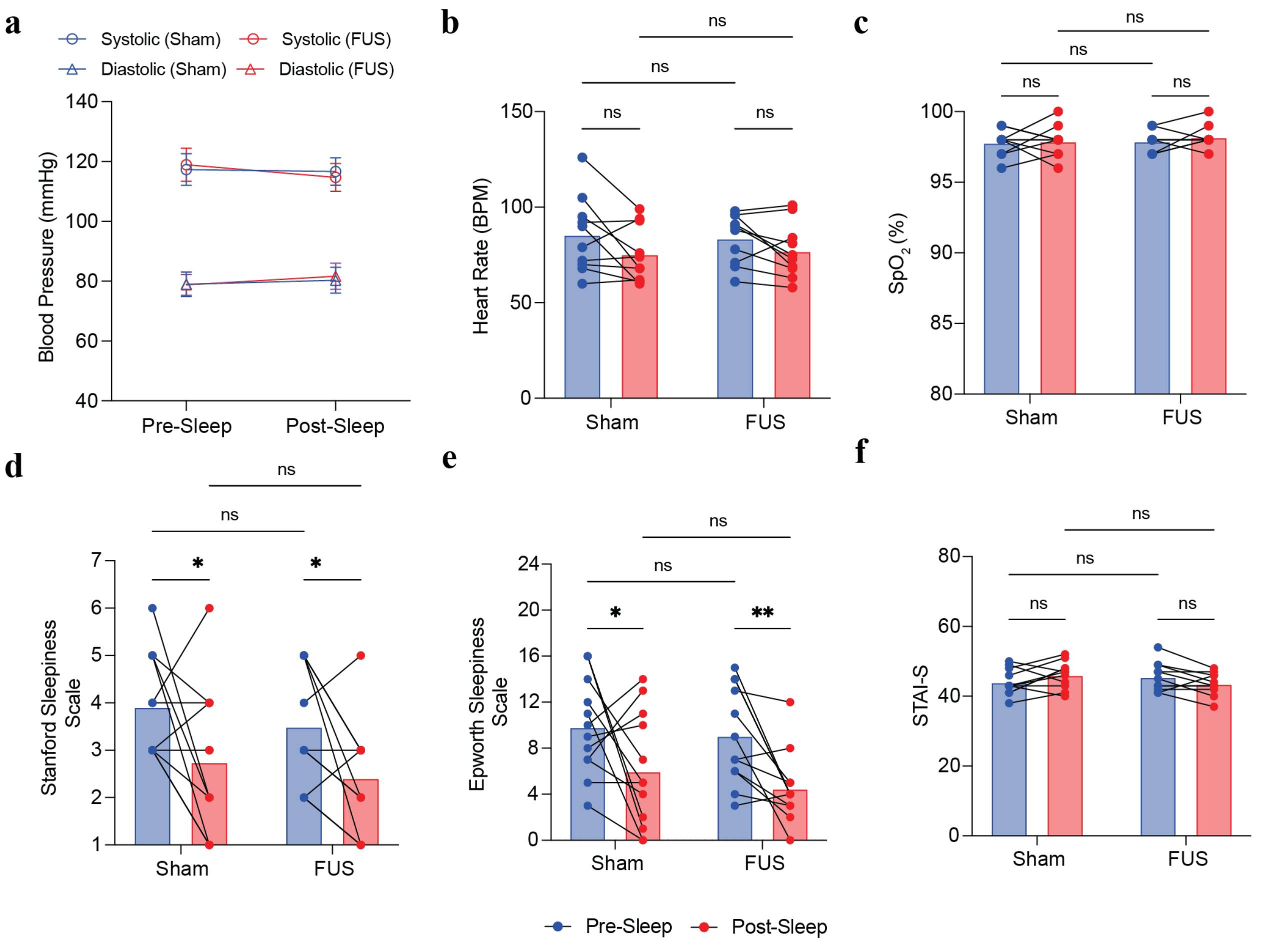

**Supplementary Figure 7. Secondary outcome of NEUSLeeP on insomnia group.** NEUSLeeP did not create adverse vital changes in blood pressure, heart, and $SpO_2$. Quantitative results of subject self-report showed improved wakefulness in both sham and FUS nights, with more prominent decrease in Epsworth Sleepiness Scale in insomnia groups.

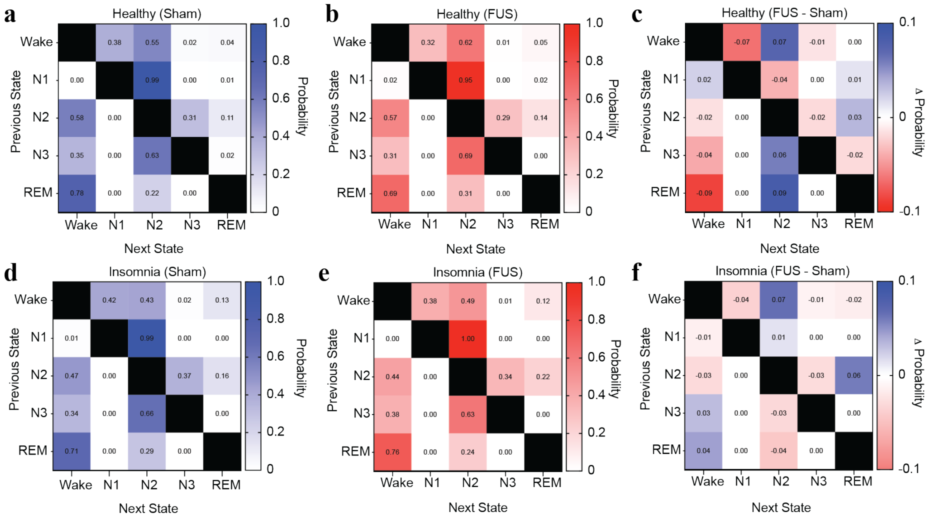

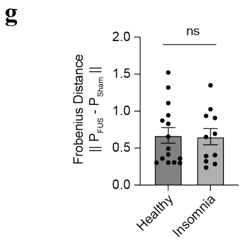

h

| States | $X^2$ (p-value) | | Mantel Test (r) | |
|---|---|---|---|---|
| | Healthy | Insomnia | Healthy | Insomnia |
| Wake | 0.334 | 0.709 | 0.957 | 0.986 |
| N1 | 0.535 | N/A | | |
| N2 | 0.766 | 0.691 | | |
| N3 | 0.663 | 0.997 | | |
| REM | 0.911 | 0.992 | | |

**Supplementary Figure 8. State transition probability matrix. a**) Probability matrix of healthy population during sham condition. **b)** Probability matrix of healthy population during NEUSLeeP-enabled STN-FUS. **c)** Change in probability matrix in healthy populations. **d**) Probability matrix of insomnia population during sham condition. **e)** Probability matrix of insomnia population during NEUSLeeP-enabled STN-FUS. **f)** Change in probability matrix in insomnia populations. **g)** Frobenius distance of state transition matrices between FUS vs. Sham for healthy and insomnia populations. (n = 15 and 11, Two-sided t-test). **h)** Statistical analysis of state transitions (Row-based Chi-squared test and Mantel Spearman Test).

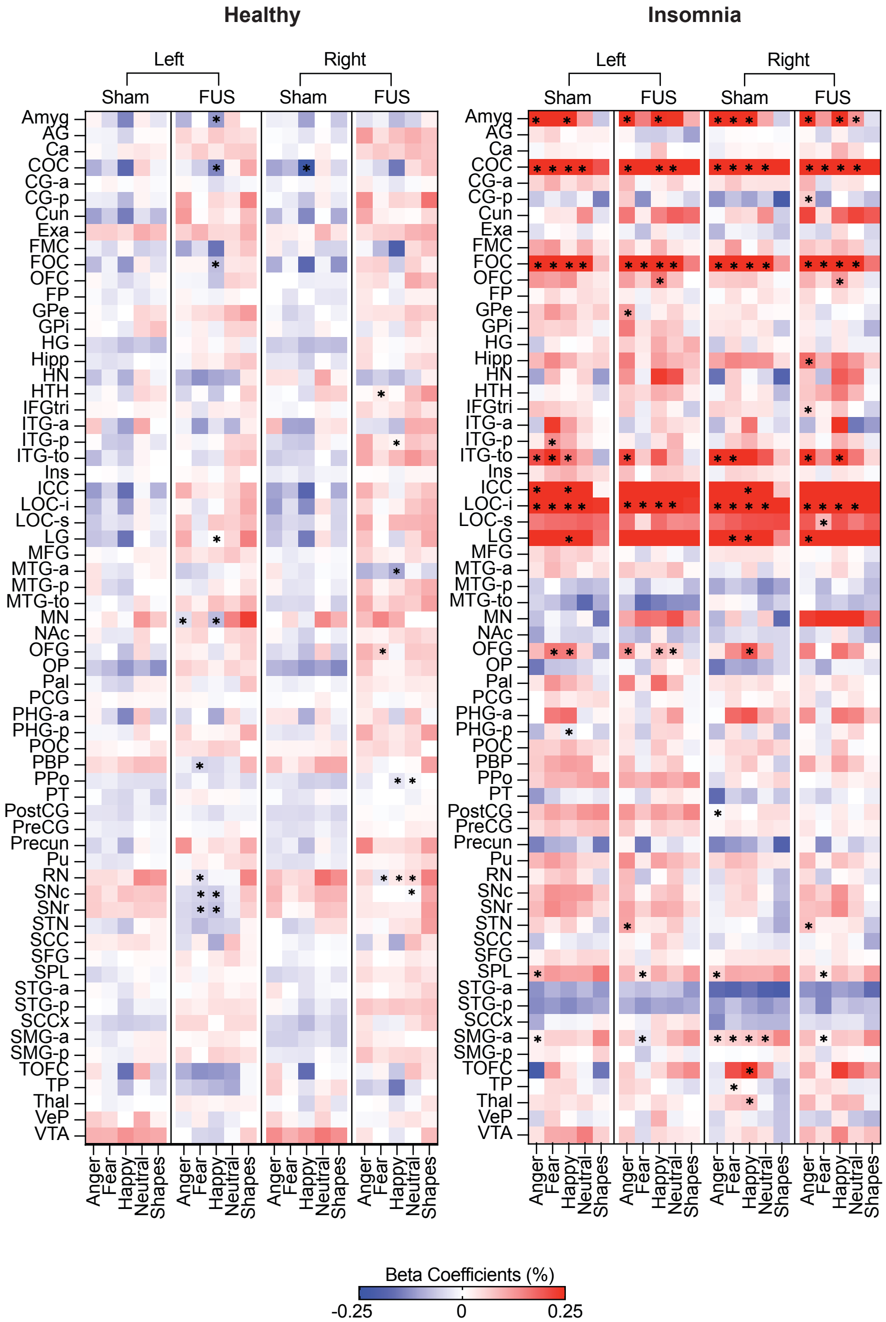

**Supplementary Figure 9. Task-based comparison of beta coefficients between faces and shapes for emotional salient recognition.** Summary of significant different beta coefficients in brain regions across healthy and insomnia patients under varying conditions (FUS, Sham) x (Anger, Fear, Happy, Neutral, Shapes) x (Healthy, Insomnia) with FUS compared to Sham (n = 16 and 12, Two-way ANOVA and Bonferonni's multiple comparison correction). Significant difference compared conditions are labelled with *.

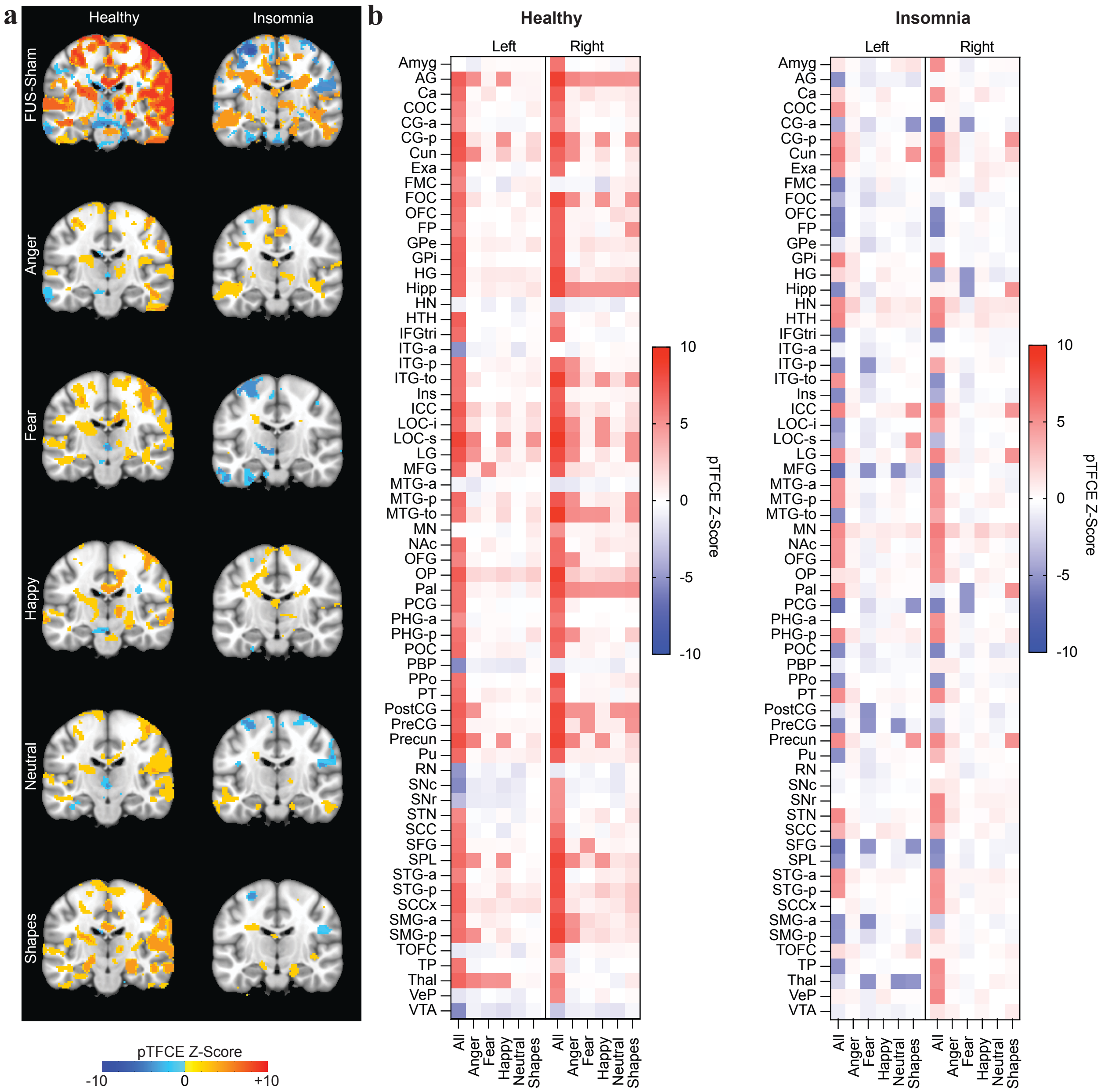

**Supplementary Figure 10. Voxelwise Group-Level Analysis with probabilistic threshold free cluster enhancement of Hariri Task. a)** Sagittal image of voxelwise group-level analysis with pTFCE under conditions in healthy and insomnia groups, thresholded at pTFCE Z-score = 4.90. **b)** Summary of effects under FUS vs. Sham for pTFCE Z-scores in brain regions across healthy and insomnia patients under varying conditions (Anger, Fear, Happy, Neutral, Shapes). pTFCE values were zeroed between -4.90 and 4.90 for statistical representation.

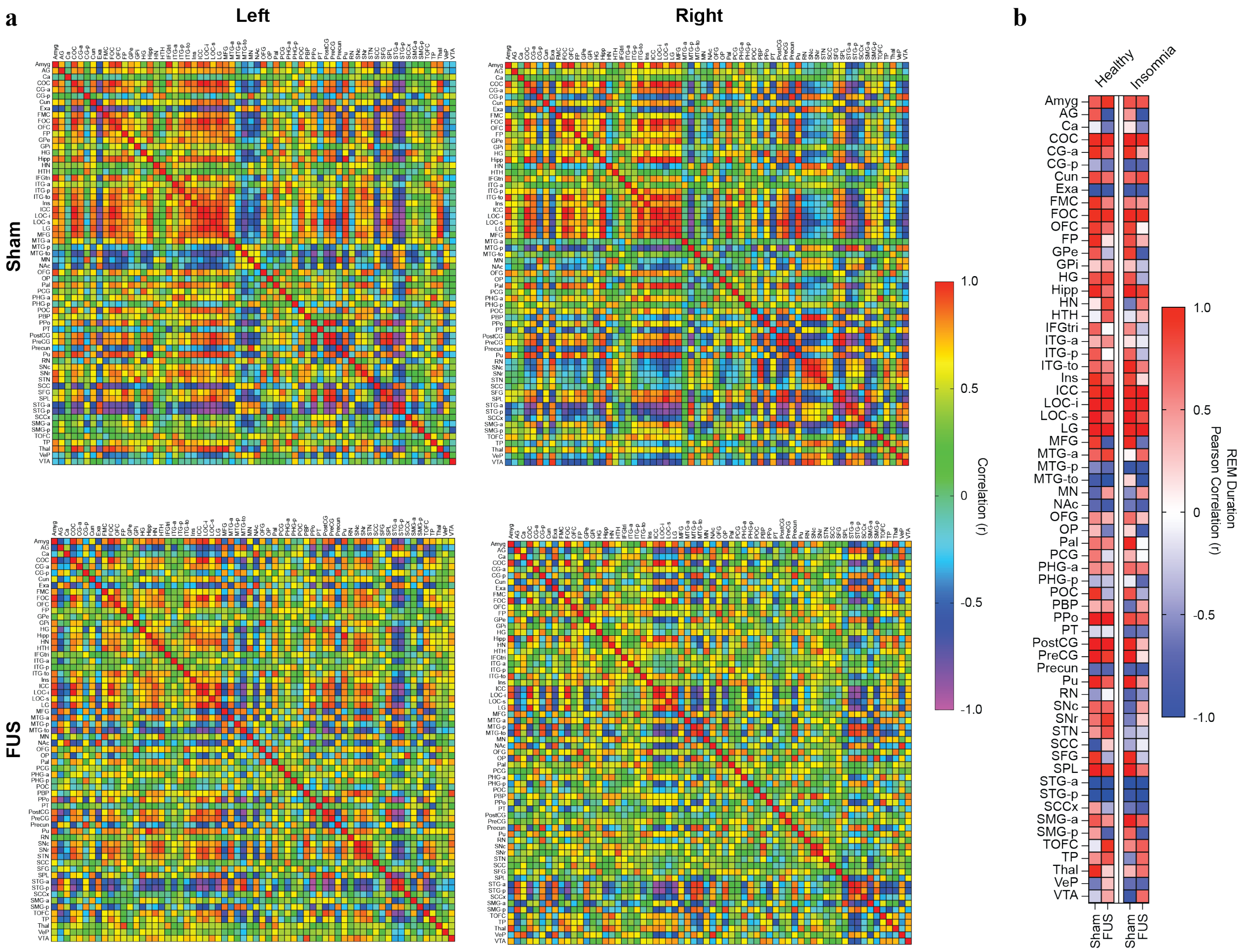

**Supplementary Figure 11. Correlation analysis in effects of STN-FUS. a)** Correlation matrices of left and right ipsilateral ROI in response to Sham vs FUS conditions of collective beta coefficients from all conditions during Hariri Tasks in all subjects (healthy + insomnia). **b)** Pearson's R correlation of beta coefficients with REM duration (%).

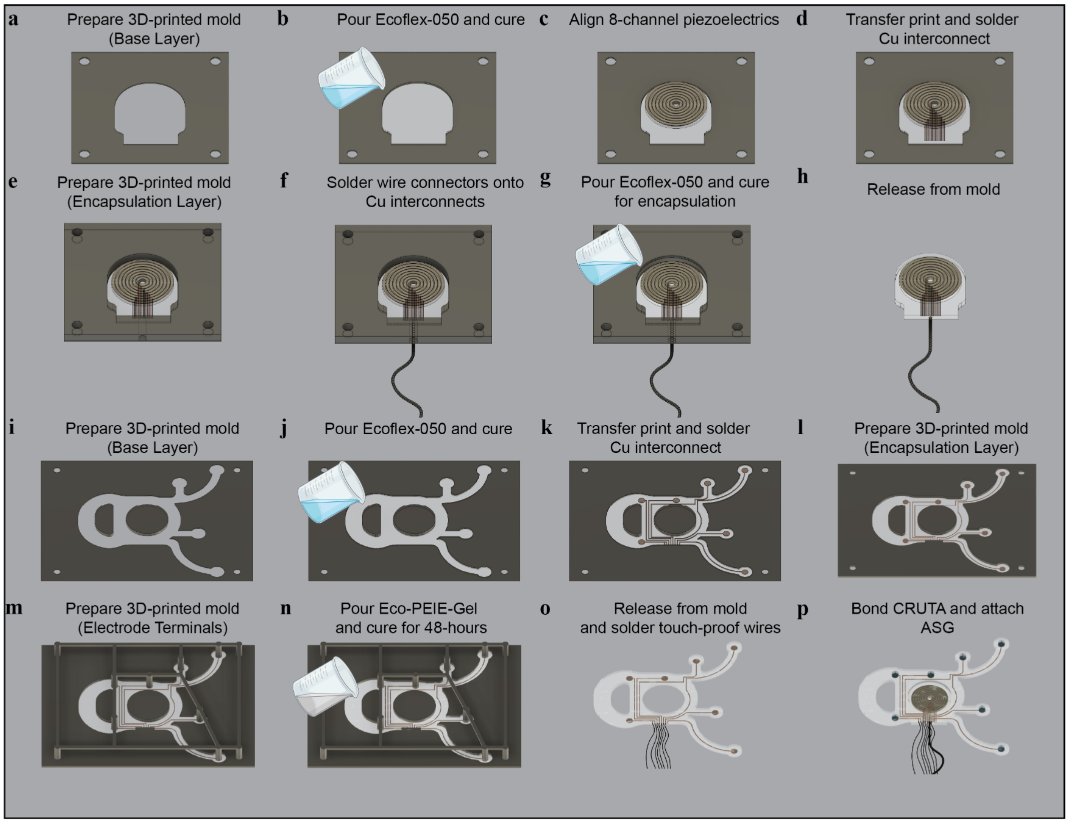

**Supplementary Figure 12. NEUSLeeP fabrication process. a)** Preparation of base layer for CRUTA with 3D-printed PLA mold. **b)** Curing of 1-mm thick backing layer with Ecoflex-050. **c)** Placement of CRUTA by concentrically aligning eight piezoelectric elements. **d)** Transfer printing of interconnect prepared for CRUTA and carefully placed/soldered on to the elements. **e)** Preparation of encapsulation layer for CRUTA with 3.5-mm thick 3D-printed PLA mold. **f)** AWG 36 wires threaded through 0.5" silicon tube and soldered onto interconnects. **g)** Encapsulation of CRUTA with Ecoflex-050 for 60 min. **h)** Release CRUTA from mold. **i)** Preparation of base layer for EEG layer with 1-mm thick 3D-printed PLA mold. **j)** Curing of 1-mm thick backing layer with Ecoflex-050. **k)** Transfer printing of interconnect prepared for EEG layer. **l)** Preparation of encapsulation layer of EEG layer with 3.5-mm thick 3D-printed PLA mold. **m)** Placement of a negative mold for electrode opening for ASG attachment. **n)** Prepare 5% wt PEIE mixed with Ecoflex-Gel (Eco-PEIE-Gel) for encapsulation and cure for 48 hours. **o)** Rinse gently with water to remove excess uncured materials, solder touch-proof wires, and release from mold. **p)** Bond CRUTA and EEG layer with Silpoxy and attach ASG to electrode terminals.

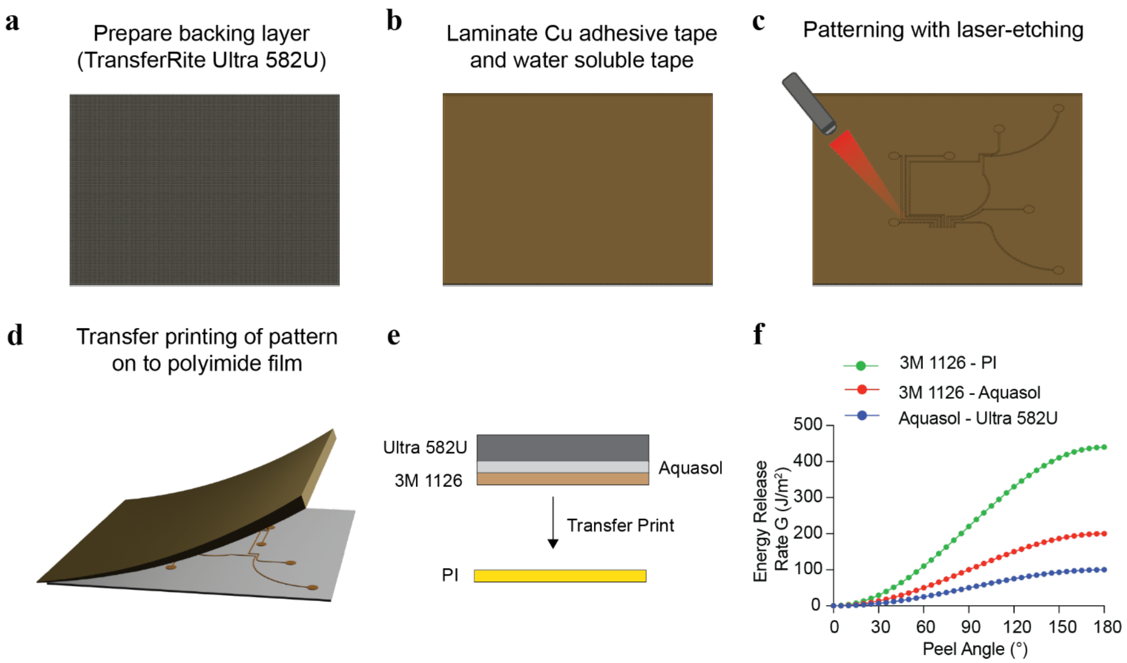

**Supplementary Figure 13. Transfer printing of interconnects. a)** Preparation of backing layer substrate using Ultra 582U adhesive for transfer printing. **b)** Lamination of copper adhesive interfaced with water soluble tape on to backing layer substrate for delamination post-transfer printing. **c)** Laser etched patterning of interconnect using LPKF U4 Protolaser. **d)** Lamination on to polyimide film and peeled at greater than 120° angle for transfer printing and release. **e)** Cross-section view of substrate layers for transfer printing. **f)** Estimated Griffith's energy release rates between substrate materials at various peel angles.

This supplementary analysis evaluates the energy release rate (G) as a function of peel angle for multiple substrate–adhesive pairs. The calculation is based on a simplified form of Kendall's peel model7:

$$G_{energy\ release\ rate} = \frac{F_{peel\ force}}{w_{sample}}\left(1 - \cos\ \theta_{peel}\right) \quad (1)$$

where $F_{peel\ force}$ is the interfacial adhesion strength (N/m) and $\theta_{peel}$ is the applied peel angle. This formulation captures the geometric contribution to crack driving force during peeling, reflecting the balance between external work and the creation of new interfacial surface area.

Four representative systems were modeled: 1) polyamide with 3M 1126 adhesive, 2) TransferRite Ultra 582U with 3M 1126, Aquasol with 3M 1126, and 3) Aquasol bonded to TransferRite Ultra 582U. The results reveal several mechanistic insights. First, energy release rate increases monotonically with peel angle. At low angles (≤30°), G is minimal since most of the applied force is dissipated in shear rather than in driving crack opening. As the angle increases (45–120°), G rises steeply, reflecting more efficient transfer of work into interfacial fracture. At 180°, G is maximized, as the entire applied force contributes to separating the interface. Substrate-dependent adhesion differences are preserved across all angles. Polyamide exhibits the highest values of G, consistent with its stronger interfacial bonding, while the Aquasol–TransferRite interface consistently shows the weakest values, indicating its susceptibility to delamination. These differences highlight the influence of substrate chemistry and interfacial compatibility on adhesive performance. The results underscore how peel geometry amplifies or suppresses apparent adhesion performance. Even strongly bonded interfaces may appear weak at high peel angles because geometry favors crack propagation, while weaker systems may remain stable in low-angle or shear-dominated regimes. This dependence emphasizes that peel strength is not solely a material constant but also a geometry-sensitive property. By framing peel energetics within this broader fracture mechanics context, the present analysis provides a bridge between theoretical crack growth criteria and applied adhesion testing. These calculations, though based on estimated adhesion values, illustrate how energy balance concepts govern detachment across different interfaces and offer mechanistic support for the experimental peel results described in the main manuscript. As such, leveraging the difference in energy release rates between materials, successful transfer printing was achievable.

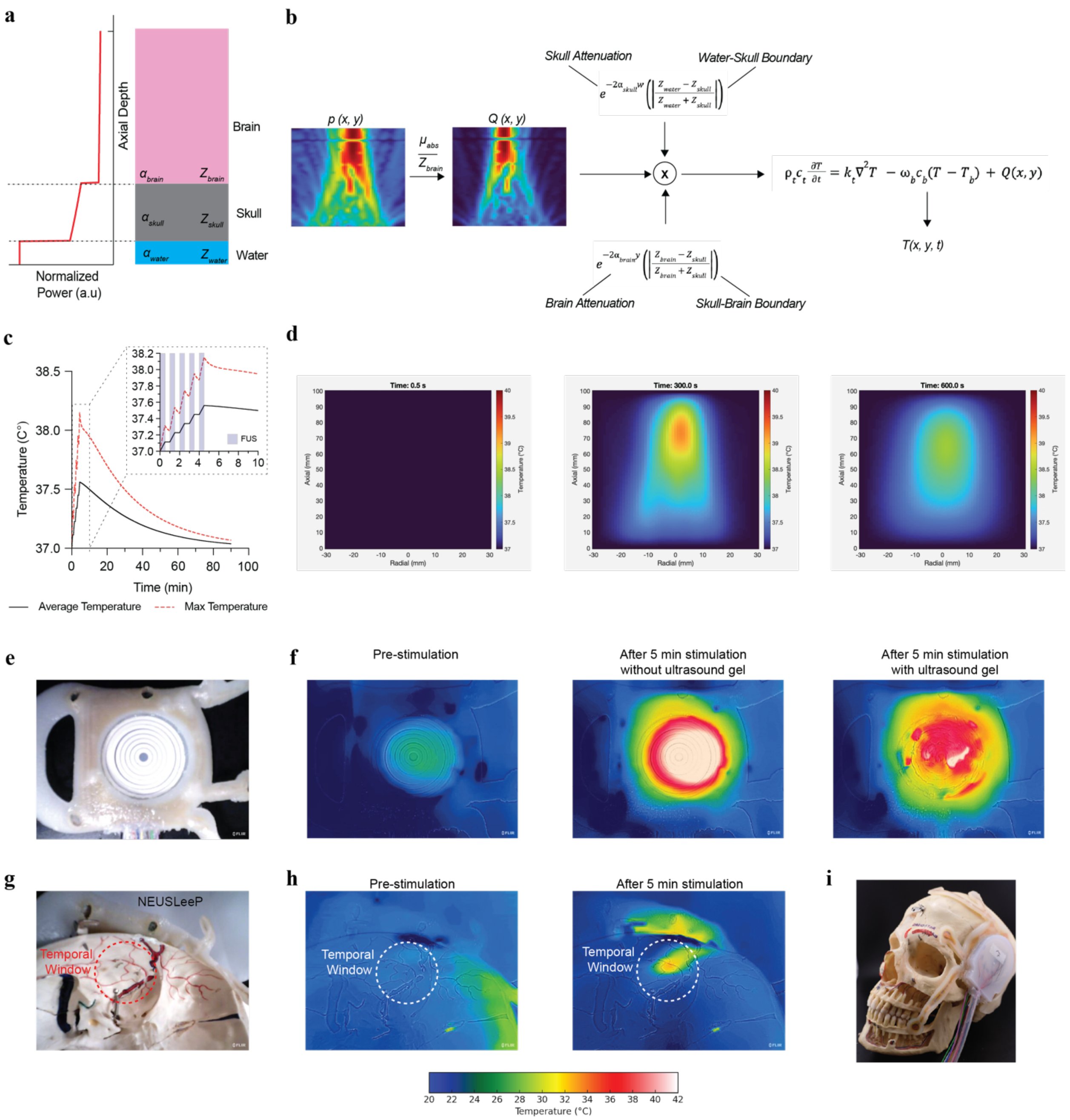

**Supplementary Figure 14. Thermal heating effects of CRUTA. a)** Schematic representation of acoustic power loss. **b)** Computational method for thermal diffusion and estimation of *in-vivo* thermal heating effects of CRUTA for STN-FUS. **c)** Calculated thermal heating response with CRUTA for STN-FUS at the desired focal depth. **d)** Simulated thermal distribution at various time points (t = 0.5s, 300s, 600s). **e)** Photograph of CRUTA in NEUSLeeP. **f)** Surface temperature of CRUTA before and after stimulation with/without ultrasound gel. **g)** Photograph of endocranium temporal window and placement of NEUSLeeP at the endocranium temporal window. **h)** Surface temperature of endocranium temporal window from CRUTA before and after stimulation. **i)** Photograph of NEUSLeeP on human skull.

A 2-D transient bioheat model was used to estimate temperature rises produced by a pulsed transcranial ultrasound field. Implementation of a finite-difference, implicit (backward-Euler) time integrator for numerical stability with inclusion of perfusion cooling and frequency-dependent acoustic attenuation was used to effectively emulate the experimental conditions as close as possible. The temperature T(x,t) evolves under a Pennes-type bioheat equation,

$$\rho_t c_t \frac{\partial T}{\partial t} = k_t \nabla^2 T - \omega_b c_b (T - T_b) + Q(x,y) \quad (2)$$

with tissue properties $c_t$=3770 J $kg^{-1}K^{-1}$, $k_t$=0.195 W $m^{-1}K^{-1}$, $\rho_t$=1040 kg $m^{-3}$. The arterial temperature is fixed at $T_a$=37CTa=37$\circ$C. Perfusion is computed from a typical gray-matter rate (80 mL/min/100 g) and converted to $\omega_b$ with density correction (Supplementary Table 6.).

$$\omega_b = \frac{\rho_b * CBF_{gray}}{\rho_t} \quad (3)$$

Next, acoustic–thermal coupling was determined by evaluating the volumetric source:

$$Q(x,y) = \mu_{abs} \frac{|p_{free-field}(x,y)|^2}{Z_{brain}} e^{-2\alpha_{skull} w} \left( \left| \frac{Z_{water} - Z_{skull}}{Z_{water} + Z_{skull}} \right| \right) e^{-2\alpha_{brain} y} \left( \left| \frac{Z_{brain} - Z_{skull}}{Z_{brain} + Z_{skull}} \right| \right) \quad (4)$$

where $P_{free\text{-}field}$ is the measured free-field acoustic pressure (scaled to Pa), Z as the acoustic impedance of the medium, and $\mu_{abs}$ the Napierian absorption coefficient derived from attenuation in dB using 1 dB=0.1151. Axial decay within tissue is estimated via $e^{(-2\mu_{abs} y)}$ Additional skull transmission loss is modeled by a scalar amplitude factor computed from α (20 dB $cm^{-1}$ $MHz^{-1}$), frequency (0.65 MHz), and skull thickness (0.6 cm). With temporal protocol, pulse gating of Q with a PRF of 100 Hz and 5% duty cycle (periodic on-time inside each 10 ms PRF window) and with 30 s ON / 30 s OFF blocks was applied. This preserves intra-PRF pulsing and block-level stimulation.

### References

1. Forkert, N. D., Li, M. D., Lober, R. M. & Yeom, K. W. Gray Matter Growth Is Accompanied by Increasing Blood Flow and Decreasing Apparent Diffusion Coefficient during Childhood. *AJNR Am J Neuroradiol* **37**, 1738–1744 (2016).
2. Sepehrband, F. *et al.* Brain tissue compartment density estimated using diffusion-weighted MRI yields tissue parameters consistent with histology. *Hum Brain Mapp* **36**, 3687–3702 (2015).
3. Trudnowski, R. J. & Rico, R. C. Specific Gravity of Blood and Plasma at 4 and 37 °C. *Clin Chem* **20**, 615–616 (1974).
4. Fjield, T., Fan, X. & Hynynen, K. A parametric study of the concentric-ring transducer design for MRI guided ultrasound surgery. *J. Acoust. Soc. Am.* **100**, 1220–1230 (1996).
5. Ko, S.-B. *et al.* Real time estimation of brain water content in comatose patients. *Ann Neurol* **72**, 344–350 (2012).
6. Gupta, S., Haiat, G., Laporte, C. & Belanger, P. Effect of the acoustic impedance mismatch at the bone-soft tissue interface as a function of frequency in transcranial ultrasound: A simulation and in vitro experimental study. *IEEE Trans. Ultrason. Ferroelectr. Freq. Control* **68**, 1653–1663 (2021).
7. Kendall, K. Thin-film peeling-the elastic term. *J. Phys. D Appl. Phys.* **8**, 1449–1452 (1975).
